# Site-resolved spatial and structural interactome of a human cell

**DOI:** 10.64898/2026.09.08.749775

**Authors:** Zehong Zhang, Nanako Yokoyama, Ying Zhu, Julia Ruta, Jiale Cheng, Vini Natalia, Tolga Soykan, Volker Haucke, Fan Liu

## Abstract

The spatial and structural arrangement of proteins determine virtually every process in human cells. We combined gentle subcellular fractionation by differential ultracentrifugation with cross-linking mass spectrometry to systematically map this cellular proteome architecture with residue-level evidence, identifying 164,146 residue-to-residue links in HEK293 cells. These links capture spatial protein arrangement at a resolution sufficient to pinpoint protein localizations at sub-organelle level, determine protein orientations within cellular membranes, and identify inter-organelle contact sites. The residue-level information provides evidence for 18,074 direct protein-protein interactions (PPIs), which we integrate into AlphaFold-based pipelines to nominate PPI-mediating short linear motifs and generate assembly models of large protein complexes. Guided by these spatial and structural readouts, we discover new PPIs within the endomembrane system that regulate the compartmental localization of trafficking machinery. Leveraging a network topology-driven strategy, we augment our HEK293 dataset with PPI data from different cell lines and methods, expanding the spatial and structural interactome of human cells.

## Introduction

Protein–protein interactions (PPIs) are fundamental to protein function and cell physiology. Proteins rarely act in isolation; instead, they form dynamic, spatially organized networks. Substantial effort has therefore been directed toward comprehensive PPI databases such as BioPlex (*1*) and OpenCell (*2, 3*), built primarily through affinity purification coupled with mass spectrometry (AP-MS). Complementary cellular interactomics efforts have exploited proximity-labeling techniques, spatial proteomics strategies that resolve subcellular localization through biochemical fractionation, fluorescence-based imaging, or combinations thereof (*3–7*). These approaches, however, cannot reliably distinguish direct from indirect interactions, provide no structural information on the interaction interfaces, and are limited in their ability to elucidate the orientation of proteins in cellular membranes. Furthermore, many of them detect PPIs from genetically engineered cell lines or tagged proteins, which might impact the interaction pattern.

Cross-linking mass spectrometry (XL-MS) provides an opportunity to fill these blind spots. XL-MS identifies maximum residue-to-residue distances across the proteome, thereby revealing PPIs as well as the corresponding protein binding interfaces, subcellular localizations, and membrane topologies (*8–10*). XL-MS has been applied for generating PPI data on cellular scale (*11, 12*), but these datasets are too incomplete to meaningfully complement resources like BioPlex and OpenCell, and systems-level analysis strategies to leverage all aspects of XL-MS information are lacking.

Here, we generate a deep structural interactome of a human cell by combining proteome-wide XL-MS with differential ultracentrifugation (DUC). Our DUC-XL-MS interactome covers 9-times more PPIs than the largest published XL-MS datasets. We exploit this dataset from the perspectives of cell, structural and systems biology. On the cellular level, we map organelle-specific interactomes, resolve membrane protein topologies, identify inter-organelle nexus proteins, and elucidate how the immunodeficiency-associated protein LRBA regulates protein compartmental localization. On the structural level, the DUC-XL-MS data guided proteome-scale screening to validate and discover PPI-mediating short linear motifs (SLiMs) as well as assembly modeling of endogenous protein machineries. For the latter, we modeled several large protein complexes by leveraging cross-linking-restrained AF3x predictions (*13*) and Monte Carlo-based hierarchical assembly. On the systems level, comparing DUC-XL-MS to published PPI resources shows that each dataset captures a distinct interactome niche. A similar level of complementarity arises between different human cell lines, as we demonstrate on cell line-specific BioPlex 3.0 data and by acquiring a second DUC-XL-MS dataset in Jurkat cells. We leverage this complementarity by developing a network topology-driven approach that generates multi-resource augmented interactomes. After validating this augmentation approach against PPIs from the Protein Data Bank (PDB), we use it to define a spatially and structurally supported human interactome of 28,906 PPIs.

## Results

### A human cell interactome that faithfully recapitulates subcellular spatial organization

To build a comprehensive XL-MS-based cellular interactome, we combined cross-linking mass spectrometry (XL-MS) with gentle cell lysis and differential ultracentrifugation (DUC)(*4*) to deepen cross-link coverage without fully disrupting the cellular context (DUC-XL-MS, Fig. S1). We cross-linked cleared lysates from immortalized non-cancer human embryonic kidney (HEK) 293 cells with the enrichable, MS-cleavable cross-linker Azide-A-DSBSO and fractionated it by DUC. The fractions recapitulated the organelle marker distributions previously reported for DUC of non-crosslinked fractions(*4*) (Fig. S2). At a 1% FDR at the residue-pair level, DUC-XL-MS yielded 164,146 total residue pairs (75,145 inter-protein residue pairs), corresponding to 18,074 inter-protein PPIs across 5,399 proteins (**Fig. 1A**, Table S1). This represents an approximately 9-fold increase in inter-protein PPI coverage over the largest published XL-MS interactomes from intact ((*14*); 9.4-fold) and fractionated ((*11*); 8.6-fold) human cell samples, and around 5-fold gain of residue-to-residue distance information (**Fig. 1B**).

**Figure 1.**
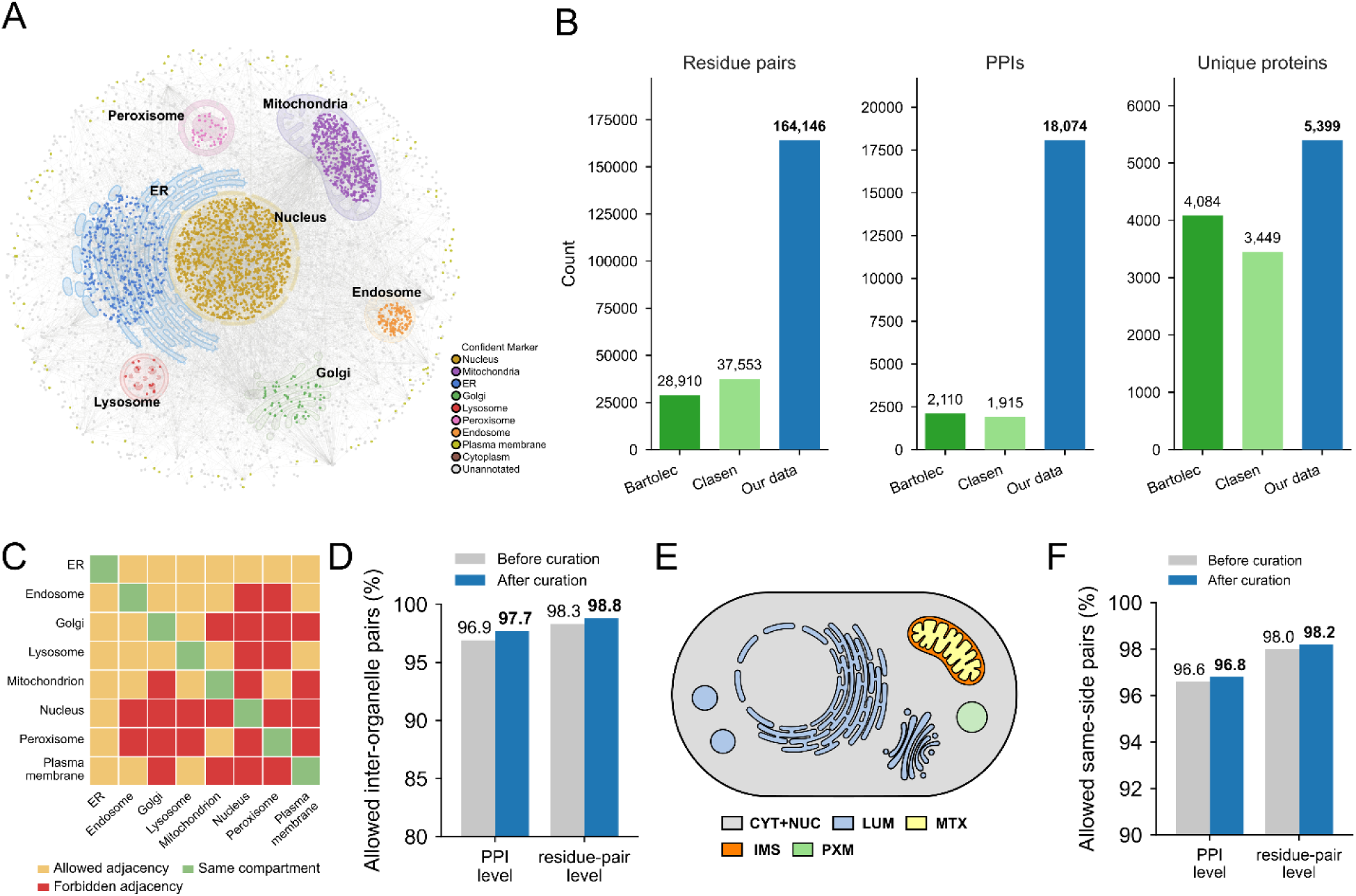
An XL-MS-derived cellular interactome and its internal consistency. (A) XL-MS-based cellular interactome of HEK293 cells. Nodes are proteins and edges are inter-protein cross-links. Nodes are colored by organelle localization, and gray nodes are unannotated. (B) Coverage comparison of the number of residue pairs, PPIs and cross-linked proteins in this study versus previously published human in-cell XL-MS datasets. (*11, 14*) (C) Organelle adjacency map defining which compartment pairs are allowed or not allowed to be cross-linked. Orange: allowed adjacency; red: forbidden adjacency; green: same compartment. (D) Organelle-level spatial consistency of the XL-MS network: the percentage of marker-marker cross-links that are spatially plausible (connecting the same or an adjacent compartment, as defined in C), before and after manual correction. (E) Side-resolved consistency map defining same-side (allowed) versus cross-side (forbidden) cross-links. Five sides are defined, each with a distinct color: CYT+NUC (cytoplasm + nucleoplasm), IMS (mitochondrial intermembrane space), MTX (mitochondrial matrix), LUM (secretory/luminal + extracellular space) and PXM (peroxisomal matrix). (F) Side-resolved spatial consistency of the XL-MS network: the percentage of marker-marker cross-links that are spatially plausible (connecting residues on the same side, as defined in E), before and after manual correction.

We tested whether the cross-links respect the cell’s spatial organization using two complementary consistency maps:

The first is organelle adjacency: proteins from two compartments may be cross-linked only if the inter-compartment contact is physiologically feasible (**Fig. 1C**). For this analysis, we selected a reference set of 2,011 organelle markers: 1,833 single-organelle and 178 dual-organelle localized proteins (Table S2, Methods - Organelle marker assignment). We classified each marker-to-marker cross-link as spatially possible or impossible according to whether the two partners’ localizations permit a direct interaction (**Fig. 1C**). Across these markers, 98.3% of all cross-linked residue pairs (96.9% of PPIs) were spatially consistent (**Fig. 1D**, Table S2).

The second is membrane separation: because the cross-linker’s spacer arm is shorter than the diameter of cellular membranes, the two cross-linked lysines must lie on the same side of the membrane. We used UniProt subcellular annotations to select cross-linked soluble proteins assigned to a single membrane-enclosed locale (**Fig. 1E**), yielding 3,587 topology markers (Table S2, Methods - Topology marker assignment). We classified each marker-to-marker cross-link as topologically possible only when both residues occupy the same side. Agreement among the topology markers is 98.0% at the residue-pair level (96.6% at the PPI level) (**Fig. 1F**, Table S2).

Since both marker sets are part of an interconnected network, they can be additionally validated against their cross-linked neighbors. At both the organelle and topology levels, DUC-XL-MS gave a consistency rate very similar (<1% difference) to that of our previously published XL-MS datasets of intact HEK293 cells (*14*), confirming that DUC fractionation did not disrupt protein arrangement in the tested compartments (Supplementary Note 1). Furthermore, the DUC-XL-MS consistency rate was 15–60% higher than a randomized null in which cross-links connect random protein pairs (Supplementary Note 1). Beyond assessing dataset quality, this spatial consistency validation also flags suspicious UniProt annotations for manual curation (summarized in Table S2). Applying these corrections increased the agreement rate within the organelle and topology marker sets by 0.2–0.8% (blue bars in **Fig. 1D,F**) and yielded a final reference of 2,001 high-confidence organelle markers and 3,587 high-confidence topology markers.

### DUC-XL-MS residue-level resolution reveals protein localization, membrane topology, and organelle contacts

Having validated the network, we used the organelle and topology markers as seeds for spatial annotation, working from whole-compartment localization to the topology of individual membrane proteins and the contacts between compartments.

We first propagated the organelle markers one step to the remaining non-marker proteins using a graph-based approach with defined rules (**Fig. 2A**, Methods: Graph-based propagation of protein localization). Starting from the 2,001 high-confidence organelle markers, propagation expanded the spatial annotation to 2,639 proteins across all eight compartments (**Fig. 2B**, Table S3). These annotations closely recapitulated the spatial organization resolved by LOPIT-DC in principal component space (Fig. S3)(*4*) and reproduced the organelle topology of the OpenCell image-based UMAP embedding (Fig. S4)(*3*), confirming their agreement with independent spatial proteome resources.

**Figure 2.**
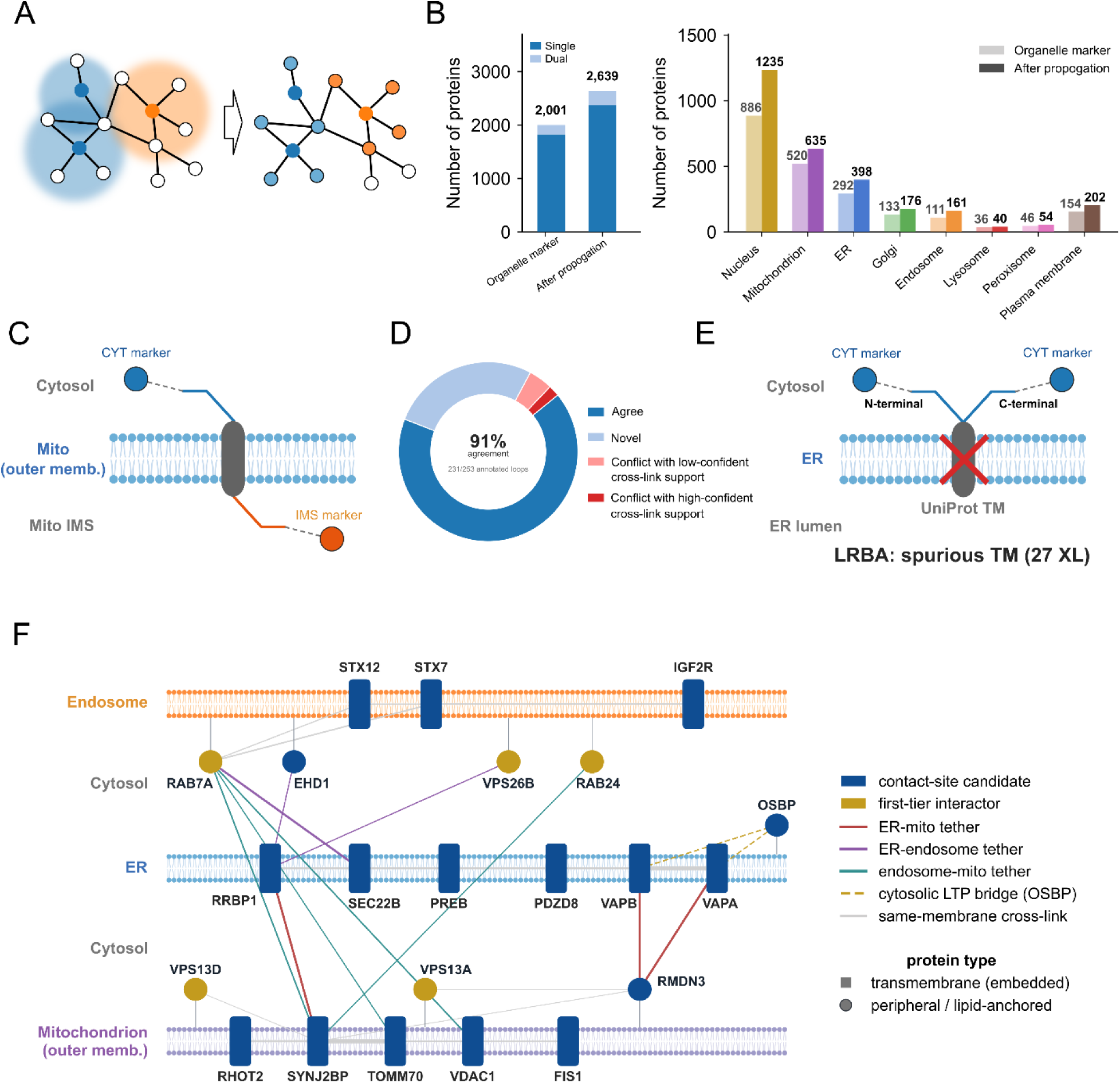
XL-MS reveals protein localization, membrane topology, and organelle contacts. (A) Schematic of graph-based annotation propagation. Starting from marker proteins (colored), organelle labels are propagated across cross-link edges to unannotated neighbors. (B) Number of spatially annotated proteins before and after propagation, shown as an overall total and distributed across the eight cellular compartments. (C) Schematic diagram of the topology assignment of transmembrane proteins. (D) Transmembrane-loop topology assignment from XL-MS and its comparison with the UniProt annotation. Categories are color-coded: agree (green); conflict, comprising high-confidence conflicts that are credibly correctable (dark red) and those with only low-confidence cross-link support (pink); and novel, i.e. not annotated in UniProt (pale blue). (E) Cross-link correction of a spurious transmembrane annotation. UniProt annotates LRBA as a single-pass transmembrane protein, but cross-links place both its N-terminus and C-terminus in the cytosol. Because a genuine single-pass protein cannot have both termini on the same side of the membrane, the transmembrane annotation is spurious (red cross) and LRBA is a soluble BEACH-domain protein. (F) Cross-link network of ER–mitochondria–endosome contact sites. Nodes are drawn by topology: transmembrane proteins (squares) embedded in the endosome (top), ER (middle) or outer mitochondrial membrane (bottom); peripheral or lipid-anchored proteins (circles) on the cytosolic face; and soluble ER-luminal proteins (open circles). Blue nodes are contact-site candidates (dual-localized or resolved-topology transmembrane proteins); gold nodes are their first-tier cross-link interactors. Edge color marks the tether type (red, ER–mitochondria; purple, ER–endosome; teal, endosome–mitochondria; dashed gold, cytosolic OSBP (oxysterol-binding protein) –VAP (VAMP-associated protein) lipid-transfer bridge; gray, same-membrane), and edge thickness scales with the number of supporting residue pairs. The late-endosomal GTPase RAB7A (dashed ring) bridges all three membranes from the cytosolic face.

At finer resolution, residue-level connectivity predicts which side of the membrane each transmembrane-protein loop faces (**Fig. 2C**), as we have shown in mitochondria (*9*). Using the 3,587 topology markers, we assigned membrane sides to 346 loops across 304 transmembrane proteins. Of these, 231 loops (205 proteins) agree with the existing UniProt topology, 93 (82) are new assignments, and 22 (20) conflict with prior annotation (**Fig. 2D**). Conflicts with strong cross-link support can credibly flag existing topology annotations for correction (Table S3). For example, kinectin’s C-terminal domain is annotated as luminal, yet DUC-XL-MS data place it in the cytosol as expected for the kinesin receptor (Fig. S5).

For proteins with at least two assigned loops, the loops must fit a single alternating topology, because each conventional transmembrane helix places its flanking loops on opposite sides of the membrane. This internal consistency check validates 31 of 40 testable proteins. As an agreeing example, the 13-transmembrane ER oligosaccharyltransferase subunit STT3B is resolved by three mutually consistent loops (Fig. S6). Conversely, the Lipopolysaccharide-responsive and beige-like anchor protein (LRBA) has a single transmembrane segment according to UniProt, yet our cross-links place both flanking residues on the cytosolic side (**Fig. 2E**), thus revealing a spurious transmembrane annotation and indicating that LRBA, instead, associates peripherally with organelle membranes. As we show below, this reclassification is consistent with a role for LRBA as a soluble trafficking regulator.

Next, we mined the DUC-XL-MS data for inter-organelle contact sites, focusing on dual-organelle localized proteins and transmembrane proteins with a resolved membrane topology. Overlapping these candidates with the trafficking complexes from CORUM (*15*) and membrane contact site (MCS) databases (*16*) recovers known machineries (Fig. S7, Table S3). For instance, at the ER–mitochondria interface, the cross-links recapitulate the RRBP1–SYNJ2BP (*17*) and VAPB–RMDN3 (PTPIP51) (*18*) tether pairs, and we additionally detect the ER-anchored OSBP–VAP lipid-transfer bridge (*19*) (**Fig. 2F**). Beyond these pairwise contacts, the late-endosomal GTPase RAB7A cross-links to the outer mitochondrial membrane (SYNJ2BP, TOMM70, VDAC1), to endosomal partners (STX7, STX12), and to the ER (SEC22B), consistent with the tri-membrane ER–endosome–mitochondria contacts described (*20*) (**Fig. 2F**). The RAB7A–SEC22B cross-link (Fig. S8) places SEC22B at ER–late endosome contacts, which to our knowledge has not been reported previously.

Finally, we focused on the multi-localized protein LRBA, which our topology assignment reclassified as peripherally membrane-associated and which based on our DUC-XL-MS data, localized to endosomes and the Golgi (**Fig. 3A**). LRBA functions in trafficking of the immune-checkpoint receptor cytotoxic T-lymphocyte-associated protein 4 (CTLA-4)(*21, 22*). Extending this role, LRBA’s cross-linking data indicate a previously unreported interaction with another trafficking regulator: the endosomal SNARE protein STX7 (**Fig. 3A**). LRBA and STX7 co-localize in the perinuclear area of HeLa cells, likely corresponding to endosomes and the trans-Golgi network (TGN) (**Fig. 3B**). LRBA-knockout (KO) cells showed significantly reduced intensity of STX7 puncta within the TGN/endosome area compared to wild-type (WT) controls (**Fig. 3C**), while the overall STX7 protein levels remained constant (**Fig. 3D**, Fig. S9). We therefore hypothesized that LRBA controls the subcellular localization of STX7 rather than its abundance. To test this, we performed confocal imaging of STX7 alongside the early endosomal marker EEA1, the TGN marker TGN46, the cis-Golgi marker GM130, the late endosomal/lysosomal marker LAMP1, and the ER marker calreticulin (CALR). These experiments revealed reduced steady-state levels of STX7 in the TGN and Golgi area, at early endosomes and at late endosomes/lysosomes in LRBA KO cells, whereas ER levels of STX7 were elevated (**Fig. 3E–I**, Fig. S10). Hence, loss of LRBA causes a specific redistribution of STX7 away from endosomal/TGN compartments toward the ER, suggesting that LRBA physically and functionally chaperones STX7 trafficking and localization.

**Figure 3.**
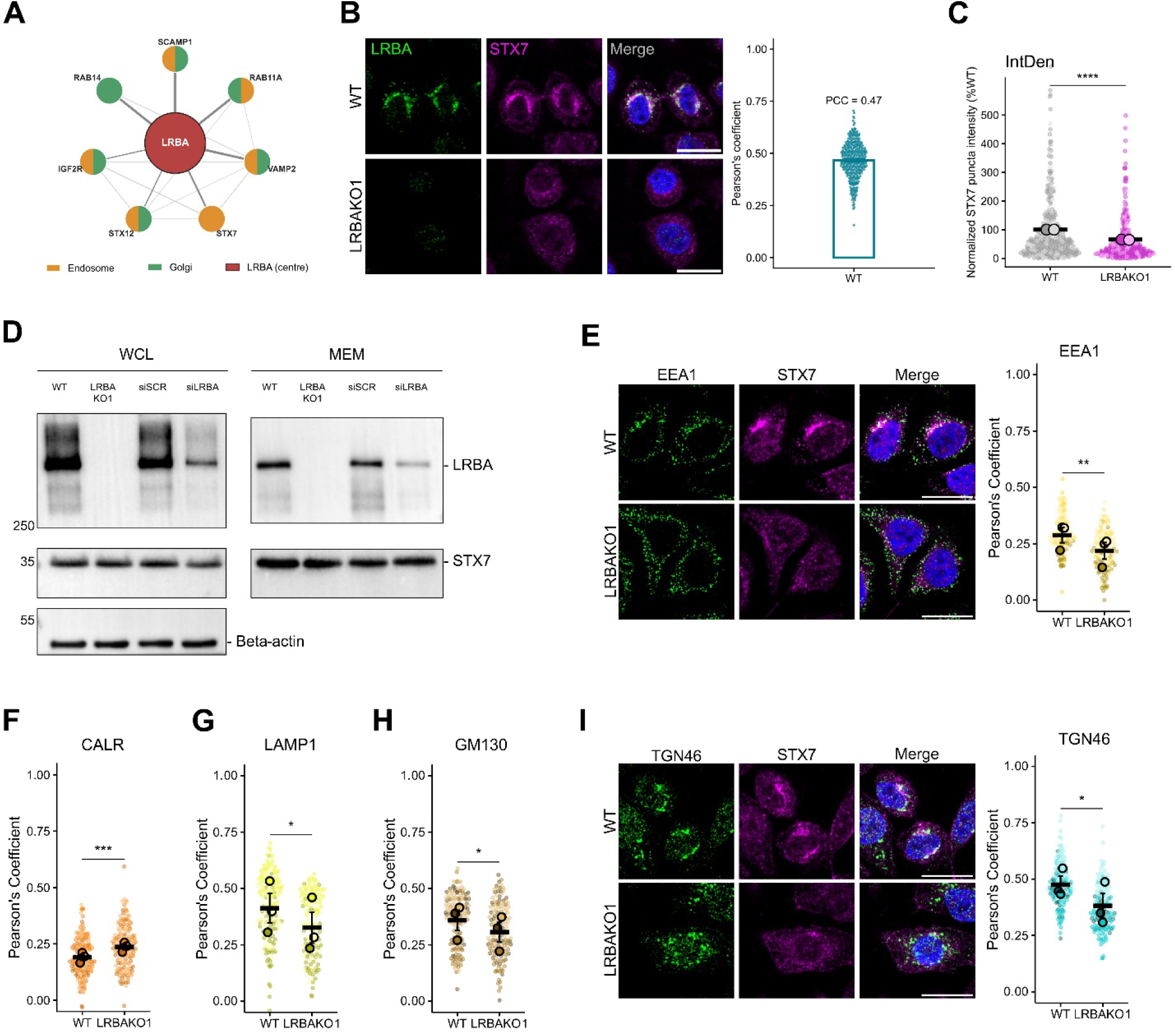
LRBA is an endosomal SNARE chaperone required for STX7 localization. (A) LRBA direct interactors with confident localization. LRBA’s direct cross-link partners, each colored by its organelle assignment. Split nodes denote dual-localized proteins and edge width scales with residue-pair count. (B) Confocal microscopy of HeLa WT and LRBA□ KO cells co□stained with anti□LRBA and anti□STX7. Colocalization of LRBA and STX7 in HeLa WT cells was quantified by Pearson’s correlation coefficient (PCC; n = 340 cells from two independent experiments). Scale bars: 20 μm. (C) STX7 puncta total intensity per cell, normalized to the WT group (% WT). Small dots represent individual cells and large color□coded dots represent replicate means from two independent experiments. Data are shown as mean ± s.e.m.; the y□axis is truncated at 600 for visualization only, and all data points were included in the statistical analysis. Statistics: two□tailed Mann–Whitney test; ****P < 0.0001; n = 336 (WT), n = 312 (KO). (D) Immunoblot analysis of STX7 protein levels upon LRBA knockout and siRNA□mediated LRBA knockdown (WCL = whole□cell lysate, MEM = membrane fraction). (E–I) Confocal microscopy of HeLa WT and LRBA KO cells co-stained with anti-STX7 and markers of endomembrane organelles. Colocalization of STX7 with each marker was quantified as Pearson’s correlation coefficient and shown as superplots: small dots represent individual cells, large color-coded dots the means of N = 3 independent biological replicates. Data are shown as mean ± s.e.m. Statistics: two-sided paired t-test; ***P < 0.001, **P < 0.01, *P < 0.05. Scale bars, 20 μm. Additional representative images are shown in Fig. S10.

### A cross-link-guided AlphaFold screen for short linear motifs

Beyond capturing protein localization and topology, the DUC-XL-MS network readily distinguishes direct from indirect PPIs, which allows prioritizing realistic interactions for structure modeling(*10*). We first focused on short linear motifs (SLiMs) – short amino acid stretches in intrinsically disordered regions that mediate many PPIs (*23*). Recent studies have shown that AlphaFold can identify a SLiM as a locally confident inter-chain contact within an otherwise low-confidence complex (*24, 25*). Therefore, we constructed an AlphaFold3 (*26*)-based pipeline to screen for motif-like behavior among the cross-link-supported direct PPIs (**Fig. 4A**, Methods: Short linear motif (SLiM) screening pipeline).

**Figure 4.**
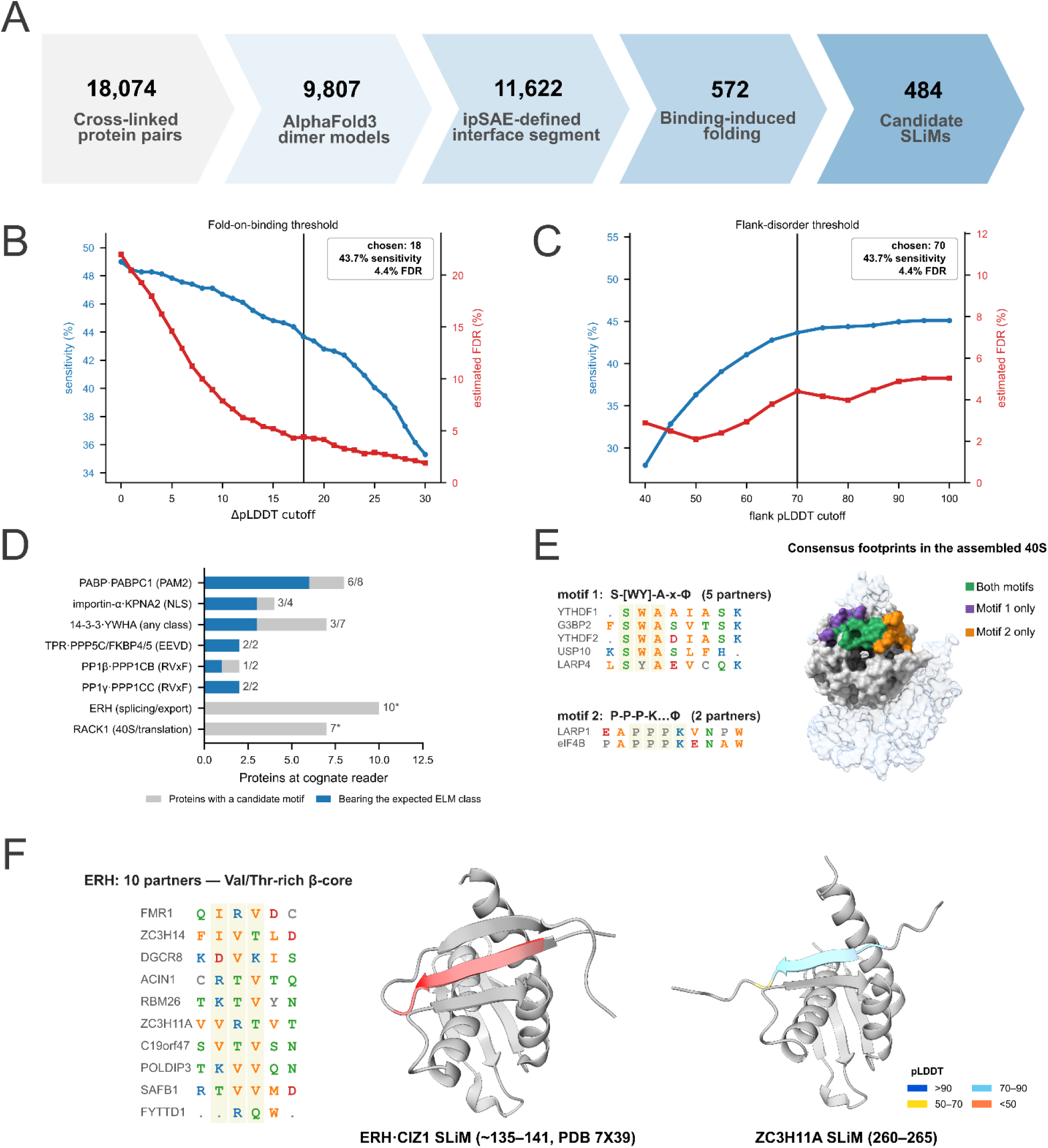
A cross-link-guided AlphaFold screen for short linear motifs (SLiMs). (A) Workflow. 9,807 AlphaFold3 dimer models were passed through the SLiM screening pipeline in four consecutive steps: (1) a confident inter-chain contact, (2) interface scoring and localization with ipSAE, (3) binding-induced folding and (4) a disordered context. The pipeline yields 484 candidate motifs across 301 proteins. (B, C) ΔpLDDT and flanking pLDDT thresholds were assessed against ELM-annotated pairs (positive control) and length- and monomer-pLDDT-matched non-interface windows (negative control). Sensitivity (fraction of the ELM-annotated pairs recovered; blue, left axis) and estimated FDR (negative-control pass rate ÷ interface pass rate; red, right axis) are shown versus the ΔpLDDT threshold (B, with flanking pLDDT < 70 held fixed) and the flanking-pLDDT threshold (C, with ΔpLDDT ≥ +18 held fixed). The selected cutoffs (solid lines; ΔpLDDT ≥ +18 and flanking pLDDT < 70) give 44% sensitivity at 4.4% FDR. (D) Recovery of known SLiM classes at cognate SLiM-binding proteins. Counts are proteins. Pale bars, candidate SLiMs bind to the indicated SLiM-binding protein; saturated bars, the subset whose motif matches that protein’s expected ELM class. ERH and RACK1 are hubs rich in SLiM candidates with no single ELM consensus (panels E, F). (E) The candidate RACK1 motifs. Left: motif 1, an aromatic-anchored S-[W/Y]-A-x-Φ consensus shared by five evolutionarily unrelated partners (YTHDF1, YTHDF2, G3BP2, USP10, LARP4; ≈25% pairwise identity, except between the YTHDF1/YTHDF2 paralogues); and motif 2, a proline-rich P-P-P-K…Φ element in LARP1 and eIF4B. Right: contact areas of the two classes on RACK1 (pale surface) within the assembled 40S (PDB 8GLP; neighboring ribosomal components, blue). RACK1 residues contacted by every partner SLiM (6 Å cutoff) are colored. Green, contacted by both classes (13 residues); purple, motif 1 only (9); orange, motif 2 only (11). The two classes share a central surface on blades 5 and 6 of RACK1 and extend from it in opposite directions: motif 1 toward blade 5, motif 2 toward the N- and C-terminal strands that close the propeller. The combined contact area (all three colors; 33 residues, 1,848 Å²), 99.3% of which is solvent-accessible upon ribosome assembly. (F) The candidate ERH binders. Left: the ten ERH partners share only a loose Val/Thr-rich, β-strand-like core (shaded) with no fixed consensus. Middle: the ERH·CIZ1 crystal structure (PDB 7X39), in which the CIZ1 peptide adds as an antiparallel strand on ERH strand 48–54. Right: a representative model (ERH·ZC3H14) reproduces the same binding site. The CIZ1 β-strand (SLKVTIL) is colored red in the crystal structure, while the modeled peptide in the representative model is colored according to AlphaFold pLDDT. ERH is shown in grey. The binding of all ten ERH partners are shown in Fig. S11.

We randomly sampled 9,807 PPIs from our dataset and predicted their AF3 dimer models, of which 2,005 carried a confidently modelled inter-chain contact. We scored each interface with ipSAE, a residue-specific interface-confidence metric(*27*). We retained confidently predicted segments that both fold on binding (measured as ΔpLDDT between the complex and monomer models(*28*)) and lie in a disordered context (defined by the mean pLDDT of the flanking regions). After optimizing ΔpLDDT and flanking pLDDT (*29*) against bespoke independent positive and negative controls, we obtained 484 candidate motifs across 301 proteins (**Fig. 4A**, Table S4) at 43.7% sensitivity and 4.4% FDR (**Fig. 4B, 4C**, Methods: Short linear motif (SLiM) screening pipeline).

Comparing our candidate SLiMs with the Eukaryotic Linear Motif (ELM) resource (*29*), we found several of them to associate with known SLiM-binding proteins. Supporting our SLiM-discovery pipeline, most candidates that bind these proteins carried the expected motif class (**Fig. 4D**, Table S4): a PAM2 motif for the MLLE domain of PABP (6/8 candidates), an NLS for importin-α (3/4), a CanoR site for 14-3-3 (3/7), the canonical RVxF site for PP1 catalytic subunits in RRP1B (1/2) and NIPP1 (2/2), and the C-terminal EEVD motif for TPR co-chaperones (2/2).

Two large hubs bound coherent SLiM candidate motifs that are absent from ELM (**Fig. 4D**, asterisks; **Fig. 4E–F**):

First, the RACK1 WD40 scaffold, a constitutive component of the ribosomal 40S subunit, recruited seven cytoplasmic translation and stress-granule factors, five of which carry an aromatic-anchored S-[W/Y]-A-x-Φ motif and two a proline-rich P-P-P-K element (**Fig. 4E**, Table S4). Despite their distinct sequences, the two classes engage the same 13-residue patch on RACK1, with their consensus contact sites extending in opposite directions. This surface spans 33 residues and 1,848 Å², of which 99.3% remains solvent-accessible in the assembled 40S ribosome (**Fig. 4E**).

Second, ERH engaged ten binders: the known interactor DGCR8(*30*), the uncharacterized C19orf47, and eight RNA-related factors, all of which share a loose Val/Thr-rich, β-strand-like core. This is intriguing because β-strand augmentation is a known interaction mode of ERH. In the ERH-CIZ1 crystal structure (PDB 7X39), the β-strand residues 48–54 lack an intramolecular main-chain hydrogen-bonding partner, and this site is bound by a cognate CIZ1 β-strand (*31*). Analogously, 8 out of 10 SLiM candidates can dock on ERH at residues 48–54 by adding a regular β-strand (**Fig. 4F**, Table S4, Fig. S11), extending the binding partners of this previously unknown motif class.

### Cross-link-restrained AlphaFold modeling of large protein complexes

Complementary to mapping minimal binding sites, we used cross-links to guide the modeling of multi-protein assemblies. We developed a pipeline that combines cross-link-supported AF3x modeling of PPIs (*13*) with Monte Carlo hierarchical assembly into larger complexes(*32, 33*) (**Fig. 5A**, Methods: AF3x modeling with Monte Carlo hierarchical assembly). As illustrative examples, we applied this pipeline to functionally important complexes of the endosomal and secretory pathways that lack high-resolution human structures.

**Figure 5.**
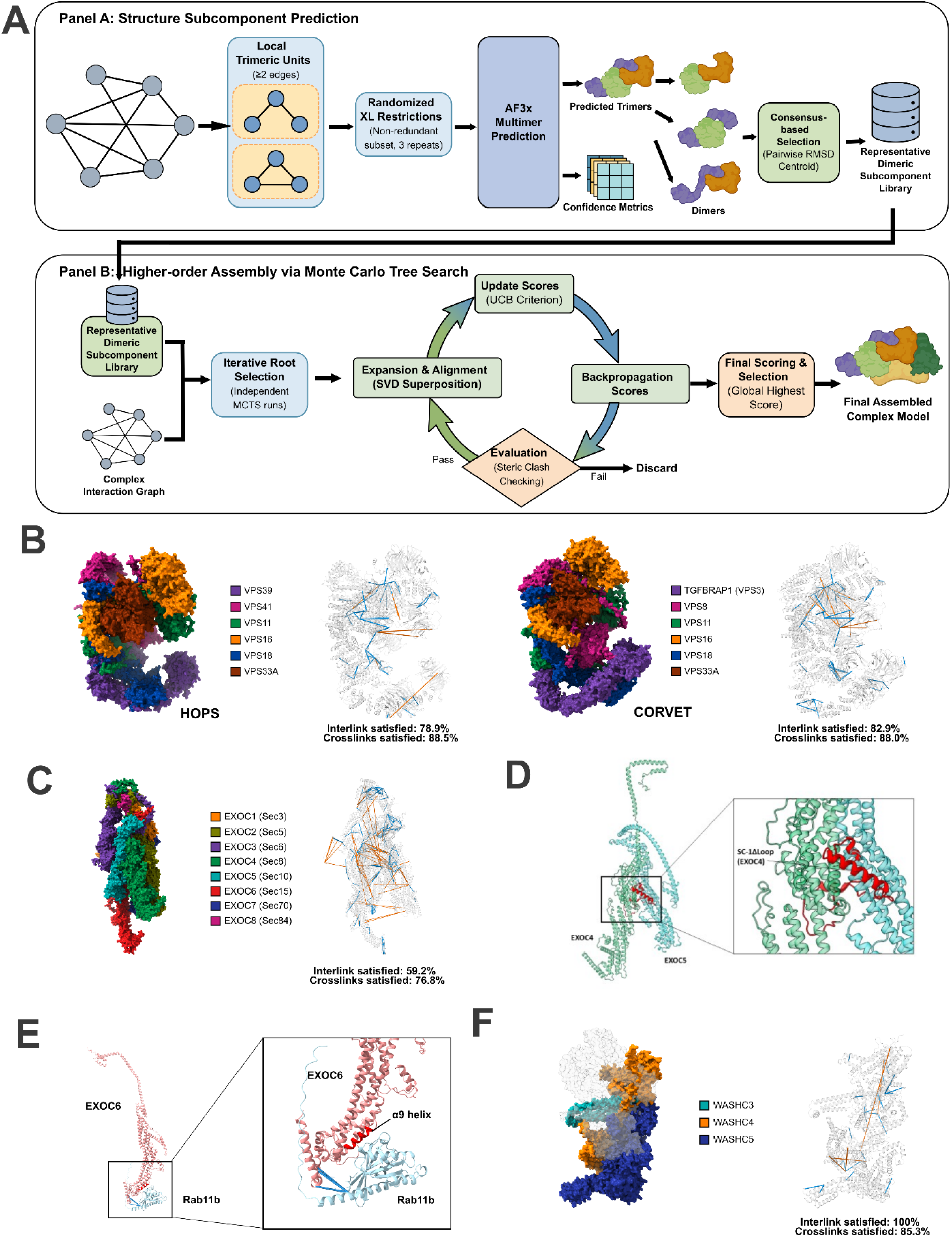
Structural modelling of multi-subunit protein complexes. (A) The modelling pipeline combines AF3x prediction with Monte Carlo–based assembly to generate structural models of multi-subunit protein complexes. Cross-links from DUC-XL-MS are used as spatial restraints within AF3x and the resulting subunit models are then assembled iteratively by Monte Carlo sampling to produce full-complex architectures. (B, C) Models of the human HOPS (B, left), CORVET (B, right) and exocyst (C) complexes. Subunits are shown in distinct colors in surface view. A 40 Å Cα–Cα distance threshold was used; satisfied cross-links are shown in blue and violated cross-links in orange. (D) Highlighted view of the EXOC4 C-terminal loop (red) interacting with EXOC5. (E) The predicted EXOC6–RAB11B model places the C-terminal region of EXOC6 in direct contact with RAB11B. The EXOC6 α9 helix (residues 690–702) is highlighted as the putative Rab11-binding element. EXOC6 is shown in pink, the α9 helix in red, RAB11B in light blue and the remaining exocyst subunits in gray. Cross-links between EXOC6 and RAB11B are shown in blue. (F) The predicted human WASH pentameric complex. WASHC3, WASHC4 and WASHC5 are shown in distinct colors in surface view; WASHC1 and WASHC2C are shown in gray and were excluded from the cross-link agreement calculation due to low model confidence (Supplementary Note 3). A 40 Å Cα–Cα distance threshold was used; satisfied cross-links are shown in blue and violated cross-links in orange.

The CORVET and HOPS complexes are homologous hexameric tethers of the endolysosomal system. They have a common class C core (VPS11, VPS16, VPS18 and VPS33) and are distinguished by complex-specific subunits: VPS3 (TGFBRAP1) and VPS8 in CORVET, and VPS39 and VPS41 in HOPS (*34*). Our models reproduced the core subunit arrangement seen in the yeast CORVET (*35*) and HOPS (*36*) structures, together indicating conservation from yeast to human (Supplementary Note 2). The cross-links further support the complex-specific subunits at defined terminal positions, with VPS3 binding VPS11 and VPS8 binding VPS18 and VPS33 in CORVET, and VPS39 binding VPS11 in HOPS (*37*) (**Fig. 5B, Fig. S12**). The CORVET-specific VPS3 and the HOPS-specific VPS39 cross-linked to the same region of VPS11 within the class C core (Fig. S12), illustrating that a mutually exclusive interaction contributes to the differentiation between CORVET and HOPS in humans.

The exocyst is an evolutionarily conserved octameric tethering complex essential for polarized exocytosis (*38*). Our assembly model places the EXOC4 C-terminal loop at the EXOC4–EXOC5 interface, supported by low PAE in this region (Fig. 5C, D, Fig. S13). This placement agrees with a published dimeric EXOC4–EXOC5 model, in which deletion of the EXOC4 C-terminal loop was shown to abrogate the EXOC4–EXOC5 interaction in vitro and in cells (*39*). We further identified a cross-link-supported interaction of RAB11 with the C-terminal domain of EXOC6 (Sec15) via the α9 helix (residues 690–702) (**Fig. 5E**). This same interface was also found in *Drosophila*, where it facilitates exocyst recruitment to Rab11-positive vesicles and polarized vesicle trafficking (*40*).

Finally, we modeled the WASH (Wiskott–Aldrich syndrome protein and SCAR homolog) complex (*41*), a pentameric endosomal activator of the Arp2/3 complex that drives branched actin polymerization and thereby regulates endosomal cargo sorting and the scission of transport carriers (*42*). No experimental structure of the assembled WASH complex has been reported, and the complex is absent from budding yeast (*43*). Our study therefore provides the first full model of the human WASH complex (**Fig. 5F**).

Assessing cross-link agreement allows us to evaluate the heterogeneity of the complexes and the accuracy of the models: intra-subunit links test each protein model, whereas inter-subunit links test how the subunits are arranged. We observed distinct cross-link agreement patterns across these complexes. In CORVET, HOPS and WASH, the violating cross-links formed discrete clusters confined to specific subunits or interfaces, indicating localized model inaccuracy or alternative conformations (Supplementary Note 3). In contrast, the exocyst had violations distributed across subunits. This pattern is consistent with the extended, minimally interacting and conformationally flexible architecture of the human exocyst reported in a recent structural study (*39*) (Supplementary Note 3).

### WASH accessory factors FKBP15 and ENTR1 promote its endosomal recruitment

The cross-link network of WASH revealed FKBP15 (FK506-binding protein 15) and ENTR1 (endosome-associated trafficking regulator 1) as accessory factors (**Fig. 6A**). FKBP15 is known to associate with the WASH–retromer network through the FAM21 (WASHC2) tail (*44*), but a direct interaction with the core subunits had not been reported. In our model, FKBP15 fitted onto the WASH complex close to the WASHC2C LFa-repeat region (**Fig. 6B**), in agreement with GST pull-downs from bovine brain extract (*44*).

**Figure 6.**
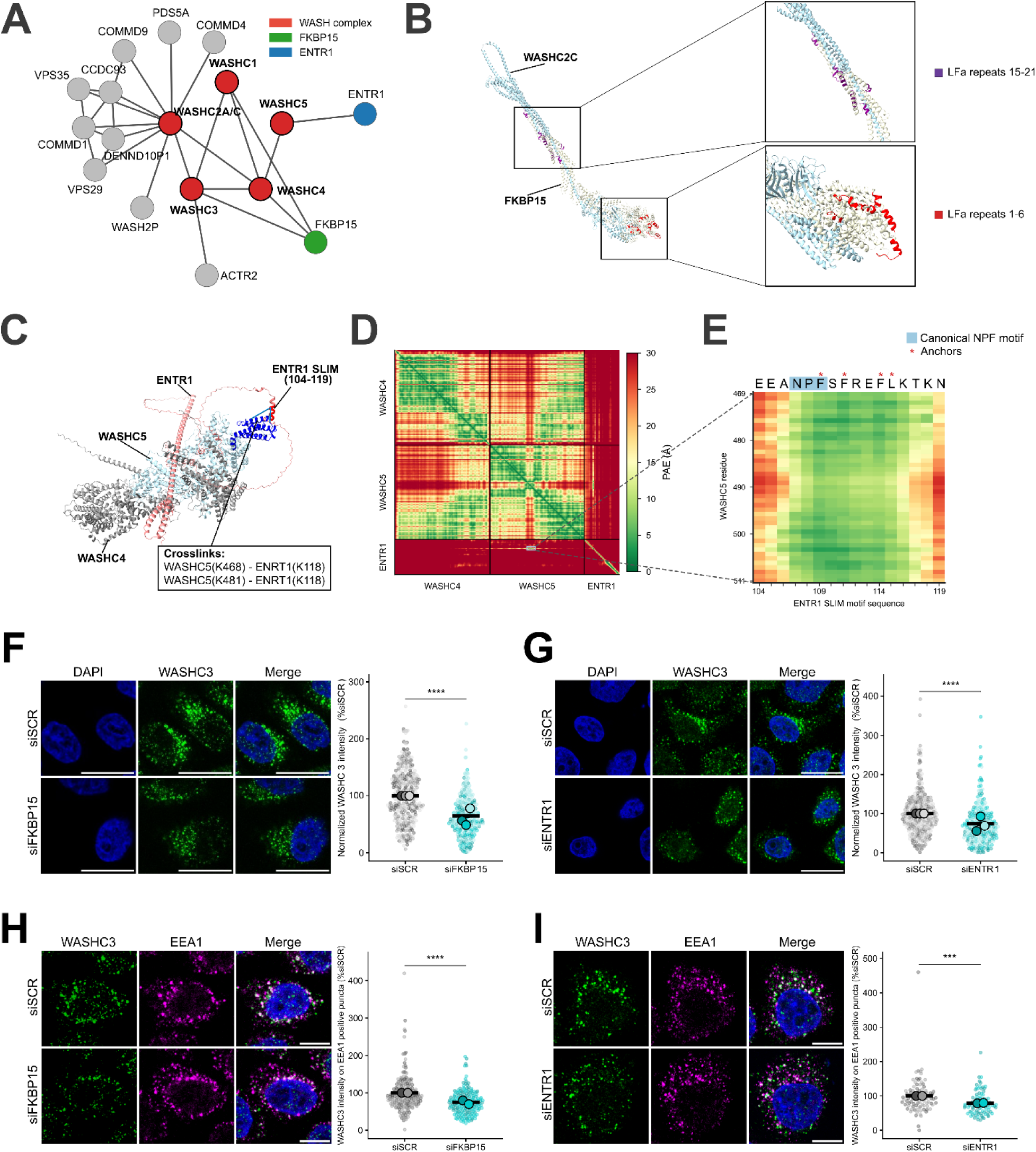
FKBP15 and ENTR1 are WASH-complex interactors that regulate endosomal WASH. (A) WASH core-complex first-tier interactors extracted from DUC-XL-MS, showing FKBP15 and ENTR1 directly cross-linked to WASH core subunits. (B) Protein-complex modelling places FKBP15 at the WASH complex, adjacent to the LFa-repeat regions of WASHC2C (*44*). WASHC2C, light blue; FKBP15, white; WASHC2C LFa repeats 1–6, red; LFa repeats 15–21, purple. (C) AlphaFold model of the WASHC4–WASHC5–ENTR1 complex; ENTR1 (salmon) with the SLiM (104–119) in dark red, WASHC5 in light blue, and WASHC4 in gray. Cross-links are indicated as blue lines. (D) PAE matrix of ENTR1 and WASHC5 interaction. The blue box marks ENTR1 104–119 × WASHC5 469–511. (E) SLiM sequence showing the canonical NPF motif (green) and hydrophobic anchors (stars). (F, G) Confocal microscopy of WASHC3 vesicles in HeLa cells transfected with siSCR and siFKBP15 (F) or siENTR1 (G), stained with anti-WASHC3. Scale bars, 20 µm. WASHC3 intensity per cell is normalized to the siSCR group (%siSCR). Small dots represent individual cells and large colour-coded dots represent replicate means from three independent experiments. Data are mean ± s.e.m.; two-tailed Mann–Whitney test; ****P < 0.0001. (F) n = 313 (siSCR) and 259 (siFKBP15) cells; (G) n = 303 (siSCR) and 245 (siENTR1) cells. (H, I) Left: confocal microscopy of WASHC3 vesicles co-stained with anti-WASHC3 and anti-EEA1 in siSCR and siFKBP15 (H) or siENTR1 (I) cells. Scale bars, 10 µm. Right: WASHC3 intensity on EEA1-positive area, normalized to siSCR (%siSCR); Small dots represent individual cells and large color-coded dots represent replicate means from two independent experiments. Data are mean ± s.e.m.; two-tailed Mann–Whitney test; ****P < 0.0001, ***P < 0.001. (H) n = 184 for both siSCR and siFKBP15 cells; (I) n = 88 (siSCR) and 85 (siENTR1) cells.

ENTR1 has been reported to colocalize with retromer-positive endosomes (*45*), but no direct interaction with the WASH core has been established. To corroborate this interaction, we collected a second DUC-XL-MS dataset from human Jurkat T lymphocytes. The Jurkat and HEK293 cross-links show a similar level of agreement with our WASH complex model (Supplementary Note 4) while increasing the density of residue-to-residue connections. Combining the two datasets yielded four ENTR1–WASH connections, one to WASHC4 and three to WASHC5 (Fig. S14). The ENTR1–WASHC5 interaction was independently annotated with high confidence in our proteome-wide SLiM screen (Table S4). Consistently, the ENTR1–WASHC4–WASHC5 trimeric model resolves a cross-link-supported interaction between the WASHC5 groove and a SLiM within ENTR1 (**Fig. 6C**). The SLiM region shows low PAE across the interface (**Fig. 6D**) and features a NPF motif (N107–P108–F109) with downstream hydrophobic anchors (F111, F114, L115) (**Fig. 6E**). NPF motifs are canonically recognized by EH domains, which are absent from WASHC5, adding it to the growing group of non-canonical NPF binders (*46*).

To assess how the associations of WASH with FKBP15 and ENTR1 influence its assembly and recruitment, we analyzed the distribution and nanoscale localization of WASH in cells depleted of either endogenous ENTR1 or FKBP15 (Fig. S15) by confocal imaging. Loss of either interactor led to an overall reduced intensity of WASH endosomal puncta (**Fig. 6F,G**) and decreased co-localization of WASHC3 with EEA1-marked early endosomes (**Fig. 6H,I**). WASH cellular abundance remained constant (Fig. S15), indicating that FKBP15 and ENTR1 specifically regulate WASH function through its assembly and endosomal recruitment.

### Systems biology augmentation defines a spatiostructural human core interactome

The analyses above highlight the ability of DUC-XL-MS to reveal spatial, structural and functional aspects of the human proteome that are not captured in published interactome resources. To understand the factors driving this complementarity, we scrutinized the dataset structures of DUC-XL-MS and EndoMAP (both XL-MS-based) as well as BioPlex 3.0 (HEK-only) and OpenCell (both AP-MS-based).

We first compared the organelle coverage of the four networks (**Fig. 7A**). BioPlex contains the most PPIs, followed by OpenCell and DUC-XL-MS, and its particularly deep coverage of the plasma membrane interactome reflects its many plasma membrane baits and the enrichment inherent to AP-MS. The organelle distribution of DUC-XL-MS proteins and PPIs most closely resembles that of OpenCell (**Fig. 7A**, Fig. S16). Notably, DUC-XL-MS identifies similar numbers of endosomal proteins and interactions to the dedicated EndoMAP network built on XL of affinity-purified endosomes, yet the overlap between them is low (**Fig. 7A, B**, Supplementary Note 5). DUC-XL-MS recovers substantially more of the cytosol-facing endosomal machinery than EndoMAP XL, spanning ESCRT-dependent multivesicular body sorting, endocytic recycling and retrograde retrieval, and macroautophagy, indicating that DUC-XL-MS preserves the transient interactions lost during endosome enrichment (Supplementary Note 5).

**Figure 7.**
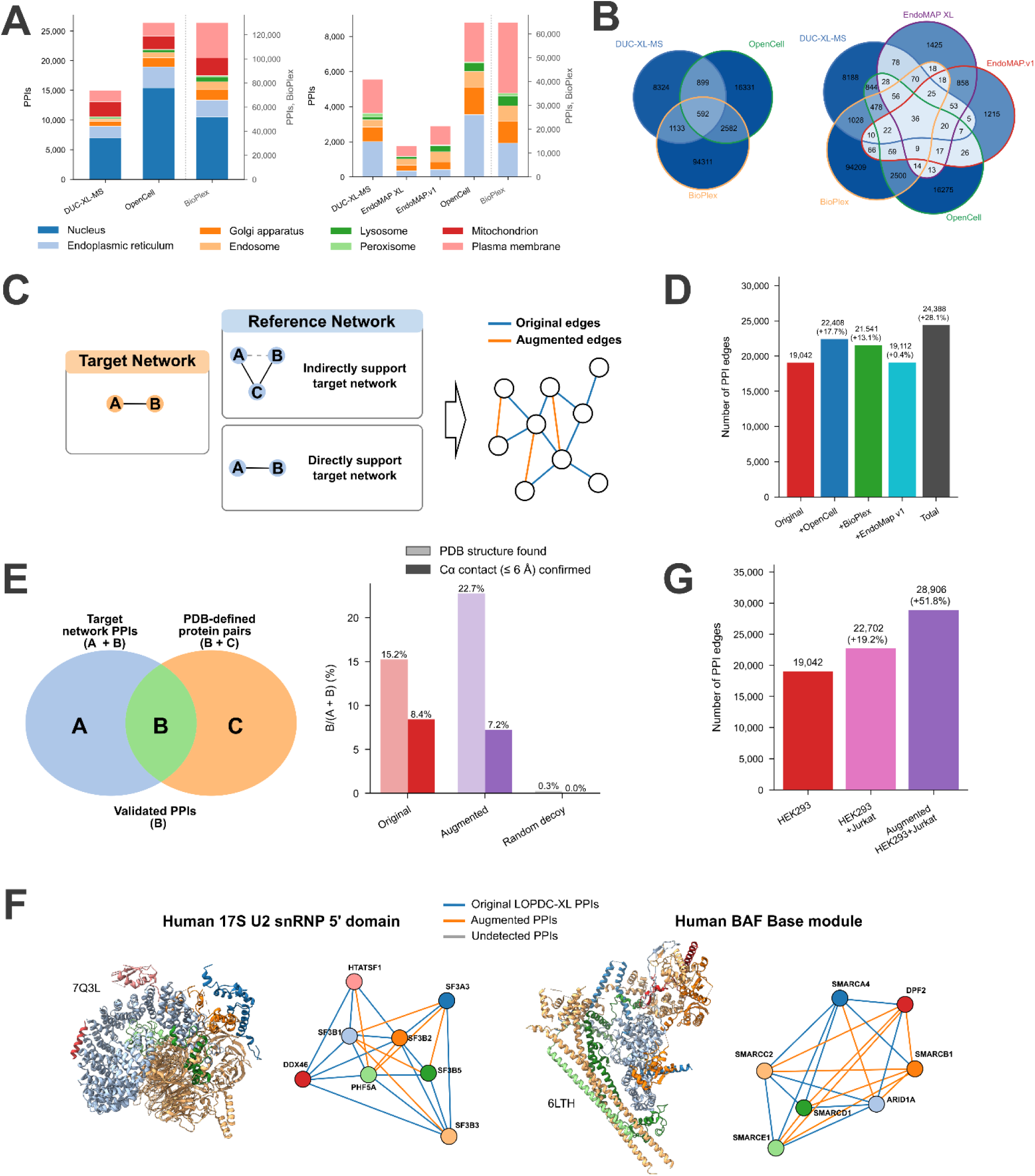
Systems biology augmentation expands the high-confidence human interactome. (A) Organelle coverage of PPIs across datasets. Stacked bar charts showing, for each dataset, the numbers of organelle-associated PPIs assigned to eight subcellular compartments (left) and six organelles (excluding nucleus and mitochondria) (right). PPIs are counted once per annotated compartment, so multi-localized entities contribute to multiple compartments. BioPlex is plotted on a separate right-hand axis. Organelle coverage of cross-linked proteins across datasets are shown in Fig. S16. (B) Overlap between PPIs reported in A after deduplication. Full pairwise-overlap statistics are provided in Fig. S17. (C) Triadic-closure principle for comparing PPI networks. Triadic closure provides a graph-theoretic basis for comparing PPI networks generated by different methods. When a target network is compared with a reference network, an edge A–B in the target is directly supported if the same A–B interaction is detected in the reference. In the absence of direct detection, A–B is indirectly supported if a shared-neighbor protein C exists such that both A–C and B–C are present in the reference, satisfying the triadic-closure condition. (D) Network augmentation by triadic closure. The DUC-XL-MS network is expanded with each reference dataset: an edge is added whenever its two proteins share a common neighbor in the network, closing the triad. (E) Structural validation of augmented edges. (Left) Structural support is defined as the number of PDB-validated PPIs divided by the total number of PPIs in the target network, counted only over pairs in which both proteins are present in the PDB. (Right) Bar charts show the percentage of structural support for three interaction categories: original edges (DUC-XL-MS interactions with direct experimental evidence); augmented edges (edges added by triadic closure from three databases (EndoMAP, BioPlex 3.0 or OpenCell 2.0), and random decoys (protein pairs sampled from the same degree distribution, as a negative control). Two metrics are shown per category: the fraction of pairs whose two proteins co-occur in a single PDB entry (PDB hit rate) and the fraction with a minimum interatomic distance ≤ 6 Å (direct-contact rate). (F) Example cases of edge augmentation. For two representative structures (PDB 7Q3L and 6LTH), the PPIs recovered by the network built from original edges alone are compared with those recovered after edge augmentation. A PPI is defined as a protein pair with a minimum inter-atomic distance ≤ 6 Å in the corresponding structure. Edge color indicates recovery status: blue, PPIs recovered by the original network; orange, PPIs recovered only by the augmented network; gray, PPIs recovered by neither. (G) The pooled, augmented human interactome. Cumulative PPI count as DUC-XL-MS data from HEK293 and Jurkat cells are pooled and then augmented with the reference datasets (EndoMAP, BioPlex 3.0 and OpenCell).

Next, we assessed PPI overlap across all datasets. We found uniformly low PPI overlap (**Fig. 7B**, Fig. S17), even when comparing datasets generated with the same method and focusing on proteins shared between resources. These results reveal an inherent complementarity of different PPI datasets. To harness this complementarity, we developed a data integration strategy that leverages network topology beyond direct pairwise detection. We applied a graph-theoretic triadic-closure approach (**Fig. 7C**), reasoning that network-level topology can recover missing but biologically supported interactions. Triadic closure posits that two proteins sharing a common partner are more likely to interact directly. Accordingly, protein pairs absent from our DUC-XL-MS data but present in at least one of three reference datasets (EndoMAP, BioPlex 3.0, or OpenCell) were defined as candidate augmented edges if they shared at least one direct interactor within the DUC-XL-MS network. This approach expanded our network by 28% (**Fig. 7D**, Table S5). Conversely, the other datasets can likewise be augmented by one another as well by DUC-XL-MS, confirming that each resource contributes information the others miss (Fig. S18).

To assess whether these augmented edges represent genuine physical associations, we tested them against PDB structures (**Fig. 7E**). Augmented edges recovered a larger fraction of the PPIs that could be mapped onto a PDB structure than original edges did, though they were slightly less likely to form a direct interface contact, defined following BioPlex 3.0 as a minimum interatomic distance of 6 Å or less (*1*). Both original and augmented edges scored far above random protein pairs. These results indicate that augmentation covers the interactions within complexes more completely. To illustrate the practical value of these edges, we examined four medium-sized complexes (6 to 8 subunits) with extensive PDB and DUC-XL-MS coverage (**Fig. 7F**, Fig. S19). In every case, augmented edges added bona fide PPIs that the original DUC-XL-MS network had missed.

Finally, we assessed the complementary value of distinct cell-line interactomes. They are present in BioPlex 3.0 (HEK293T and HCT116) and in our DUC-XL-MS resource (HEK293 and Jurkat) (Table S1). The PPI overlap between two cell lines profiled by the same method is only modest (Jaccard similarity of 23.3% for BioPlex and 24.6% for DUC-XL-MS) (Fig. S20). Nevertheless, the two DUC-XL-MS datasets showed similar accuracy both structurally (mapping cross-links onto PDB structures) (Fig. S21) and spatially (cross-validation against organelle and topology markers) (Fig. S22).

This indicates that PPI data from different cell lines, just like PPI data from different resources, represent complementary interactome segments of comparable confidence. Adding the Jurkat data increases PPI coverage by 19.2% relative to HEK293 alone (**Fig. 7G**, Table S5). Augmenting this pooled network with the available resources (EndoMAP, BioPlex from both cell lines, and OpenCell) further increases the PPI by 51.8%, yielding a human structural core interactome comprising 28,906 PPIs (**Fig. 7G**, Table S5).

## Discussion

We developed DUC-XL-MS to chart a cellular structural interactome that expands our understanding of the human proteome along three dimensions. First, its spatial analysis annotates proteins to organelles and organellar interfaces, reveals regulatory interactions through inter-organelle cross-links, and resolves the topology of the membrane proteome. Second, cross-link-restrained AlphaFold modeling yields structural insight into large protein complexes and uncovers the short linear motifs that underlie much of the human PPI network. Third, systems-biology augmentation of complementary interactomics data expands the DUC-XL-MS network into the largest spatiostructural human interactome reported to date, providing a framework for functional studies guided by proteome architecture.

Representative for its spatial informativity, DUC-XL-MS enabled the systems-scale dissection of subcellular protein-interaction networks and their rewiring upon genetic manipulation of key interaction nodes. We identified LRBA, a protein linked to primary immunodeficiency with autoimmunity (*22*), as a chaperone that maintains the proper localization of the endosomal fusion protein STX7. Mislocalization of STX7 may thus contribute to the severe primary immunodeficiency (hypogammaglobulinemia, autoimmunity and T-cell dysfunction) caused by biallelic LRBA loss-of-function mutations in humans (*22*). Similarly, we showed that the localization and function of the actin-regulatory WASH complex at endosomes are controlled by structurally distinct direct interactions with FKBP15 and ENTR1. As these changes in the localization and interactomes of STX7 and WASH do not alter their cellular abundance, they would have been missed by conventional whole-cell proteomics, illustrating the added value of DUC-XL-MS.

On the structural level, DUC-XL-MS enabled interaction motif discovery and the reconstruction of large protein assemblies from direct pairwise distance restraints. Direct interface information sharply reduces the dimer library and the docking search space for structure modeling, which allowed us to screen SLiM candidates efficiently at proteome scale. At the larger scale, AF3x modeling together with Monte Carlo hierarchical assembly predicted the structures of large complexes from XL-MS-derived dimer libraries. Albeit satisfactory for CORVET, HOPS and WASH, cross-links alone can, however, be insufficient for complete modeling, particularly when stoichiometry, compositional heterogeneity and oligomeric state confound cross-link interpretation (Supplementary Note 6).

We anticipate that our systems level view of the human interactome will keep expanding as further datasets are acquired and the sensitivity of MS-based proteomics improves. In agreement with previous studies (*10, 47*), XL-MS as reported here and AP-MS approaches such as BioPlex 3.0 and OpenCell yield a common core of PPIs but are otherwise highly complementary. AP-MS excels at low-abundance interactors but loses weak, transient organellar interactions during lysis and purification; proximity-driven XL-MS preserves these and reports direct PPIs and their interfaces. The same distinction holds within XL-MS itself: DUC-XL-MS, which cross-links a mildly lysed whole-cell preparation, recovers substantially more of the transient, cytosol-facing endosomal machinery, including endocytic recycling, retrograde retrieval and macroautophagy proteins, than EndoMAP, which cross-links endosomes after affinity enrichment with EEA1 as the bait (Supplementary Note 5).

At present, the space of possible PPIs in a human cell still exceeds the fraction that any single experiment samples, so even datasets based on the same interactomics approach capture distinct and sparsely overlapping slices of the interactome. This pervasive complementarity across methods and datasets motivates integrative pipelines that assemble high-confidence interactomes by augmentation strategies such as our triadic closure approach, converting the low pairwise overlap from a liability into a route toward more complete coverage. In parallel to this study, we combined organellar and cellular XL-MS datasets to build the most comprehensive cross-link map of the human Nuclear Pore Complex to date. Integrated with cryo-electron tomography and AI-assisted modeling, this identified five proteins not previously annotated as nucleoporins (TMEM209, SMPD4, GANP, Centrin-2 and ENY2) and established the TREX-2 complex (GANP, Centrin-2, ENY2) as an integral structural module of the nuclear ring (*48*).

By uniting spatial, structural and systems-level readouts in a single experiment, DUC-XL-MS offers a blueprint for mapping how the human proteome is organized, assembled and rewired, in both health and disease. Nonetheless, the present study, like the AP-MS resources BioPlex and OpenCell remains qualitative. Extending XL-MS to quantitative comparisons while preserving deep interactome coverage will require more sensitive workflows compatible with metabolic or chemical labeling or data-independent acquisition, together with faster LC-MS instrumentation (*49*). Combined with deeper fractionation and continued gains in cross-linker enrichment, quantitative XL-MS at depth is the natural next step toward resolving the dynamic, condition-dependent proteome architecture.

## Materials and Methods

### DUC-XL-MS experiments

For the HEK dataset, cells were washed with PBS and resuspended in lysis buffer (250 mM sucrose, 10 mM HEPES, 2 mM EDTA, 2 mM magnesium acetate tetrahydrate, pH 7.4) supplemented with complete EDTA-free protease inhibitor cocktail. Cells were then lysed with a homogenizer at 1,000 rpm until the lysis efficiency reached 90%. Cell debris was pelleted at 200 × g for 5 min at 4 °C. The supernatant was collected and its protein concentration was determined with the BCA protein assay kit (Thermo Fisher Scientific) according to the manufacturer’s instructions. The lysate was adjusted to 10 mg/mL with lysis buffer and cross-linked with 2 mM Azide-A-DSBSO for 15 min at room temperature with constant mixing. The reaction was quenched with 20 mM Tris-HCl (pH 8.0) for 30 min at room temperature with constant mixing.

The cross-linked samples were fractionated by differential centrifugation to reduce proteome complexity (Fig. S1). In brief, samples were first clarified in a benchtop centrifuge at 1,000 × g for 30 min to pellet nuclei, unbroken cells and large debris (fraction F1). The resulting supernatant was then subjected to a series of sequential ultracentrifugation steps in 10-mL tubes using a Beckman Coulter SW40 Ti swinging-bucket rotor, each performed at 4 °C. Pellets were collected at successively increasing speeds and durations: 4,100 rpm for 30 min (F2; ∼3,000 × g), 5,300 rpm for 30 min (F3; ∼5,000 × g), 7,100 rpm for 35 min (F4; ∼8,900 × g), 8,200 rpm for 35 min (F5; ∼11,900 × g), 9,200 rpm for 35 min (F6; ∼15,000 × g), 13,000 rpm for 40 min (F7; ∼30,000 × g), 21,100 rpm for 63 min (F8; ∼79,000 × g) and 26,000 rpm for 65 min (F9; ∼120,000 × g). After the final step, the remaining supernatant was retained as the cytosol-enriched fraction (F10). Each pelleted fraction (F1–F9) and the cytosolic fraction (F10) was collected separately and stored until downstream analysis.

For the Jurkat dataset, cells were cultured in RPMI 1640 medium (with D-glucose, L-glutamine, sodium bicarbonate and sodium pyruvate; catalog #A10491-01) supplemented with 10% FBS, at 37 °C and 8% CO□, on an orbital shaker at 100 rpm. Cross-linking was performed with 4 mM Azide-A-DSBSO for 30 min at room temperature with constant mixing. All other steps were as described for the HEK dataset.

#### Protein digestion, cross-link enrichment and off-line HPLC fractionation

Each pellet was resuspended in 8 M urea in 50 mM TEAB for denaturation, then reduced and alkylated. Proteins were digested with Lys-C at an enzyme-to-protein ratio of 1:75 (w/w) for 4 h at 37 °C. After dilution with 50 mM TEAB to a final concentration of 2 M urea, trypsin was added at 1:100 (w/w) and digestion was continued overnight at 37 °C. For the HEK dataset, digestion was quenched by adding formic acid to a final concentration of 1%; peptides were then desalted with Sep-Pak C18 cartridges (Waters) according to the manufacturer’s protocol and dried in a vacuum concentrator. For the Jurkat dataset, the digested peptide mixture was subjected directly to DSBSO enrichment.

The digested peptide mixtures were enriched using dibenzocyclooctyne (DBCO)-coupled Sepharose beads. Briefly, the digested peptides were added to prewashed DBCO beads and incubated overnight at room temperature with constant mixing. After incubation, the beads were washed with water, followed by 0.5% SDS at 37 °C for 15 min. Next, the beads were washed three times with 0.5% SDS, three times with 8 M urea in 50 mM TEAB, three times with 10% ACN and twice with water, using ten bead volumes each. The cross-linked peptides were eluted with two bead volumes of 10% (v/v) trifluoroacetic acid (TFA) for 2 h at room temperature (HEK dataset) or 2% (v/v) TFA for 2 h at room temperature and additional two bead volumes of 80% ACN (Jurkat dataset) and then dried in a vacuum concentrator. The enriched cross-links were further fractionated by size-exclusion chromatography (SEC) followed by high-pH fractionation. SEC was performed on a Superdex 30 Increase 3.2/300 column (GE Healthcare) using an Agilent 1260 Infinity II system, and high-pH fractionation was performed on a Phenomenex Gemini C18 column using an Agilent 1260 Infinity II UPLC system (HEK dataset) or on a Vanquish C18+ column (1.5 µm, 2.1 × 50 mm; Thermo Fisher Scientific) using a Vanquish HPLC system (Jurkat dataset). The high-pH fractions were subjected to LC-MS analysis.

#### LC-MS

For the analysis of DSBSO-cross-linked peptides, collected fractions were analyzed by LC-MS on an UltiMate 3000 RSLCnano system coupled online to an Orbitrap Fusion Lumos or an Orbitrap Exploris 480 mass spectrometer (Thermo Fisher Scientific) for the HEK or Jurkat dataset, respectively. Reversed-phase separation was performed on an in-house-packed C18 analytical column (Poroshell 120 EC-C18, 2.7 µm; Agilent Technologies), and fractions were run with 3-h LC gradients. 384 fractions for HEK and 226 fractions for Jurkat cells were measured. The following MS parameters were used: MS resolution, 120,000; MS2 resolution, 60,000; charge states 4–8 enabled for MS2; MS2 isolation window, 1.6 m/z; stepped normalized collision energy, 19/25/30%; and FAIMS compensation voltages, −50, −60 and −75 V.

#### Data analysis

Data analysis was performed with pLink v3.1.8 (https://github.com/pFindStudio/pLink3) using the following parameters: minimum peptide length, 6; maximum peptide length, 60; and missed cleavages, 3. The fixed modification was cysteine carbamidomethylation, and the variable modifications were methionine oxidation and protein N-terminal acetylation. The DSBSO cross-linker mass was specified as 308.0388 Da (short arm, 54.0106 Da; long arm, 236.0177 Da). The precursor and fragment mass tolerances were set to 10 ppm and 20 ppm, respectively. MS2 spectra were searched against a reduced target-decoy Swiss-Prot human database derived from the proteomic measurements of all fractions. The HEK and Jurkat databases contained 6,232 and 7,249 proteins, respectively, corresponding to the proteins identified from all fractions prior to DSBSO enrichment. For both datasets, pLink3 results were filtered to a 1% FDR at the residue-pair level. The HEK dataset comprises 164,146 residue pairs and 18,074 inter-protein PPIs across 5,399 proteins, and the Jurkat dataset comprises 124,835 residue pairs and 9,391 inter-protein PPIs across 5,710 proteins (Table S1).

### Protein organelle localization and membrane topology analysis

#### Organelle marker assignment

Protein subcellular localizations were derived from the UniProtKB “Subcellular location [CC]” field for proteins in the cross-link network. Within each protein’s CC annotation, membrane-topology and attachment annotations (single-pass, multi-pass and peripheral membrane protein; lipid- and GPI-anchor; and membrane-side descriptors) were discarded, and the top-level compartment (the term preceding the first comma) was retained and mapped, through a manually curated dictionary (Table S2), onto a set of canonical compartments: nucleus, cytoplasm, mitochondrion, endoplasmic reticulum (ER), ER-Golgi intermediate compartment (ERGIC), Golgi apparatus, endosome, lysosome, peroxisome, plasma membrane, secreted and lipid droplet. A catch-all “other” category was assigned to non-organelle or spatially ambiguous terms (cell projection, cell junction, midbody, cleavage furrow, melanosome, synapse, generic “membrane,” vacuole and virion).

Each protein’s parsed annotation was reduced to its set of canonical organelle compartments; “other” terms were discarded. A protein was classified as single-localized when it resolved to exactly one of the eight organelle compartments (nucleus, mitochondrion, ER, Golgi, endosome, lysosome, peroxisome or plasma membrane) and carried no cytoplasm annotation. Proteins annotated solely as cytoplasm, secreted or lipid droplet were removed.

A protein was classified as dual-localized when it resolved to exactly two of the seven cytoplasmic organelles (mitochondrion, ER, Golgi, endosome, lysosome, peroxisome and plasma membrane; the nucleus is excluded, and the ERGIC counts as ER plus Golgi) and those two compartments form an adjacency-allowed pair (Fig. 1C).

To obtain the final list of organelle markers, vesicle-pool and ambiguously cross-linked proteins (see section “Exclusion of promiscuously cross-linked proteins”) were removed from the marker list, and the remaining markers were curated after validating cross-link spatial consistency (see section “Validation of cross-link spatial arrangement”).

#### Topology marker assignment

Protein subcellular localizations were derived from the UniProtKB “Subcellular location [CC]” field for proteins in the cross-link network. Compartments were reduced to five topological sides: the cytoplasmic-nucleoplasmic continuum (CYT+NUC), joined through nuclear pores; the exoplasmic/lumenal side (LUM), comprising the ER, Golgi, endosomal and lysosomal lumen together with the secreted and extracellular space; the mitochondrial matrix (MTX); the mitochondrial intermembrane space (IMS); and the peroxisomal matrix (PXM). Topology markers were proteins assigned unambiguously to a single side; proteins resolving to two or more sides, or to none, were not used. Transmembrane proteins were excluded. For soluble mitochondrial proteins, IMS and MTX annotations were taken from our published cross-link-derived sub-compartment analysis (*9*) where available, and otherwise from an explicit UniProt “matrix” or “intermembrane space” annotation. A bare “Mitochondrion” label in Uniprot was not used.

To obtain the final list of topology markers, promiscuously cross-linked proteins (see section “Exclusion of promiscuously cross-linked proteins”) were removed from the marker list, and the remaining markers were curated after validating cross-link spatial consistency (see section “Validation of cross-link spatial arrangement”).

#### Exclusion of promiscuously cross-linked proteins

Several protein classes cross-link non-specifically across compartments, so that a cross-link to them reports protein abundance or a shared trafficking pool rather than a specific subcellular location. These were removed by a single exclusion rule applied across the marker sets and the propagation predictions: organelle markers and organelle propagation excluded all four classes below, whereas topology markers excluded only classes ii, iii and iv. The classes are: (i) vesicular-carrier proteins, which are transient trafficking intermediates rather than residents of a single organelle (identified by any vesicular-carrier term in the UniProt localization text, including coated pit, coated vesicle, cytoplasmic vesicle, clathrin, COPI, COPII, secretory vesicle, transport vesicle or “vesicle”); (ii) the ubiquitous post-translational modifiers ubiquitin, SUMO and NEDD8, which are conjugated to substrates proteome-wide; (iii) histones (linker H1, the core histones H2A, H2B, H3 and H4, and the macroH2A variants); and (iv) the highly abundant cytosolic ribosomal proteins and cytosolic molecular chaperones (for example the HSP70 and HSP90 families and the chaperonin CCT/TCP1), which engage nascent and misfolded clients promiscuously throughout the cytoplasm. Organelle-resident members of these families that themselves mark a compartment, namely the mitochondrial ribosomal proteins and the ER-luminal chaperones (calnexin, BiP/HSPA5, GRP94/HSP90B1 and the protein disulfide isomerases), were retained as markers.

#### Validation of cross-link spatial arrangement

*Organelle-level consistency:* inter-organelle connections were defined a priori to reflect established membrane-contact sites and trafficking routes (Fig. 1C): the ER was permitted to connect to any compartment (“ER-everything”), together with endosome-Golgi, endosome-lysosome, endosome-mitochondrion, endosome-plasma membrane, Golgi-lysosome, lysosome-mitochondrion, lysosome-plasma membrane and mitochondrion-peroxisome; the nucleus was permitted to contact only the ER. For each protein pair, a cross-link was classified as allowed if the two proteins’ localization sets shared a permitted connection; as not allowed (a conflict) if the proteins occupied distinct compartments with no permitted connection between them; and as unannotated if either protein lacked a localization assignment. Proteins reassigned by manual curation were listed in Table S2.

*Topology-level consistency*: the five topological sides are separated by membranes, so the two residues of a cross-link must lie on the same side. Each marker-to-marker cross-link was therefore classified as topologically possible only when both residues occupied the same side. Proteins reassigned by manual curation were listed in Table S2.

#### Graph-based propagation of protein localization

Protein localization was inferred by propagating the curated organelle markers across the cross-link interaction network, on the assumption that cross-linked proteins tend to occupy the same compartment. Each annotated compartment of a marker was counted once per neighbor. A compartment assignment was considered valid only if supported by at least two independent markers, or by a single marker linked through at least two distinct residue pairs. For each protein, the score for a given compartment was defined as the number of markers supporting that compartment divided by the total number of markers across all of the protein’s valid compartments. A protein whose top compartment reached a score of at least 0.80 was assigned to that single compartment. A protein with two compartments each scoring at least 0.30, provided those two compartments form an allowed inter-organelle adjacency, was assigned as dual-localized. Proteins receiving only weak or split evidence that met none of these criteria were left unclassified. Three categories were flagged as unreliable and excluded from the confident set: (1) proteins supported by four or more valid compartments, (2) single-compartment proteins with a high network degree (at least ten cross-link partners) whose top compartment supported by only one marker, and (3) the promiscuously cross-linked proteins described in an earlier sub-section (cytosolic ribosomal proteins, cytosolic chaperones, ubiquitin, SUMO and NEDD8, and histones). The full protein annotation list after propagation is in Table S3.

#### Topology analysis of transmembrane proteins

For each transmembrane protein, the UniProt transmembrane helices (TRANSMEM) were used to partition the sequence into alternating soluble loops (the N-terminal segment, each inter-helical loop, and the C-terminal segment). A loop was assigned a side if at least 80% of its cross-linked residue pairs (minimum: 2 pairs) supported the assignment. Loops with fewer than two residue pairs, or without an 80% majority, were left unassigned.

Loops were annotated over two rounds. In the first round, only the curated, soluble topology markers were used to support assignments. In the second round, each loop whose first-round assignment matched the protein’s UniProt topology annotation was added to the topology marker list to extend the side assignment to more transmembrane-protein loops.

Each assigned loop was compared with the protein’s own UniProt topological domains (TOPO_DOM) and scored as agreeing (sides match), conflicting (sides differ), or novel (no UniProt annotation for that loop). Proteins were classified as validated (all annotated loops agree), conflict (at least one annotated loop conflicts), or prediction (only novel loops). For proteins with at least two assigned loops, the assigned sides were checked for consistency with a single alternating topology. Proteins that violated this alternation (for example both flanks of a single-helix protein on the same side) were flagged as internally inconsistent, typically indicating a spurious transmembrane helix or a contested topology. The full topology annotation list is in Table S3.

### Short linear motif (SLiM) screening pipeline

#### AlphaFold3 predictions

AlphaFold3 was used to model cross-linked protein pairs using the default inference pipeline with standard MSA and template search (*26*). The resulting pair models were filtered to those carrying at least one confidently modelled inter-chain contact, defined as a minimum inter-chain predicted aligned error (MiniPAE) (*25*) of at most 4 Å (MiniPAE ≤ 4 Å). For every protein, an unbound monomer was also modelled.

#### Interface scoring and localisation with ipSAE

Each protein pair interface was scored with the Interaction Prediction Score from Aligned Errors (ipSAE) (*27*) in its residue-specific form (d0res) at a 15 Å PAE cutoff, following the vectorised reference implementation. For a residue i on one chain, valid partners are those residues j on the opposite chain with PAE_ij_ < 15 Å. With n□ such partners, the local scaling factor is

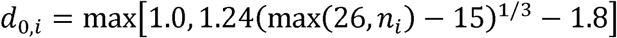

and the residue score is the mean over valid partners of the pTM-style transform

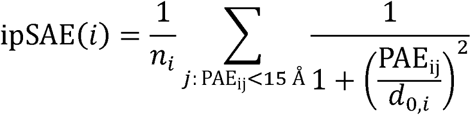

ipSAE therefore returns a per-residue vector that both localizes and scores the interface. Restricting the score to residue pairs that pass the PAE cutoff avoids diluting it with disordered flanks and uncoupled domains. This is critical because a SLiM complex consists of a short peptide bound to a large, folded domain, so a single per-complex score would be dominated by residues outside the interface.

On each chain, residues with ipSAE ≥ 0.10 were grouped into segments, allowing gaps of at most two positions between consecutive residues, and segments of 3 to 30 residues were retained as interface segments. For every protein pair passing the MiniPAE filter, all interface segments from both chains were exported.

### Binding-induced folding (ΔpLDDT) and intrinsic disorder (flanking pLDDT)

#### Score definitions

ΔpLDDT was defined as the mean Cα pLDDT (predicted Local Distance Difference Test) over the interface segment residues in a protein pair model minus the mean over the identically numbered residues in the corresponding monomer model. This binding-induced folding score follows the pLDDT gain between the unbound and bound states used by AlphaSLiM (*50*). The principle that intrinsically disordered regions fold conditionally, as reflected in AlphaFold confidence, was previously established (*28*). Flanking pLDDT was defined as the mean unbound-monomer pLDDT of the residues within 10 positions on each side of an interface segment (N- and C-terminal), each flank averaged over whatever residues were present (up to 10). If an interface segment was located at one of the protein termini without any flanking residues, it was scored on its single existing flank. Together these scores capture the defining property of a SLiM: a short segment that is disordered in isolation and folds upon binding (ΔpLDDT) (*50*) and is embedded in a disordered context (flanking pLDDT) (*51, 52*). To set appropriate score thresholds, positive and negative controls were designed.

#### Positive control

The 694 interacting pairs annotated in the ELM database (*29*) were predicted as AlphaFold3 dimers, together with the corresponding monomers. The fraction of these 694 pairs whose designated SLiM passed the pipeline (the pass rate) served as the positive control for measuring sensitivity across different ΔpLDDT and flanking pLDDT cutoffs.

#### Negative control

For every interface segment, a control segment was selected from the same chain of the same dimer model, matched in length and in mean unbound-monomer pLDDT (±5) and containing no interface residue. This control is true-negative because a segment that contacts no interface cannot fold on binding. The rate at which this negative control passes a given score threshold, thus, estimates the false-positive background. Matching on monomer pLDDT excludes segments in which a large ΔpLDDT is absent only because the segment is already ordered in the monomer. Of the 11,622 interface segments, 11,400 had an eligible matched window; the remaining 222 were removed from the interface count as well as from the control count.

#### Determining ΔpLDDT and flanking pLDDT thresholds

We screened a range of ΔpLDDT and flanking pLDDT thresholds, using the positive and negative controls to estimate sensitivity and FDR, respectively. For ΔpLDDT, the FDR falls steeply between +15 and +18 and then plateaus, while sensitivity declines only gently (44.8% → 43.7%); a ΔpLDDT threshold of +18 was therefore selected. For flanking pLDDT, a cutoff of 70 was used, corresponding to AlphaFold’s standard threshold separating confident (pLDDT ≥ 70) from low-confidence predictions. When these thresholds are applied to the 11,400 matched segments, 4.18% of interface segments and 0.18% of matched controls pass. This corresponds to a 23-fold separation (4.18% ÷ 0.18%) and an estimated FDR of ≈4.4%, calculated as the control pass rate divided by the interface pass rate (0.18% ÷ 4.18%), since the length- and pLDDT-matched controls are true negatives that pass only by chance.

#### Recovery of known motif classes

For motif classes whose cognate SLiM-binding protein is present in the network, each ELM class regex (release 1.4, 353 classes, downloaded 2026-07-19 (*29*)) was matched against the full monomer sequence of the candidate protein and required to overlap the predicted segment. Matching the full sequence rather than the segment alone is necessary because several ELM classes specify flanking positions that fall outside the tightly localized ipSAE segment. Each candidate SLiM was tested against every ELM class read by its partner’s domain. Counts are reported as proteins, because ELM class membership is a property of the protein and multiple segments on one protein at the same reader represent a single interaction.

### AF3x modeling with Monte Carlo hierarchical assembly

#### Structural modelling of protein complexes

Trimeric complex units were extracted from our DUC-XL-MS dataset. For each candidate trimer, AF3x (*13*) input files were generated for each candidate trimer from the corresponding residue-pair connections. Because AF3x allows each residue to participate in only one cross-link (whereas in XL-MS data a single residue can cross-link to several partners), we sampled three independent random subsets of cross-links per prediction, in each of which every cross-linked residue appears only once. Trimeric complexes exceeding the practical AlphaFold3 sequence-length limit (>2,500 residues on our GPU) were split into pairwise dimers before prediction; all such dimers fell within this limit.

Predicted trimeric structures were converted to standardized PDB files with consistent chain identifiers, and confidence metrics, including residue-level pLDDT values and residue-pair PAE matrices, were extracted from the AF3x output. Trimer predictions were then decomposed into pairwise interfaces. When multiple models were available for the same dimer, the representative structure was chosen as the model with the lowest mean Cα RMSD to all other predictions after pairwise structural superposition.

#### Monte Carlo tree search assembly

Higher-order protein complexes were assembled from predicted dimeric subcomplexes using a Monte Carlo tree search (MCTS)-based framework adapted from MoLPC (*32*). Pairwise interactions were represented as an interaction graph in which nodes corresponded to protein subunits and edges represented binary interactions supported by experimental XL-MS PPIs. During assembly, MCTS tree nodes represented partial assembly states rather than individual proteins, each storing the set of incorporated chains and their three-dimensional coordinates.

Whereas the original MoLPC implementation initializes assembly from a fixed reference chain, here independent MCTS searches were initiated from every subunit to avoid bias toward a predefined assembly order. The highest-scoring complete assembly across all searches was retained as the final structural model.

Starting from a root chain, tree traversal followed the standard upper confidence bound (UCB) criterion with an exploration constant of 2,

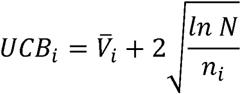

where V □_i is the mean rollout score of node i, N is the visit count of its parent node, and n_i is the visit count of node i. During expansion, all interaction edges connecting the current assembly to unincorporated subunits were considered candidate actions. The corresponding predicted dimer was docked by rigid-body superposition of the shared chain using the Kabsch algorithm, and the resulting transformation was applied to the incoming chain. Candidate assemblies were discarded when severe steric clashes were detected (>50% of the Cα atoms of the smaller chain located within 5 Å of the existing assembly).

Expanded nodes were evaluated by stochastic rollouts in which additional chains were randomly incorporated until either a complete assembly was generated or no further valid expansions remained. Rollout scores were back-propagated to all ancestor nodes and used to update subsequent tree traversal.

#### Assembly scoring (back-propagation score)

Each partial or complete assembly was evaluated using a composite score combining structural confidence and agreement with experimental XL-MS data,

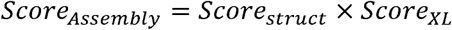

The structural score was calculated from the number of inter-chain interface contacts (Cβ–Cβ distance ≤8 Å; Cα for glycine; the same contact criterion as in the original MoLPC framework), weighted by the mean interface pLDDT:

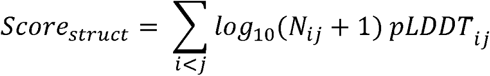

where Nij denotes the number of interface contacts between chains i and j, and *pLDDT_ij_* represents the mean pLDDT of the corresponding interface region. Cross-link agreement was quantified as the fraction of experimentally observed inter-chain cross-links whose Cα–Cα distance satisfied a cross-linker-specific threshold L:

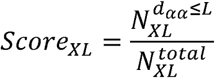

During MCTS-based assembly scoring, a threshold of L = 35 Å was used for DSBSO cross-links to improve discrimination among candidate assemblies. For final structural evaluation, cross-link satisfaction was assessed separately using a 40 Å Cα–Cα threshold. The composite score therefore favors assemblies with extensive, high-confidence interfaces that are simultaneously consistent with the XL-MS restraints.

#### Final scoring and selection

Each independent MCTS search (one per starting subunit) produced a complete assembly with an associated assembly score. The assembly with the highest assembly score across all starting points was selected as the final model.

### Biochemistry and imaging experiments

#### Cell culture of HeLa WT and LRBAKO1 cells

HeLa WT and LRBAKO1 cells (obtained from Absea biotechnology) were cultured in Dulbecco’s Modified Eagle Medium (DMEM; Gibco) supplemented with 10% fetal bovine serum (FBS) at 37 °C in a humidified incubator with 10% CO□.

#### siRNA-mediated gene silencing

HeLa WT cells were seeded in 12-well plates one day before transfection. The following siRNAs were used in this study: ON-TARGETplus Non-targeting siRNA (Dharmacon, catalog #D-001810-10-05) and ON-TARGETplus siRNAs against LRBA, FKBP15 and SDCCAG3 (Dharmacon, catalog #L-012751-00-0005, #L-029587-01-0005 and #L-016427-00-0005, respectively). Cells were transfected using Lipofectamine 2000 (Invitrogen) according to the manufacturer’s instructions. Briefly, siRNA (25 or 50 pmol per well) and 1 µL Lipofectamine 2000 were diluted separately in Opti-MEM (Gibco), combined, and incubated for 20 min at room temperature before being added dropwise to the cells. After 4 h, the transfection medium was replaced with fresh complete DMEM containing 10% FBS. Forty-eight hours after transfection, cells were harvested and processed for immunofluorescence or SP3 digestion.

#### Membrane fraction isolation

Cells were harvested, washed three times with PBS, and resuspended in starting buffer (225 mM mannitol, 75 mM sucrose, 30 mM Tris-HCl, pH 7.4) supplemented with complete EDTA-free protease inhibitor cocktail. Cells were homogenized at 1,000 rpm until 80–90% were broken. The homogenate was centrifuged three times at 10,000 × g for 10 min at 4 °C, discarding the pellet each time; the supernatant was then centrifuged at 25,000 × g for 20 min at 4 °C to pellet the PM (plasma membrane) and PAM (plasma membrane-associated membranes). The resulting supernatant was centrifuged at 95,000 × g for 1 h at 4 °C to pellet the microsomal fraction. The pellets were stored at -20 °C overnight. The next day, the PM/PAM and microsomal pellets were resuspended in 4% SDS and combined. Protein concentration was determined with the BCA protein assay kit (Thermo Fisher Scientific) according to the manufacturer’s instructions. Samples were then processed for western blot analysis or digested using the SP3 protocol.

#### Western blot analysis

Protein samples were separated by SDS-PAGE and wet-transferred to a 0.45 µm Immobilon-PSQ PVDF membrane (Millipore) at 110 V for 90 min. Blots were blocked with 3% BSA for 30 min at room temperature and incubated with the corresponding primary antibody at 4 °C overnight. The following antibodies were used in this study: anti-LRBA (1:500, rabbit; Sigma, HPA019366); anti-STX7 (1:1000, sheep; R&D Systems, AF5478); anti-β-actin (1:1000, mouse; Sigma, A5441). After washing three times with TBST, blots were incubated with an HRP-conjugated secondary antibody (goat anti-rabbit, goat anti-mouse or rat anti-sheep IgG) for 1 h at room temperature. After washing three times with TBST, blots were developed with SuperSignal West Pico PLUS chemiluminescent substrate (Thermo Fisher Scientific) and imaged on a ChemiDoc MP imager (Bio-Rad).

#### Confocal immunofluorescence

HeLa WT cells subjected to siRNA-mediated knockdown were fixed 48 h after transfection. Untreated HeLa WT and LRBA-KO1 cells were fixed 1–2 days after seeding, once they had reached the desired confluence. Cells were fixed with fixation buffer containing 4% paraformaldehyde and 4% sucrose for 15 min at room temperature. Following three washes with PBS, cells were permeabilized with PBS containing 0.1% Triton X-100 for 10 min, washed once with PBS, and blocked in PBS containing 1% BSA for 30 min at room temperature. Cells were then incubated with primary antibodies diluted in blocking buffer overnight at 4 °C in a humidified chamber. The following primary antibodies were used: anti-LRBA (1:100, rabbit; Sigma, HPA019366); anti-STX7 (1:100, sheep; R&D Systems, AF5478); anti-EEA1 (1:100, mouse; BD Transduction, 610456); anti-TGN46 (1:100, rabbit; Abcam, ab50595); anti-GM130 (1:100, mouse; BD Transduction, 610822); anti-CALR (1:100, rabbit; Abcam, ab92516); anti-LAMP1 (1:100, rabbit; Cell Signaling Technology, D2D11); and anti-WASHC3 (1:100, rabbit; Sigma, HPA038339). After three washes with PBS, cells were incubated with the secondary antibodies for 1 h at room temperature. Following three washes with PBS, coverslips were mounted with ROTI Mount FluorCare DAPI mounting medium (Roth) and imaged on an LSM710 confocal microscope (Zeiss).

For CALR staining, cells were fixed with prewarmed (37 °C) fixation buffer containing 4% paraformaldehyde and 4% sucrose for 30 min at 37 °C. Following three washes with PBS, cells were incubated with 3% BSA in 0.3% PBST (PBS containing 0.3% Triton X-100) for 30 min. In the same buffer, cells were incubated with primary antibodies for 2 h at room temperature, washed three times with 0.3% PBST, and then incubated with secondary antibodies for 1 h at room temperature.

#### Image quantification

For each experiment, approximately 200 cells per condition were quantified from 3 × 3 tile scans. Cells were segmented using a custom Fiji macro to enable per-cell quantification. Pearson’s correlation coefficients were calculated using the JACoP plugin in Fiji. For puncta intensity measurements, a fixed intensity threshold was applied prior to quantification. For vesicle quantification, images were TopHat-filtered for particle detection, and vesicle size, number, mean intensity and integrated intensity were measured using the Analyze Particles function in Fiji. Identical image-processing and thresholding parameters were applied across all experimental groups within each experiment.

### Systems biology augmentation

#### Four-database comparison (DUC-XL-MS, EndoMAP, OpenCell, BioPlex)

Organelle annotation was performed as described in “Organelle marker assignment”, parsing the UniProtKB “Subcellular location [CC]” annotation into eight canonical compartments: nucleus, mitochondrion, endoplasmic reticulum (ER), Golgi apparatus, endosome, lysosome, peroxisome and plasma membrane; the ER–Golgi intermediate compartment (ERGIC) was assigned to both ER and Golgi. Multi-localized proteins were retained in each of their compartments, and proteins annotated to more than three canonical compartments were excluded. Proteins of classes (i)–(iv) defined in “Exclusion of promiscuously cross-linked proteins” were removed. Dataset-specific hub proteins, defined per dataset as proteins contributing more than 1% of that dataset’s interaction edge ends, were also removed: RPS27A, HSPA1A and NPM1 in DUC-XL-MS; EEA1, the immunoisolation target, and HSP90AB1 in EndoMAP; and CAPZB in OpenCell; no BioPlex protein met the threshold. All interactions of an excluded protein were discarded, so an interaction was retained only when both partners passed all filters.

#### Triadic-closure-based augmentation of the DUC-XL-MS interactome

DUC-XL-MS interactome was augmented with interactions reported from three complementary resources: BioPlex 3.0 (AP–MS), OpenCell (native organelle IP–MS) and EndoMAP (XL-MS). The filtered data (see section *Four-database comparison (DUC-XL-MS, EndoMAP, OpenCell, BioPlex)* were used for augmentation. Each candidate pair was retained if it was directly detected by DUC-XL-MS or, if not directly observed, if the two proteins shared at least one neighbors in the DUC-XL-MS network, adapting the mutual clustering criterion (*53*). The resulting augmented interactome combined experimentally detected XL-MS interactions with topology-supported associations, providing expanded coverage of the cellular interactome.

#### PDB structure mapping

Cross-linked residue pairs from the pooled HEK293 and Jurkat DUC-XL-MS datasets were mapped systematically across the PDB. Each UniProt accession carrying a cross-link was mapped through SIFTS (*54*) to all annotated PDB chains, identifying 113,406 of the 222,787 pooled residue pairs (50.9%) for which both partner proteins were present in at least one deposited structure. To reduce redundancy, PDB entries were grouped by protein composition and a greedy set-cover algorithm selected 2,000 representative structures, together covering 111,339 of the mappable pairs (98.2%). Cross-link positions were required to be lysines in the canonical UniProt sequence, and per-protein numbering offsets were corrected against that sequence. Each deposited chain was then checked against the same canonical sequence and renumbered where necessary. Sites and chains that could not be reconciled were discarded.

For each retained structure, Cα–Cα distances were calculated for all mapped cross-linked residue pairs, taking the minimum distance across all compatible chain-copy pairs in homo-oligomeric structures. Same-residue self-links were excluded, as they cannot be intramolecular. When a residue pair was represented in multiple retained structures, the minimum distance across all structures was used, so that a cross-link was counted as satisfied if any deposited conformational state placed the two residues within 40 Å, the maximum Cα–Cα distance compatible with the DSBSO spacer arm.

Conformationally heterogeneous structures were removed from the distance analysis using a criterion applied uniformly to the pooled HEK293 and Jurkat measurements: a PDB entry contributing ≥200 cross-linked residue-pair measurements was excluded when more than 30% of the measurements on that structure exceeded 40 Å. Nine structures met this criterion: five ribosome-associated assemblies (PDB 4V6X, 9PA7, 6ZMO, 8G6J and 9I2E), mitochondrial Hsp60–Hsp10 (9SHG), mitochondrial Hsp70 (mortalin)–GrpE1 (9QIN), Ku70/80–WRN (9HZG) and a kinesin-1 heterotetramer (9PMB). Of the 2,000 selected structures, 1,618 yielded measurable cross-links and 1,609 were retained after this exclusion.

## Data availability

The mass spectrometry proteomics data have been deposited to the ProteomeXchange Consortium via the PRIDE partner repository with the dataset identifier PXD081158 (HEK) and PXD081113 (Jurkat). Search results are presented in Table S1. Source data are provided with this paper.

## Acknowledgements

Funding was provided by the Leibniz Association (Leibniz-Wettbewerb, P70/2018) and the European Research Council (ERC-2020-StG, project no. 949184) to F.L.; by the Deutsche Forschungsgemeinschaft (DFG, HA2686/24-1) and the Leibniz Prize (HA2686/25-1) to V.H.; and by the China Scholarship Council (no. 202304910020) to Z.Z. Computational resources were granted by the Resource Allocation Board and provided on the supercomputer Emmy/Grete at NHR-Nord@Göttingen as part of the NHR infrastructure under the project beb00048. The authors thank Philip Lössl (Absea Biotechnology, Berlin) and Jan Kosinski (EMBL, Hamburg) for critically reviewing and editing the manuscript.

## Author Contributions

F.L. conceptualized the project. N.Y. and Y.Z. performed the experiments. Z.Z., F.L. and N.Y. performed data analysis. Z.Z. and F.L. curated the data. V.N., J.C. and T.S. contributed to experiments and data analysis. J.R. predicted the AF models for the SLiM analysis. V.H. and F.L. procured resources and supervised the research. F.L. wrote the manuscript. Z.Z., N.Y. and V.H. contributed to the writing. All authors reviewed and edited the manuscript.

## Competing interests

F.L. is a shareholder of Absea Biotechnology and Proxima. Y. Z is an employee at Absea Biotechnology. The remaining authors declare no competing interests.

