## Supplementary Figures for "Site-resolved spatial and structural interactome of a human cell"

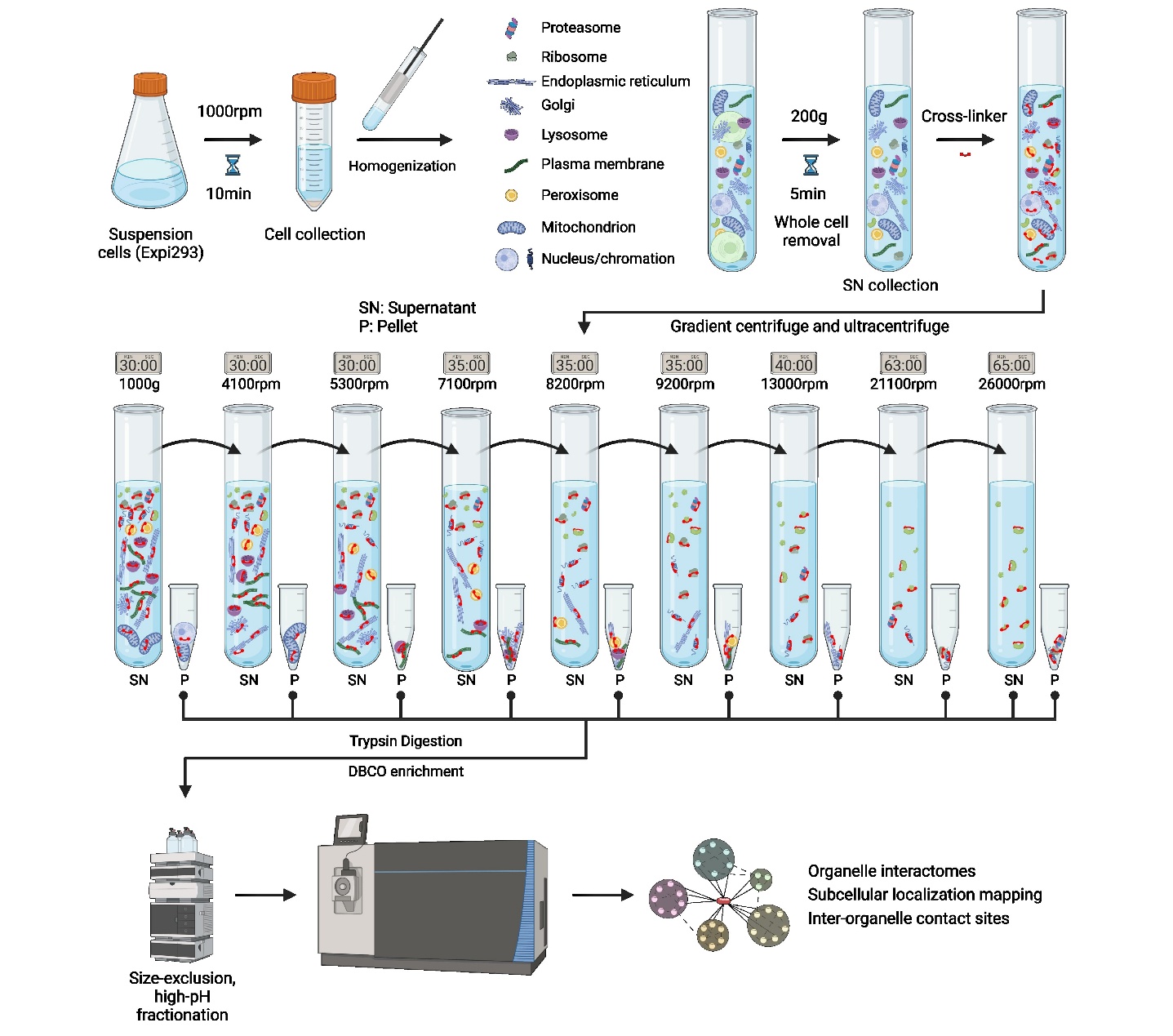

**Fig. S1 | Overview of the differential ultracentrifugation cross-linking mass spectrometry (DUC-XL-MS) workflow.** HEK293F cells grown in suspension are harvested (1,000 rpm, 10 min) and homogenized, and whole cells and large debris are removed by low-speed centrifugation (200 × g, 5 min). The cleared homogenate is cross-linked with DSBSO and fractionated by a series of differential centrifugation and ultracentrifugation steps at increasing speeds, generating organelle-enriched pellets (P) and successive supernatants (SN). All fractions are digested with trypsin; cross-linked peptides are enriched on DBCO beads, separated by size-exclusion chromatography and analyzed by LC–MS/MS, yielding a proteome-wide interaction map that includes inter-organelle contact sites. Image is created with BioRender.

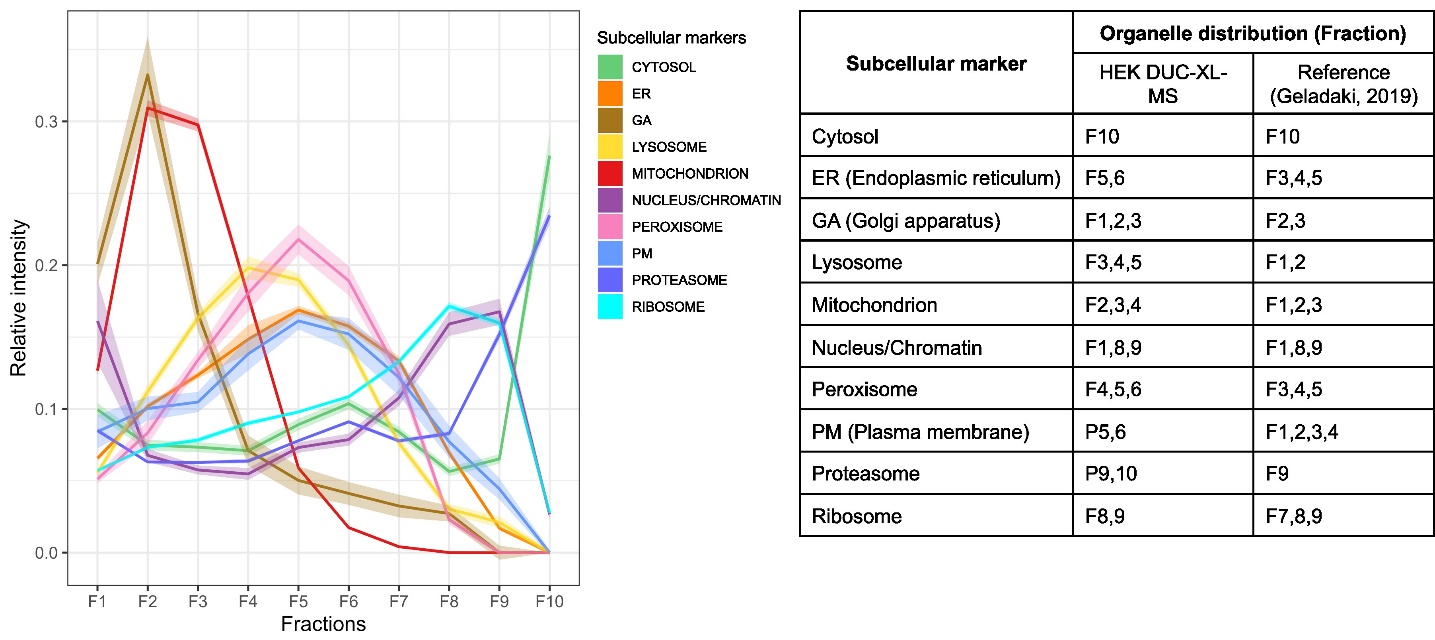

**Fig. S2 | Organelle distribution across DUC fractions.** (Left) Median organelle-marker profiles along all subcellular gradient fractions of the HEK DUC-XL-MS sample; shaded regions denote ± s.e.m. (Right) Summary table comparing DUC-XL-MS with the original LOPIT-DC study (*1*); enriched fractions (relative intensity > 0.15) are shown.

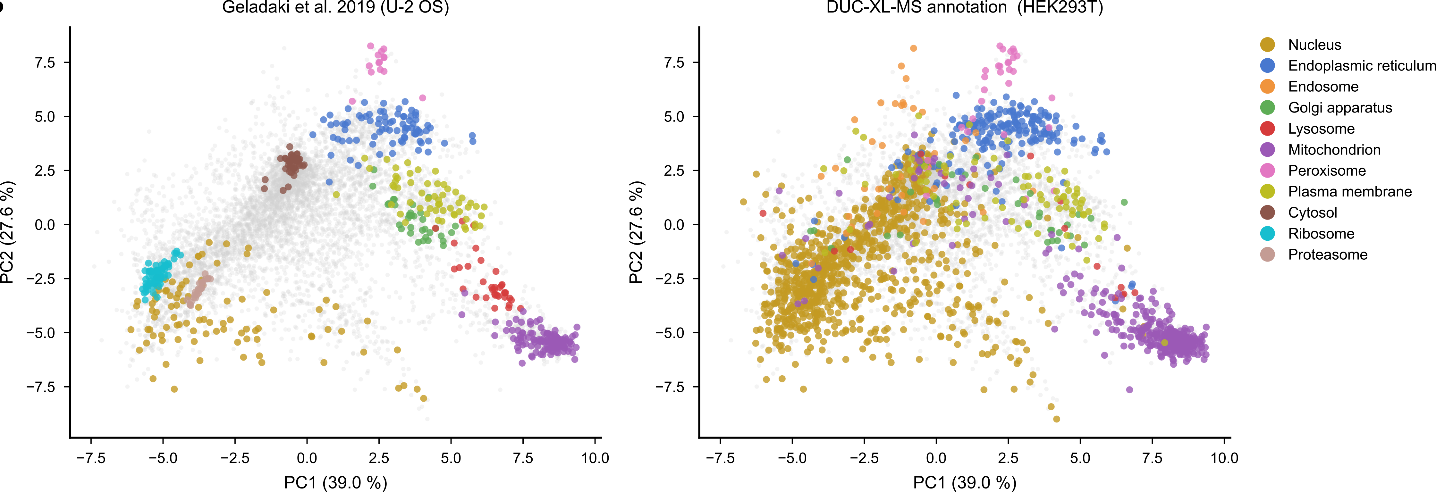

**Fig. S3 | DUC-XL-MS spatial annotations compared with LOPIT-DC reference localizations.** PCA coordinates were taken from the published LOPIT-DC fractionation analysis of U-2 OS cells (*1*); each point is one protein, and both panels use identical coordinates colored by two independent annotations. Left, curated LOPIT-DC subcellular localizations (Geladaki et al. (*1*), Fig. 2). Right, DUC-XL-MS graph-propagation annotations (Table S3), comprising single-compartment assignments supported by organelle marker proteins or by propagation; proteins that were unannotated or assigned to multiple compartments are shown in gray.

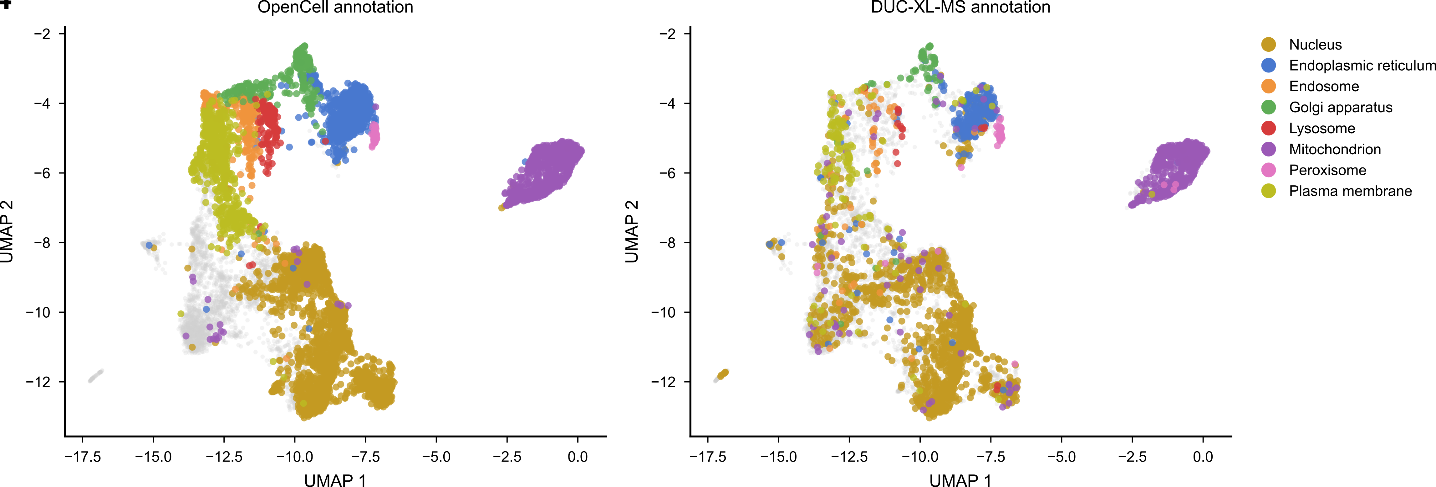

**Fig. S4 | DUC-XL-MS spatial annotations compared with OpenCell UMAP localizations.** UMAP coordinates were taken from the published OpenCell embedding of HEK293T cells (*2*); each point is one protein, and both panels use identical coordinates colored by two independent annotations. Left, OpenCell subcellular localizations (adapted from Cho et al. (*2*), Fig. 3D). Right, DUC-XL-MS graph-propagation annotations (Table S3), comprising single-compartment assignments supported by organelle marker proteins or by propagation; proteins that were unannotated or assigned to multiple compartments are shown in gray.

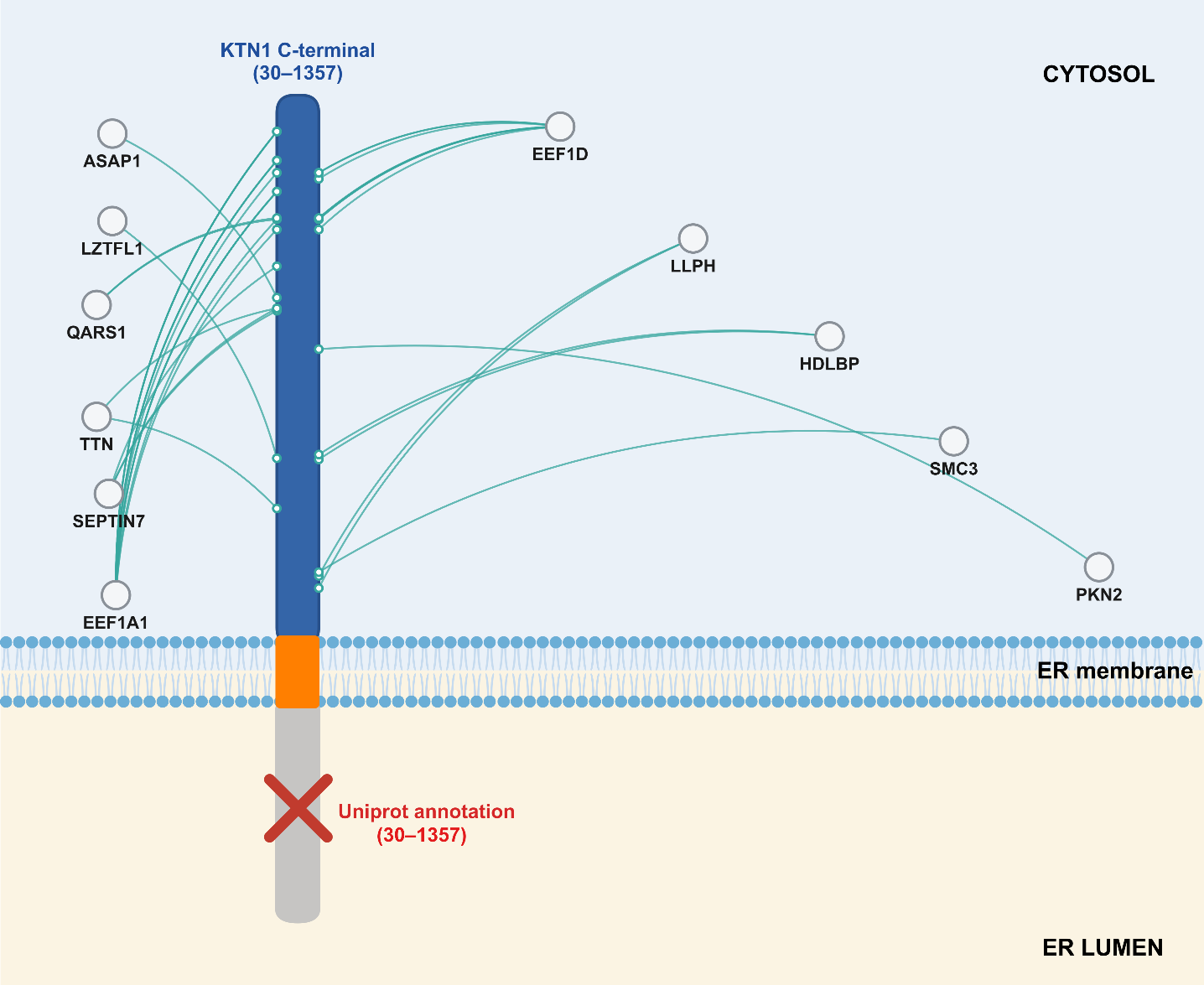

**Fig. S5 | Cross-links correct the annotated membrane topology of KTN1**. Schematic of the ER membrane protein kinectin-1 (KTN1, Q86UP2), comprising a UniProt-predicted type II signal anchor (residues 7–29, dashed gray) and the C-terminal coiled-coil region (residues 30–1357, blue). UniProt annotates residues 30–1357 as a lumenal topological domain; 36 cross-links to 11 cytosolic proteins instead place this region in the cytosol, contradicting that assignment (red cross). Numbers above each node give the cross-links to that partner.

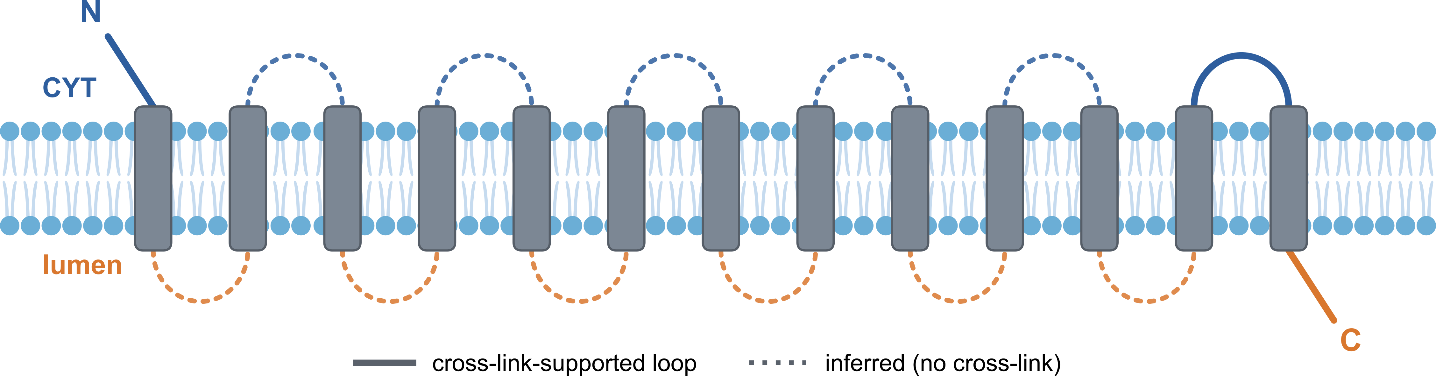

**Fig. S6 | Cross-link-resolved membrane topology of STT3B.** Schematic of the 13 transmembrane helices (gray) of the ER oligosaccharyltransferase catalytic subunit STT3B (Q8TCJ2). Cross-links between transmembrane loops and soluble rulers assign the N-terminus and the internal loop L12 to the cytosol (blue) and the C-terminus to the ER lumen (orange), consistent with an alternating topology. Loops directly supported by cross-links are drawn solid (N-terminus, L12, C-terminus; 31 cross-link residue pairs in total, all agreeing with UniProt); the remaining loops, inferred from the alternating topology with no direct cross-link, are dotted.

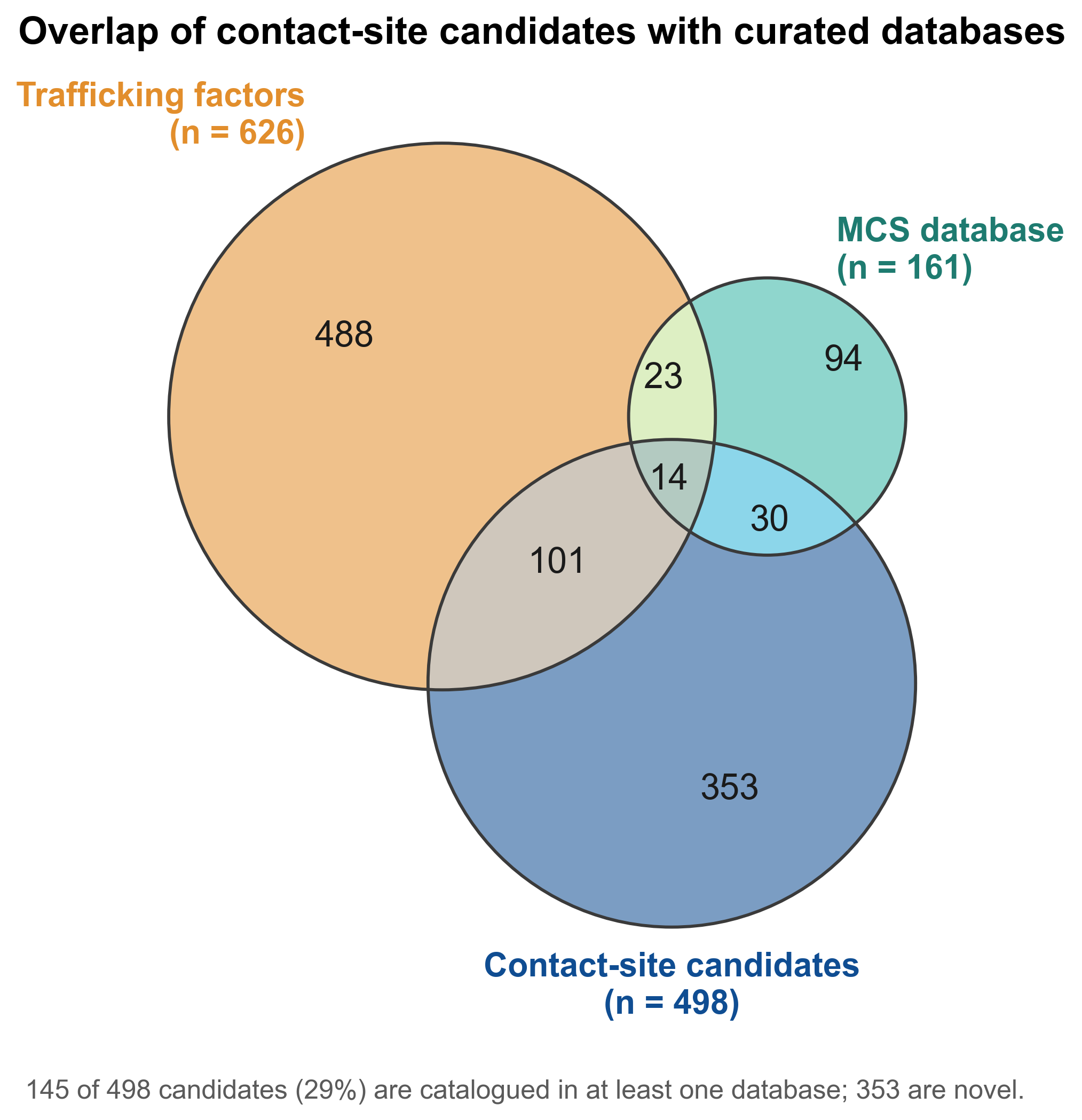

**Fig. S7 | Contact-site candidates overlap partially with other databases**. Three-way overlap of the cross-link-derived contact-site candidates (n = 498; dual-localized proteins with allowed-adjacency membrane compartments plus resolved-topology transmembrane proteins), the membrane-contact-site (MCS) database (n = 161) and curated trafficking-complex factors from CORUM (n = 626).

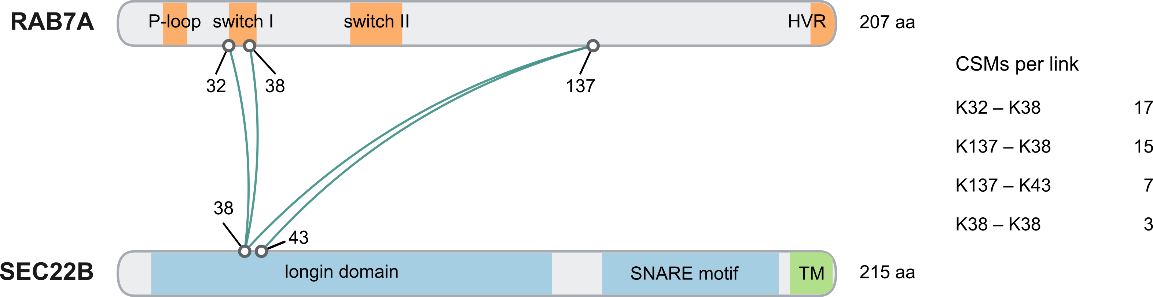

**Fig. S8 | Residue-level cross-links between RAB7A and SEC22B.** Cross-link map of RAB7A (late-endosomal/lysosomal Rab GTPase; orange) and SEC22B (ER/ERGIC longin R-SNARE; blue), drawn to scale with domain annotations.

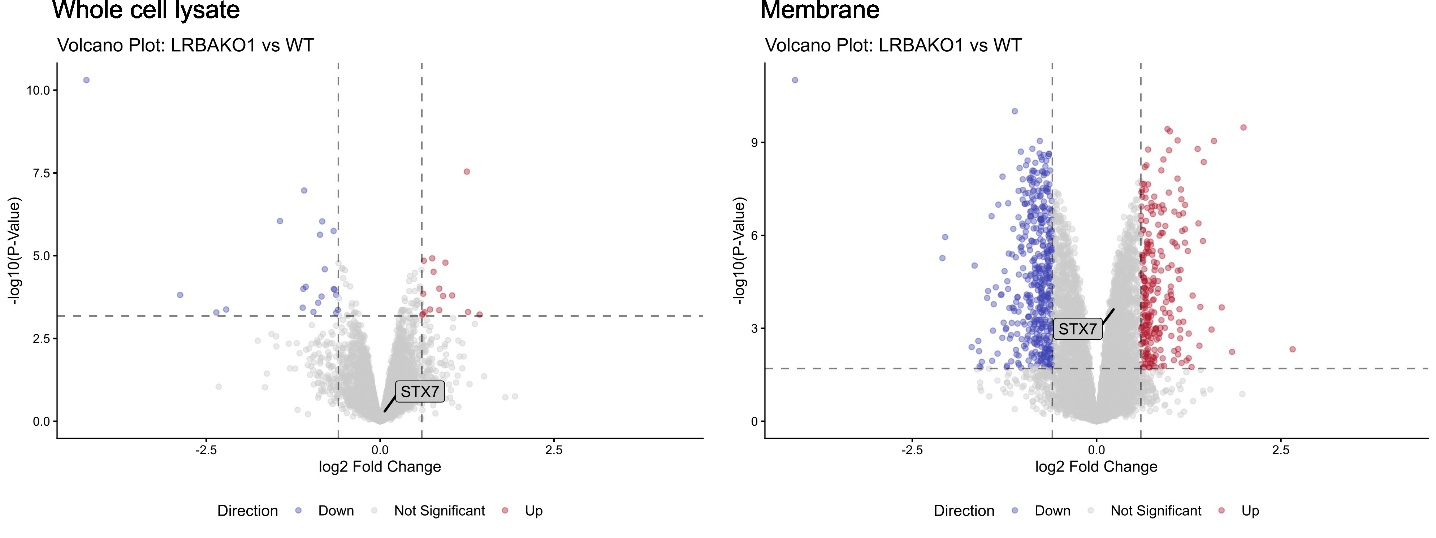

**Fig. S9 | Proteome analysis of LRBA-KO cells versus WT.** Volcano plots of protein abundance changes in LRBA-KO cells relative to WT cells for the whole-cell lysate (left) and the membrane fraction (right). Significance thresholds were an adjusted P value < 0.05 and |log₂ FC| > 0.6. STX7 abundance was unchanged in both fractions.

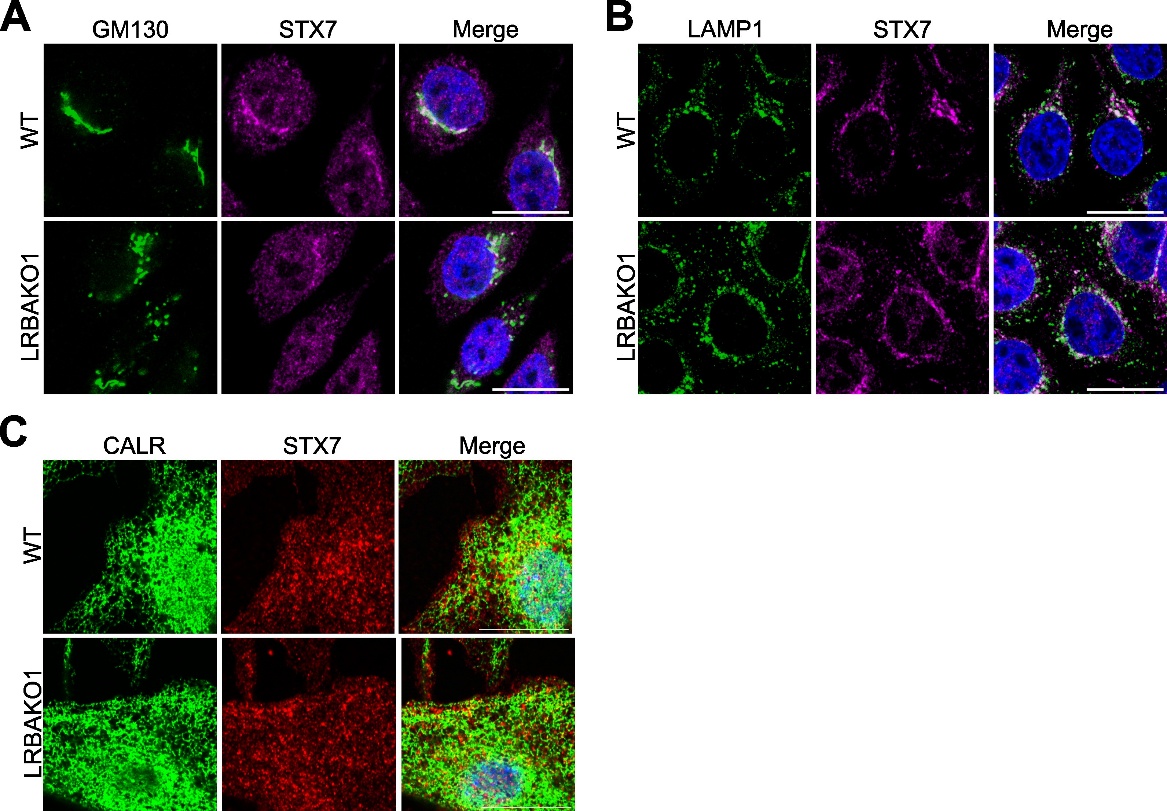

**Fig. S10 | Confocal images of STX7 with Golgi, lysosome and ER markers.** Confocal microscopy images of HeLa WT and LRBAKO1 cells co-stained with antibodies against STX7 and GM130, LAMP1, or CALR. Scale bars, 20 μm.

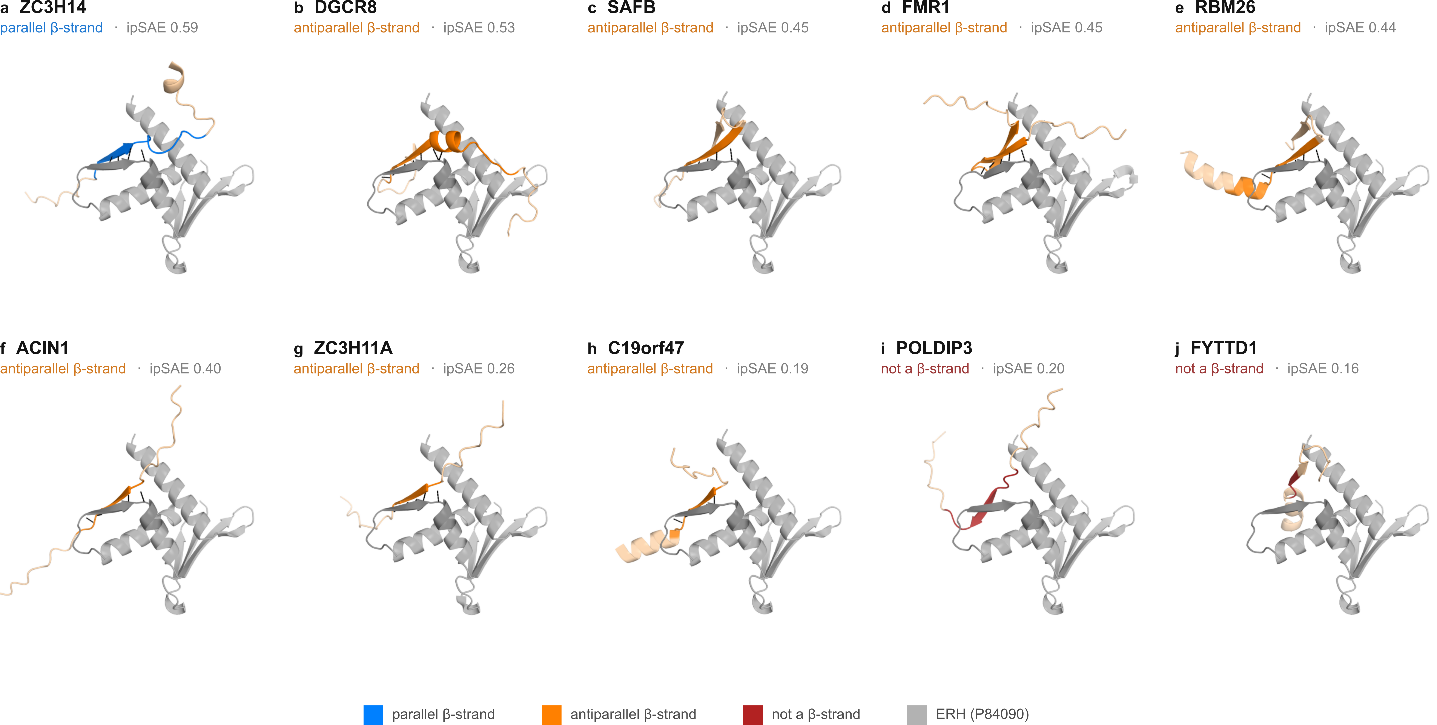

**Fig. S11 | ERH candidate SLiMs on the ERH β-sheet.** AF3 dimer models of ten candidate SLiM partners, all superposed on ERH (gray cartoon, edge strand 44–58 darker) and shown in the same orientation. Each SLiM core is colored by β-pairing geometry: parallel (blue), antiparallel (orange), or not β-paired (red), with ±10-residue flanks in wheat and backbone–backbone H-bonds to ERH as black dashes. Panels labelled with partner gene and ipSAE.

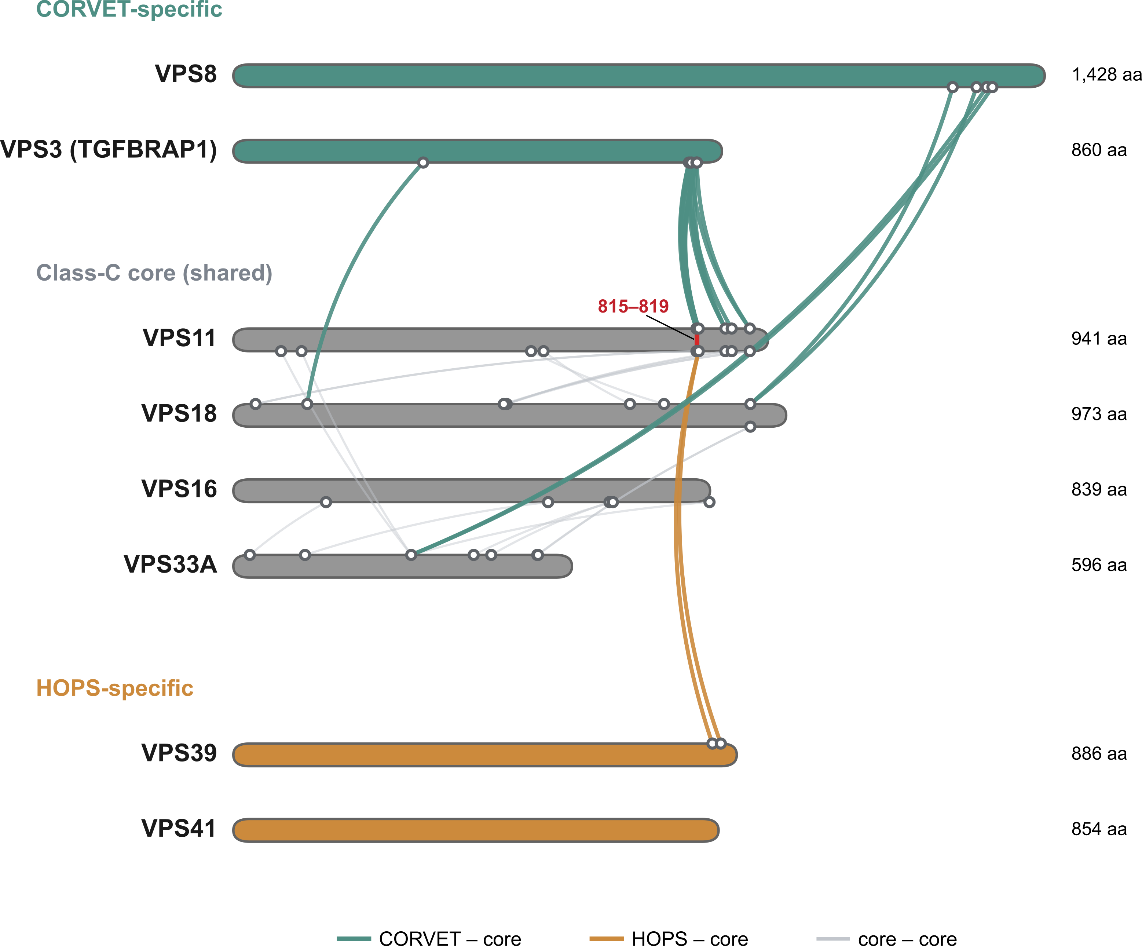

**Fig. S12 | A shared VPS11 docking site for the CORVET and HOPS-specific subunits.** Cross-link map of the human Class C core (VPS11, VPS18, VPS16, VPS33A; gray), the CORVET-specific subunits VPS8 and VPS3/TGFBRAP1 (teal), and the HOPS-specific subunits VPS39 and VPS41 (orange). Bars scale with protein length, with the residue number given at each C-terminus; ticks mark cross-linked residues and lines are cross-links, colored by whether they connect a CORVET-specific subunit to the core (teal), a HOPS-specific subunit to the core (orange) or two core subunits (gray). Cross-links from VPS3 and from VPS39 converge on the same region of VPS11 (residues ~815–819, red arrow), consistent with the two subunits docking mutually exclusively at this site and marking the point at which the CORVET and HOPS architectures diverge.

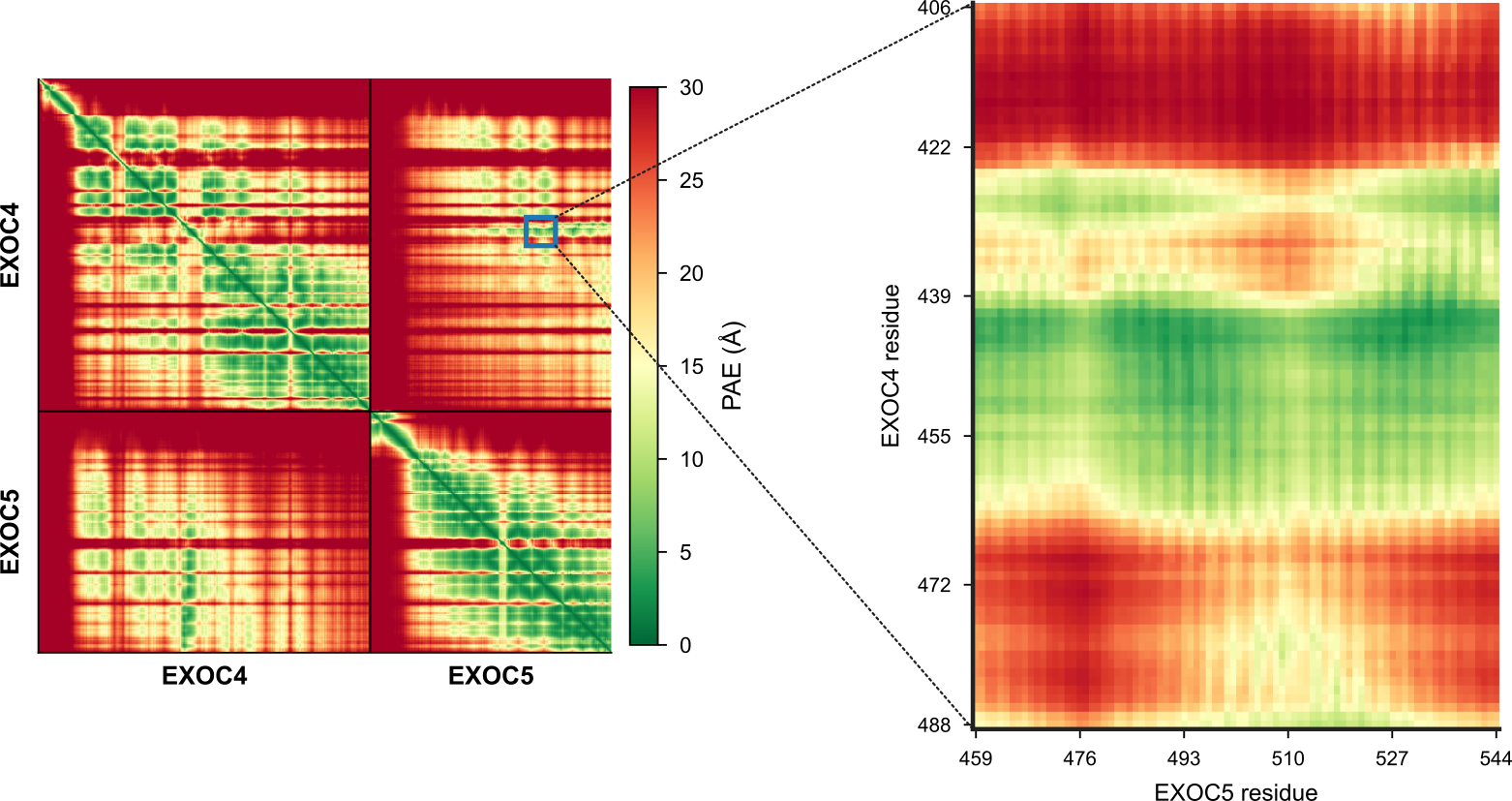

**Fig. S13 | PAE metrics for the exocyst EXOC4–EXOC5 interface.** (Left) Full predicted aligned error (PAE) matrix; the blue box marks the inter-subunit contact region, enlarged (right) over EXOC4 residues 406–488 × EXOC5 residues 459–544. A band of low PAE (green; centered on EXOC4 ~439–455) spans the EXOC5 residue range, indicating a well-defined, confidently predicted contact between the EXOC4 C-terminal loop and EXOC5.

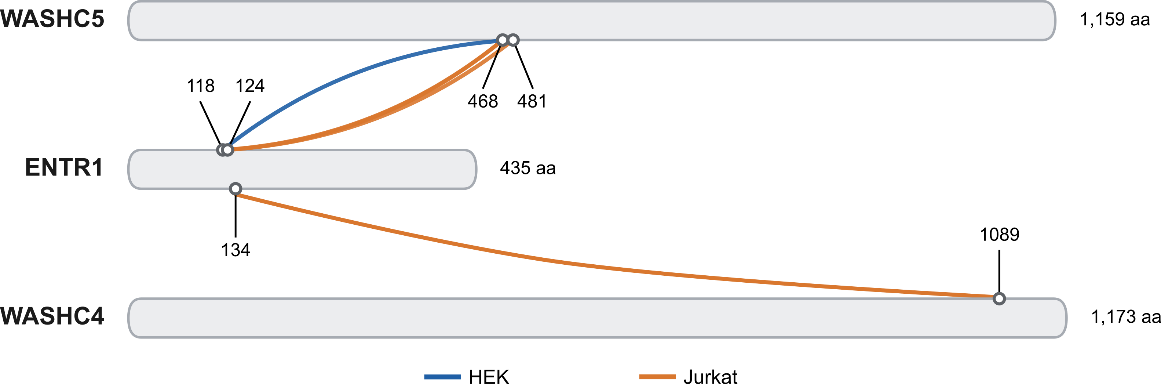

**Fig. S14 | ENTR1 cross-links to the WASH complex in HEK and Jurkat DUC-XL-MS data.** Residue-resolved cross-links from ENTR1 to the WASH-complex subunits WASHC5 and WASHC4; protein bars are drawn to scale and open circles mark cross-linked residues. Cross-links detected in the HEK network are shown in blue and those in the Jurkat network in orange (1 HEK and 4 Jurkat cross-links, 1 in both).

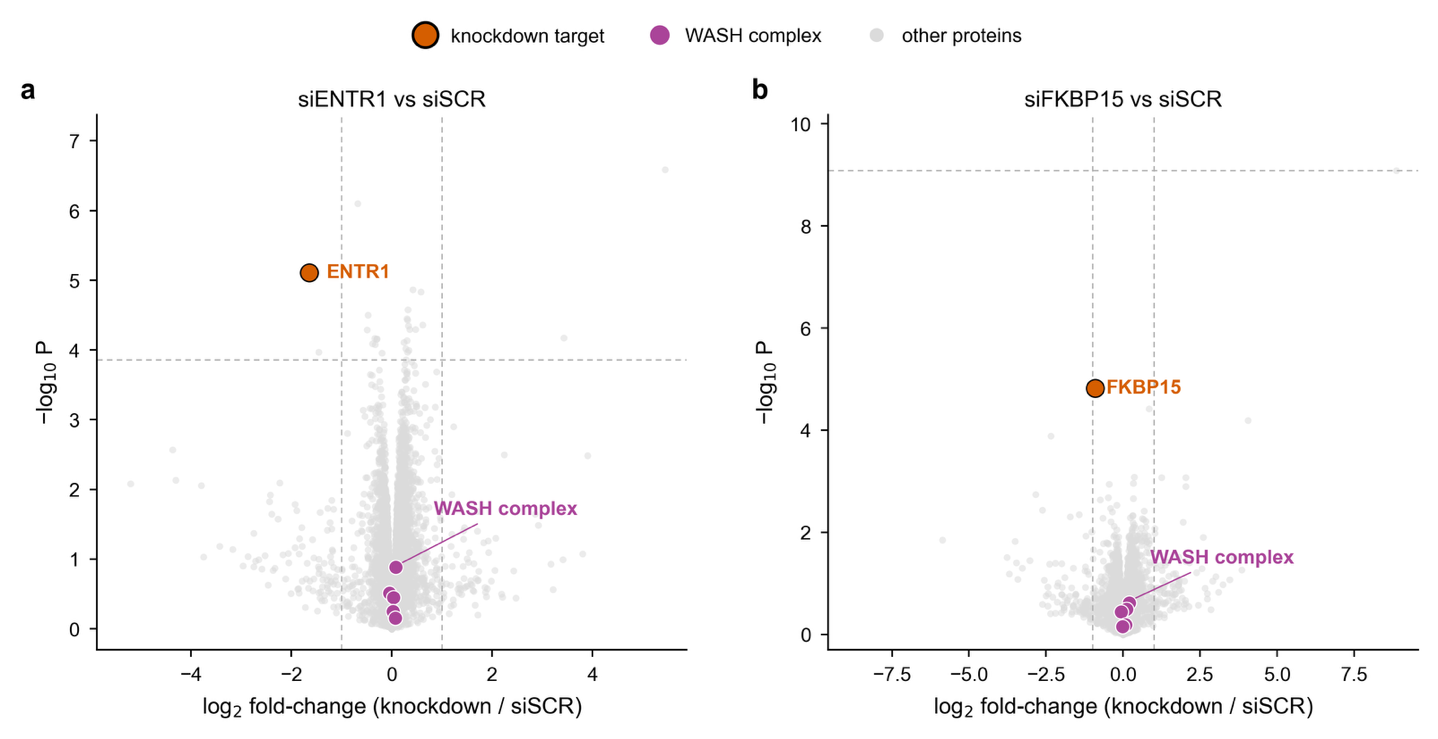

**Fig. S15 | Knockdown of ENTR1 or FKBP15 does not alter WASH-complex abundance.** Whole-proteome volcano plots after siRNA knockdown of (a) ENTR1 or (b) FKBP15 versus a non-targeting control (siSCR). Axes show log₂ fold change (knockdown/siSCR) and −log₁₀ P (limma moderated t-test, Benjamini–Hochberg FDR). The knockdown target (orange) and WASH-complex subunits (WASHC1–WASHC5; purple) are highlighted, all others gray. Dashed lines mark |log₂ FC| = 1 and adjusted P = 0.05. The target is specifically depleted in both experiments whereas WASH-complex subunits remain at log₂ FC ≈ 0 and non-significant, indicating that neither knockdown affects steady-state WASH-complex abundance.

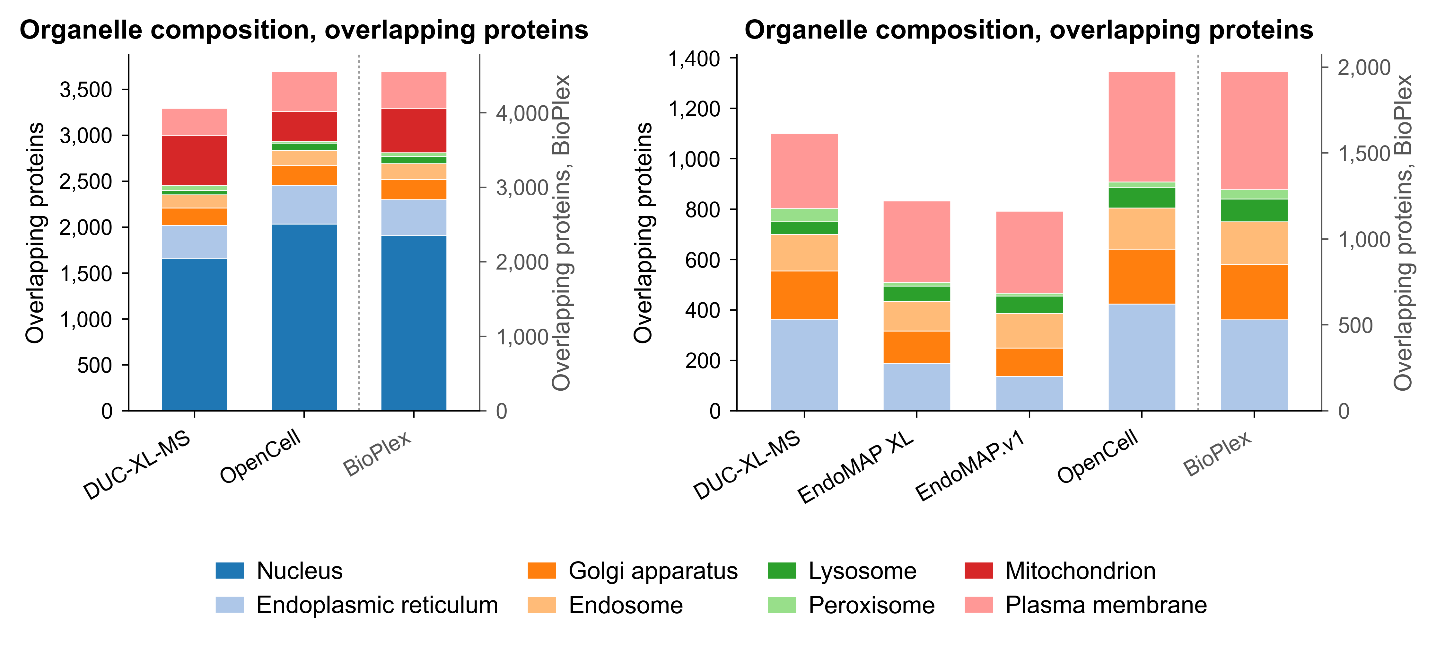

**Fig. S16 | Organelle composition of interactome proteins across datasets.** Stacked bar charts showing, for each dataset, the number of proteins assigned to eight subcellular compartments (left) or six compartments (nucleus and mitochondrion excluded; right). Only proteins shared with at least one other dataset of the same comparison are counted. Proteins are counted once per annotated compartment, so multi-localized proteins contribute to multiple compartments. Protein sets are derived from the filtered PPIs underlying Fig. 7A; BioPlex is plotted on a separate right-hand axis.

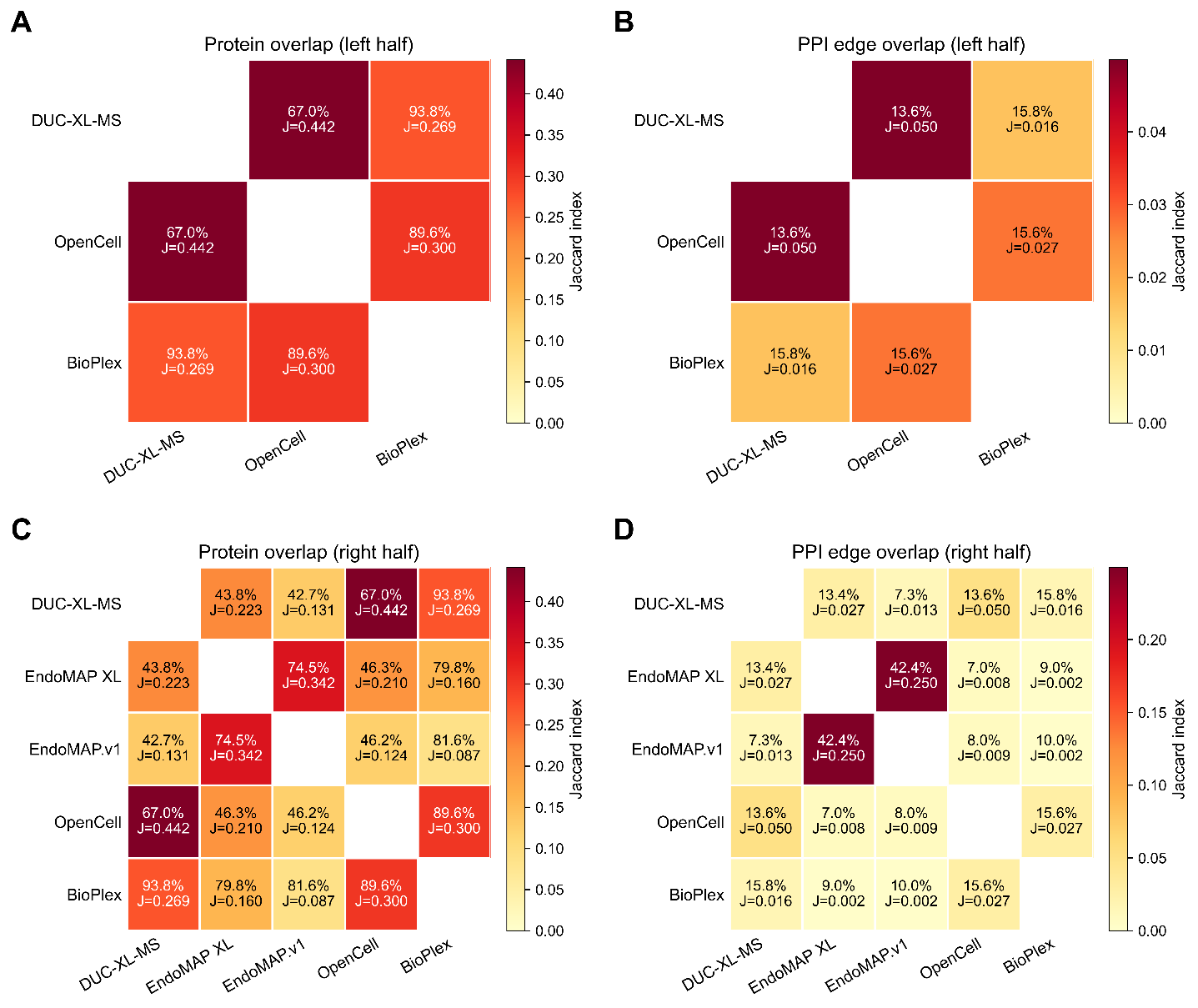

**Fig. S17 | Pairwise overlap between DUC-XL-MS and reference interactome datasets.** Protein-level (A, C) and interaction-level (B, D) overlap of the filtered datasets underlying Fig. 7A, for the three-dataset (A, B) and five-dataset (C, D) comparisons. Each off-diagonal cell reports the percentage of overlapping interactions, defined as ∣A∩B∣/min(∣A∣,∣B∣), together with the Jaccard index (J=∣A∩B∣/∣A∪B∣), where A and B are the sets of unique proteins (A, C) or unique PPIs (B, D) in each dataset. Cells are colored according to the Jaccard index. Datasets share 43–94% of proteins but only 7–16% of interactions (excluding the EndoMAP XL–EndoMAP.v1 pair).

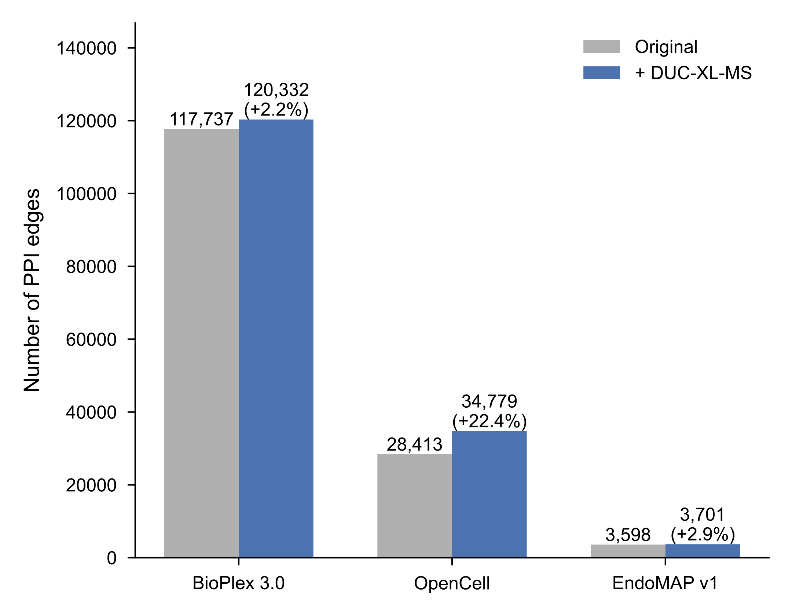

**Fig. S18 | DUC-XL-MS-supported augmentation of BioPlex 3.0, OpenCell and EndoMAP v1 PPI networks.** The number of PPIs before (gray) and after (blue) DUC-XL-MS augmentation. Labels indicate the total number of edges, with percentages showing the relative increase over the original networks.

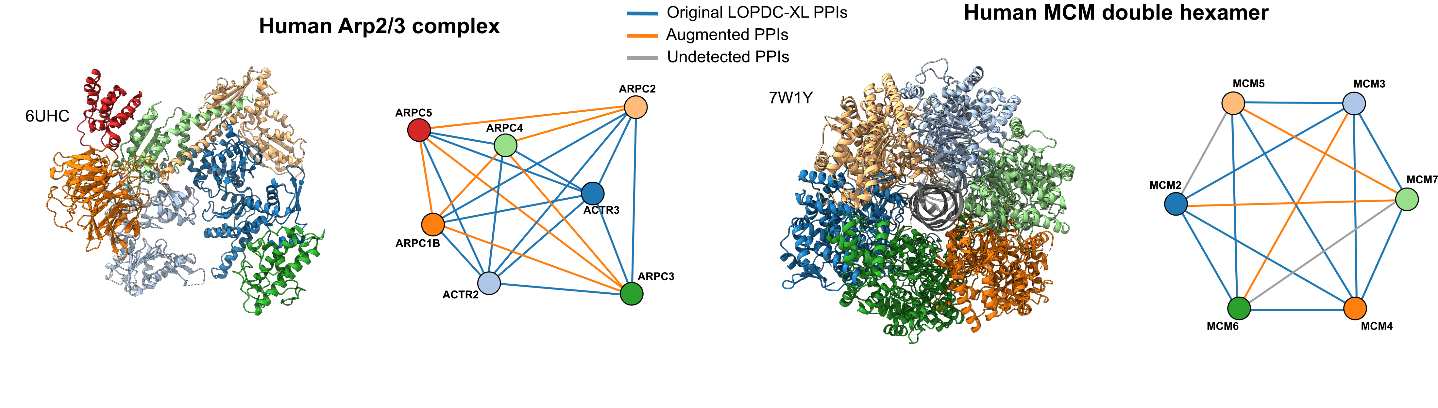

**Fig. S19 | PDB-based validation of augmented PPI edges (companion to Fig. 7F).** Two complexes whose subunits are all contained within a single deposited structure: human Arp2/3 complex (PDB 6UHC) and human MCM double hexamer (PDB 7W1Y). Left, structure with subunits individually colored. Right, graph of all pairwise subunit combinations, with edges colored by detection status: blue, present in the original DUC-XL-MS network; orange, added by triadic closure against OpenCell, BioPlex 3.0 and EndoMAP.v1; grey, recovered by neither.

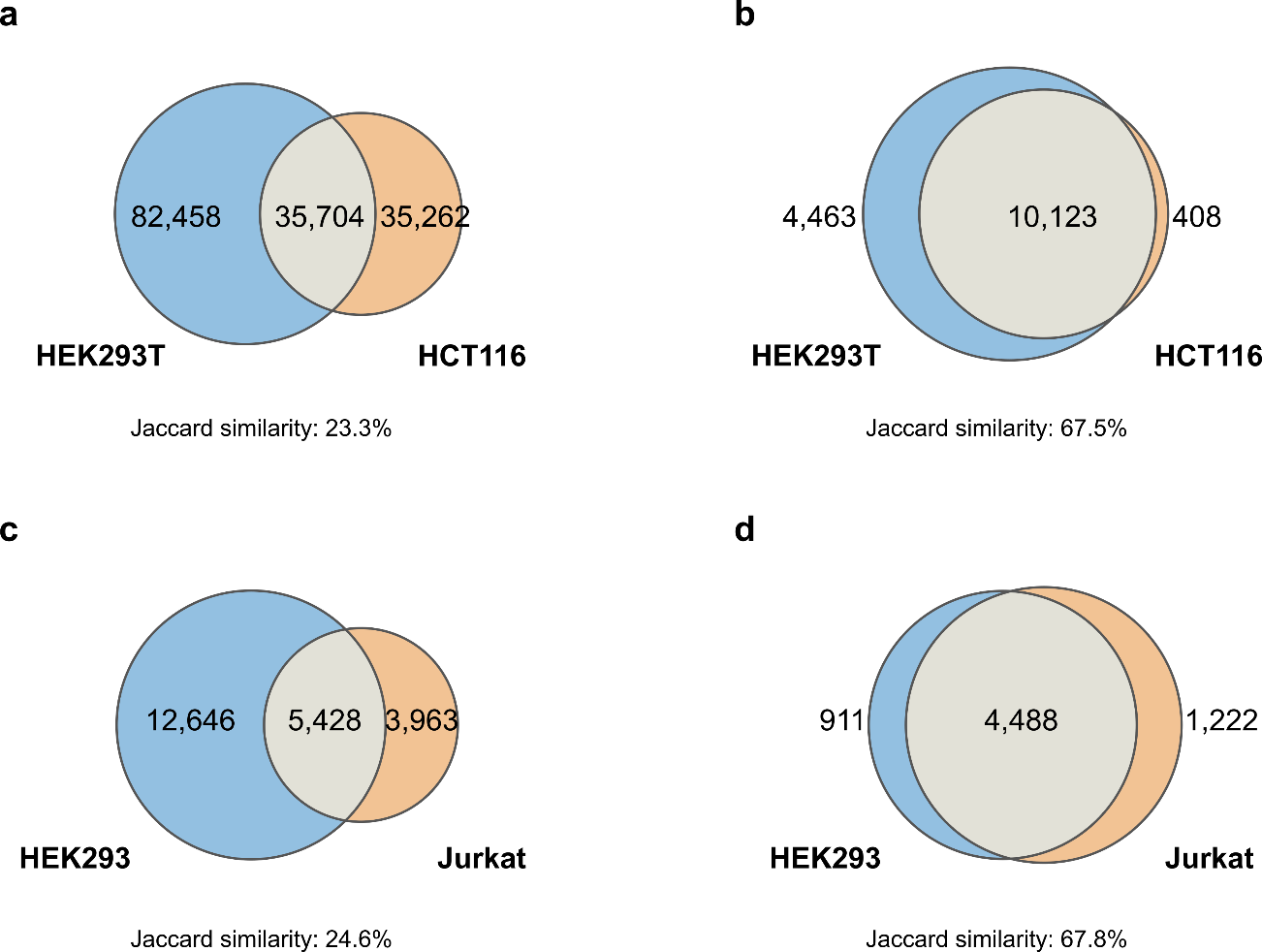

**Fig. S20 | Cross-cell-line reproducibility of interactomes and proteomes.** Venn diagrams of protein–protein interactions (PPIs; **a**, **c**) and proteins (**b**, **d**) identified in two cell lines by BioPlex 3.0 (**a**, **b**; HEK293T and HCT116) and DUC-XL-MS (**c**, **d**; HEK293 and Jurkat). Numbers indicate cell-line-specific and shared identifications. Interaction overlap is similar for both methods, whereas protein overlap is considerably higher than interaction overlap. Jaccard similarity is calculated as the intersection divided by the union of the two sets.

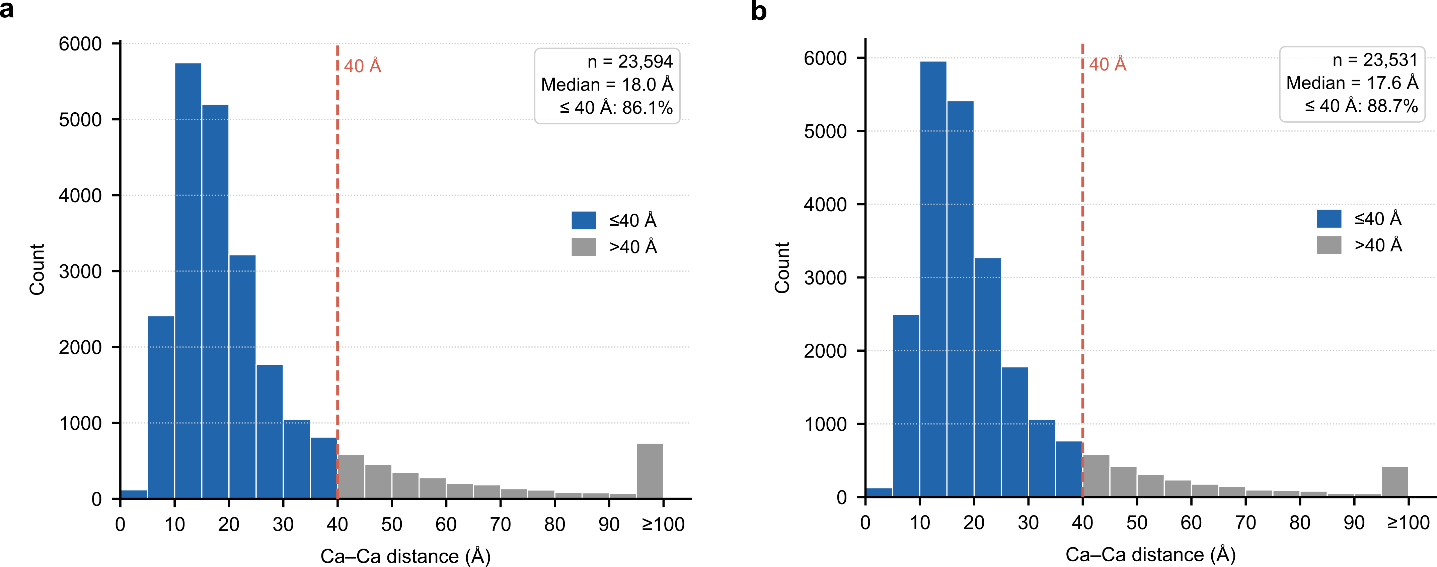

**Fig. S21 | Mapping of DUC-XL-MS cross-links onto PDB structures**. Cα–Cα distance distributions for cross-linked residue pairs mapped onto experimental structures in HEK293 and Jurkat cells. Left, HEK293: 23,594 cross-links across 1,469 PDB structures, 86.1% within the 40 Å Cα–Cα threshold. Right, Jurkat: 23,531 cross-links across 1,444 structures, 88.7% within the threshold.

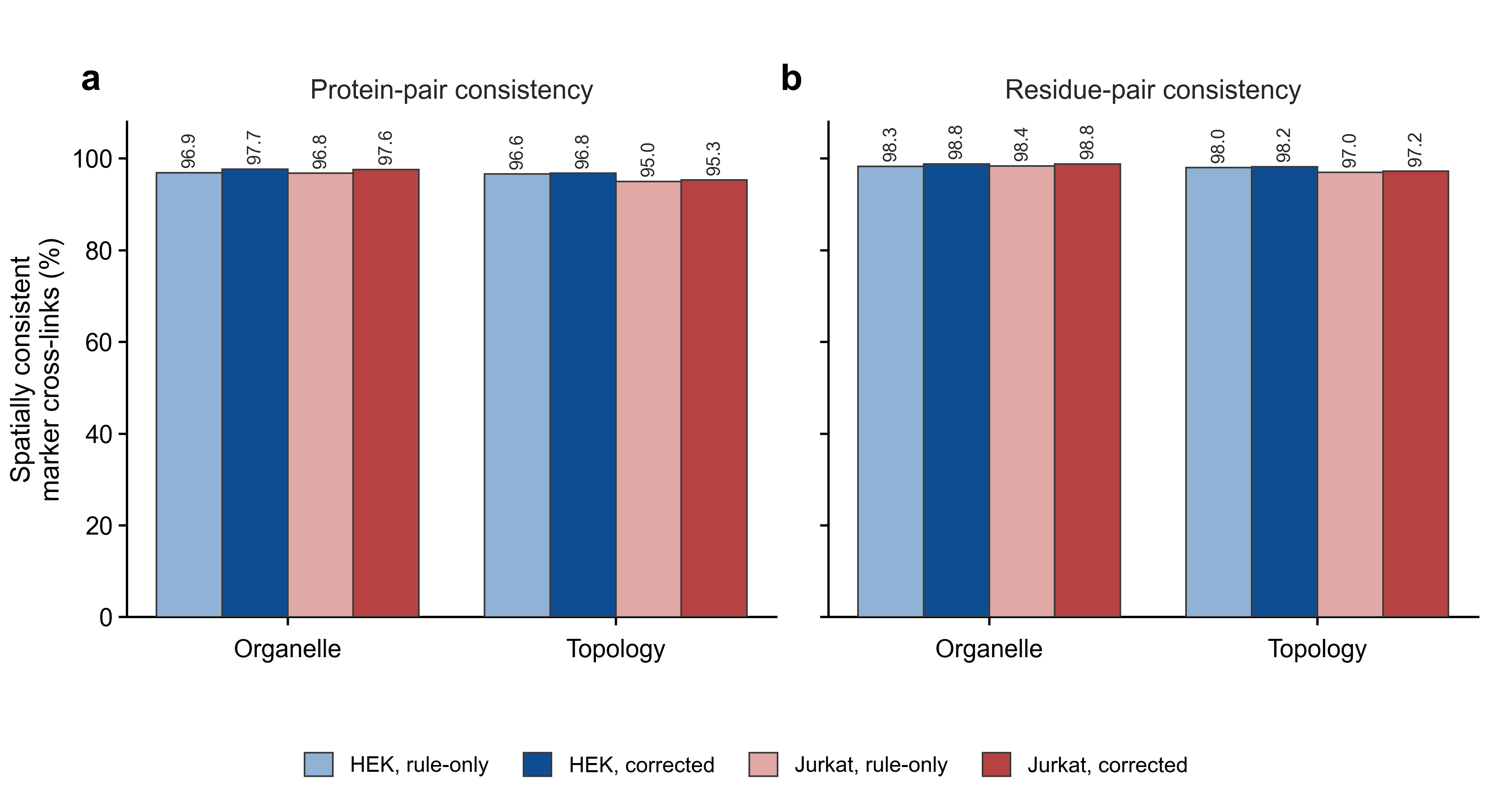

**Fig. S22 | HEK and Jurkat marker cross-link consistency is comparable across cell lines.** Organelle-level (a) and topology-level (b) consistency in the HEK and Jurkat DUC-XL-MS datasets, quantified as the percentage of marker cross-links that are spatially possible (possible connections divided by all connections), before (rule-only; lighter bars) and after (corrected; solid bars) manual curation.
