## Supplementary Notes for "Site-resolved spatial and structural interactome of a human cell"

**Supplementary Note 1: Cross-link spatial consistency of DUC-XL-MS, in-cell XL-MS and a random-network null**

***Rationale***

To test whether disrupting the cell membrane and cross-linking on cell lysate still preserves spatial organization, we selected two of our published in-cell XL-MS datasets and combined them for higher XL-MS coverage, then used the combined data to evaluate if there is a bias in spatial consistency between intact-cell and cell-lysate cross-linking.

To test whether the observed consistency reflects genuine spatial organization rather than protein abundance or hub structure, each marker-marker network was randomized while every protein retained its marker annotation. This degree-preserving null rewired the network so that each protein kept the same number of cross-links but to randomly reassigned partners (via repeated edge-swaps), controlling for protein abundance and hub effects. The null distribution was generated from 500 rewired networks per dataset, and the observed consistency was compared against it.

***Results***

The comparison was performed either on the entire network, using all selected organelle and topology markers (Fig. SN1), or on a reduced subset of markers (Fig. SN2). For the subset, only three organelles (mitochondrion, plasma membrane and nucleus) were used as organelle markers, and only one-third of the CYT+NUC topology markers were retained. The former choice provides a cleaner test environment: the mitochondrion and nucleus each contain a large number of proteins, and connections between any two of these three compartments are forbidden. The latter choice was made because the CYT+NUC compartment is far larger than the other topological compartments and would otherwise dominate the topology-marker set; downsampling it makes the five topology compartments more comparable in size and therefore renders the null more stringent.

Together, both the DUC-XL-MS and in-cell XL-MS datasets showed cross-link spatial consistency at very similar levels (differing by <1%), and both lay far above the degree-preserving null at the organelle and topology levels. For the full marker set, the two datasets exceeded the random null by 15%-30%, and for the stringent, size-balanced subset both datasets maintained a similarly high consistency, and 40%-60% above the null. These results indicate that DUC-XL-MS preserves protein spatial organization to a similar extent as in-cell XL-MS, and that the high consistency does not arise from protein abundance or network connectivity but instead reflects genuine subcellular spatial organization retained in the DUC-XL-MS data.

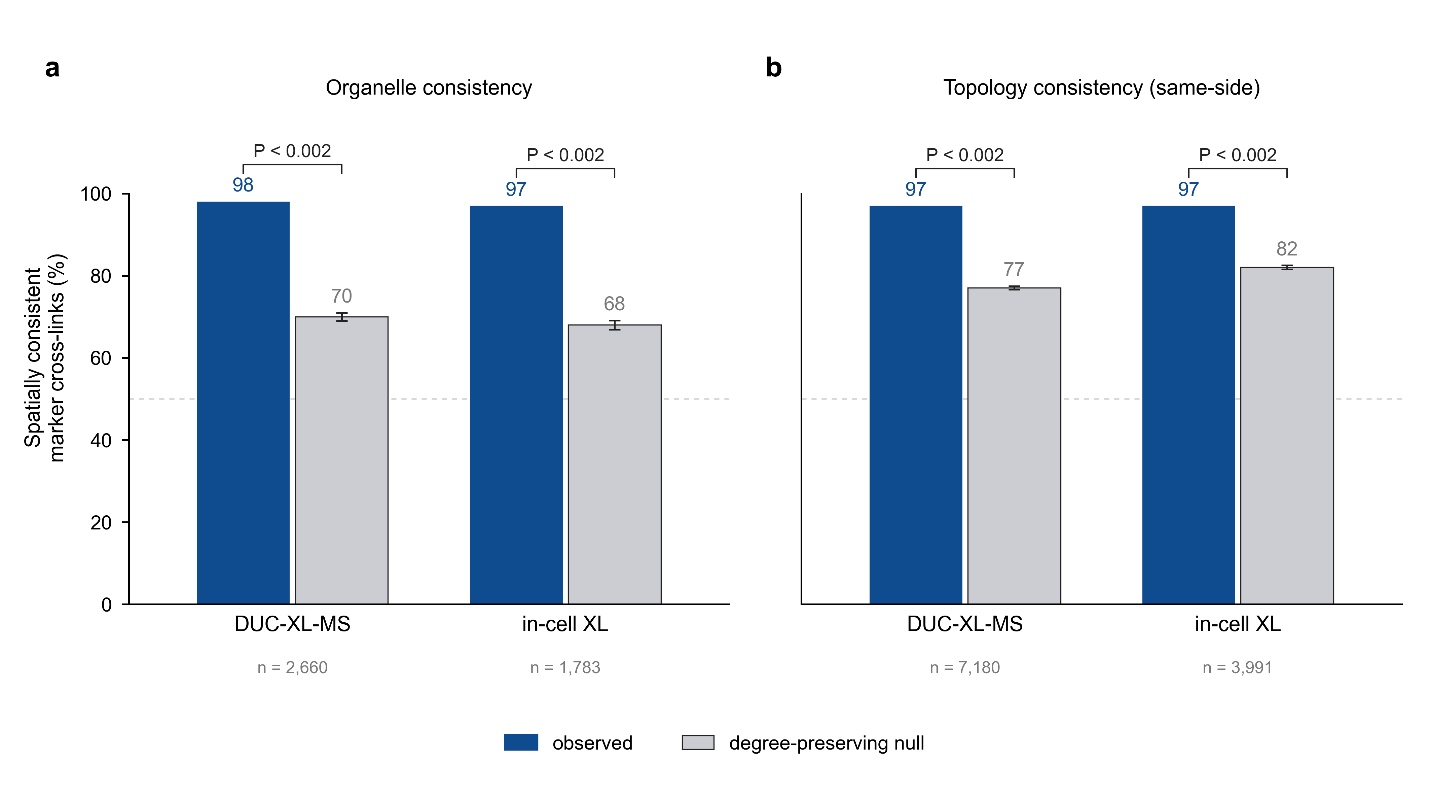

**Fig. SN1 | Cross-link consistency of organelle and topology markers exceeds a degree-preserving random-network null.** (a) Organelle-marker consistency for the networks from DUC-XL-MS and intact HEK293 cells: observed value (blue) versus the degree-preserving null (grey; mean ± s.d. over 500 rewired networks). (b) Topology-marker (same-side) consistency, same design. Bars are annotated with the percentage; brackets give the empirical permutation P. n = marker–marker cross-links (PPIs): organelle 2,660 (DUC) and 1,783 (in-cell); topology 7,180 (DUC) and 3,991 (in-cell). Observed consistency lies far above the null in every case. Source data are provided as a Source Data file.

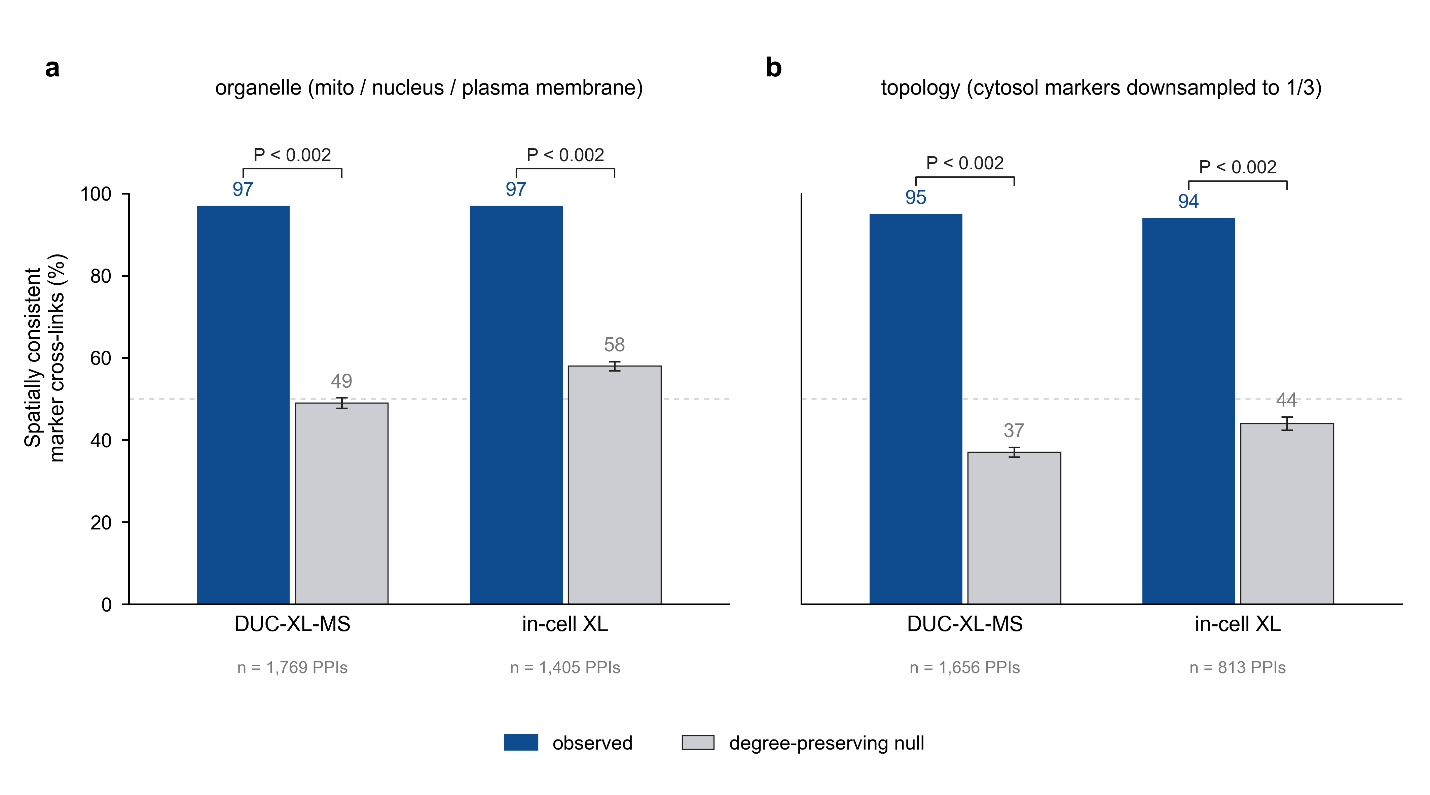

**Fig. SN2 | Consistency remains far above the null on stringent, balanced marker subsets.** (a) Organelle consistency restricted to single markers of three mutually non-adjacent compartments: mitochondrion, nucleus and plasma membrane, for which every inter-compartment cross-link is spatially impossible. (b) Topology consistency after downsampling CYT+NUC markers to one third while retaining all luminal, matrix, intermembrane-space and peroxisomal markers. Bars show observed (blue) versus the degree-preserving null (grey; mean ± s.d., 500 rewired networks); brackets give the empirical permutation P. n = marker–marker cross-links (PPIs): a 1,769 (DUC) and 1,405 (in-cell); b 1,656 (DUC) and 813 (in-cell). Restricting to these subsets drops the null baseline to near coin-flip levels (organelle 49–58 %; topology 37–44 %) while observed consistency holds at 94–97 %, indicating that the spatial signal is not an artefact of marker abundance, high-degree hubs, or the dominant CYT+NUC proteins.

**Supplementary Note 2: The CORVET/HOPS Class C core is structurally conserved from yeast to human**

***Rationale***

Since experimental structures of HOPS and CORVET are available from yeast (PDB 7ZU0 and 8QX8), we asked whether the yeast architecture is a legitimate reference for the human complexes, by comparing the shared Class C core (Vps11, Vps16, Vps18 and Vps33/VPS33A) in sequence and in structure (Fig. SN3).

***Results***

Sequence conservation is low: 21–24% identity between the yeast and human core subunits (Table SN1). Their structures, however, show substantial structural similarity: **TM-align (*1*) yielded 3.27–6.34 Å Cα RMSD over the structurally aligned core regions**, consistently for both structure pairs (Table SN2, SN3). The Class C core thus provides a clear example of structural conservation despite substantial sequence divergence, and the yeast assemblies provide a valid architectural reference for the human complexes.

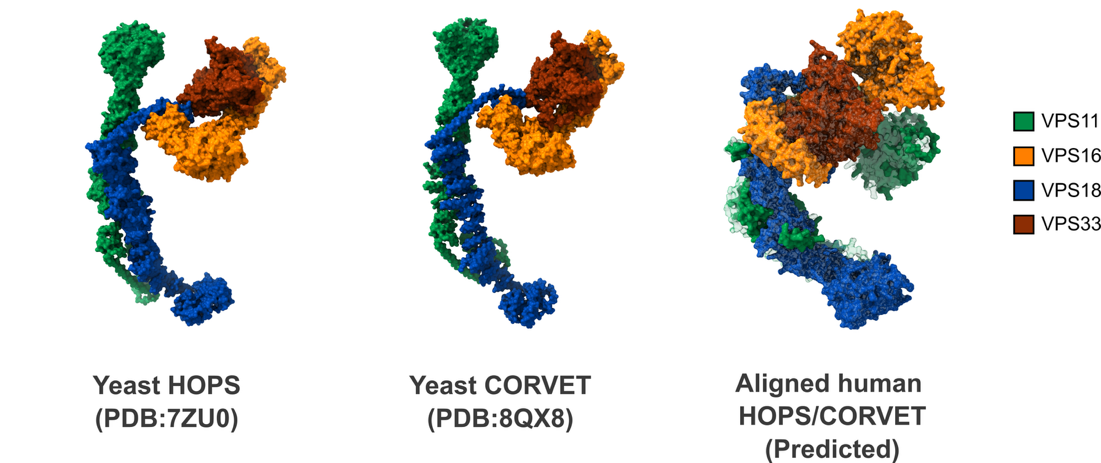

**Fig. SN3 |** **Structural comparison of the homologous Class C core subunits of the HOPS and CORVET complexes between yeast and humans.** The four shared Class C core subunits (Vps11/VPS11, Vps16/VPS16, Vps18/VPS18, and Vps33/VPS33A) are shown for yeast HOPS (left, PDB 7ZU0), yeast CORVET (middle, PDB 8QX8), and the predicted human HOPS/CORVET core models (right). The predicted human HOPS and CORVET core models are highly similar and are superposed, with HOPS shown in opaque and CORVET in semi-transparent representation.

**Table SN1 | The Class C core subunits of yeast and human**

| Subunit pair (yeast/human) | Yeast UniProt | Yeast length (aa) | Human UniProt | Human length (aa) | Sequence identity (%) |
| --- | --- | --- | --- | --- | --- |
| VPS11 | P12868 | 1,029 | Q9H270 | 941 | 21.7 |
| VPS16 | Q03308 | 798 | Q9H269 | 839 | 22.0 |
| VPS18 | P27801 | 918 | Q9P253 | 973 | 21.4 |
| Vps33 / VPS33A | P20795 | 691 | Q96AX1 | 596 | 23.8 |

**Table SN2 | TM alignment of yeast CORVET (PDB 8QX8) core subunits and the human CORVET model**

| Subunit (yeast/human) | Aligned Cα pairs | RMSD (Å) |
| --- | --- | --- |
| Vps11 / VPS11 | 469 | 4.52 |
| Vps16 / VPS16 | 635 | 4.32 |
| Vps18 / VPS18 | 548 | 5.23 |
| Vps33 / VPS33A | 520 | 3.64 |

**Table SN3 | TM alignment of yeast HOPS (PDB 7ZU0) core subunits and the human HOPS model**

| Subunit (yeast/human) | Aligned Cα pairs | RMSD (Å) |
| --- | --- | --- |
| Vps11 / VPS11 | 501 | 5.32 |
| Vps16 / VPS16 | 670 | 4.85 |
| Vps18 / VPS18 | 674 | 6.34 |
| Vps33 / VPS33A | 516 | 3.27 |

**Supplementary Note 3: Cross-link agreement with the structural models**

***Rationale***

Cross-links mapped onto a predicted model can be used to assess its accuracy. We evaluated the models in two steps: first the individual subunits, using intra-protein cross-links and the per-residue confidence (pLDDT), and second the interfaces, using inter-protein cross-links, which test how two subunits are placed relative to one another. We reason that an interface is interpretable only if both subunit models are themselves confidently predicted and well supported by their own intra-protein cross-links.

***Results***

***Individual subunit models***

All intra-protein cross-links were mapped onto the protein models (Fig. SN4). Agreement was high in all four complexes: 44 of 48 cross-links satisfied (91.7%) in CORVET, 39 of 42 (92.9%) in HOPS, 124 of 138 (89.9%) in the exocyst and 100 of 113 (88.5%) in WASH, and 275 of 307 (89.6%) pooled over the twenty distinct proteins. HOPS and CORVET share the Class C core (VPS11, VPS16, VPS18 and VPS33A), onto which each complex adds its specific subunits: VPS39 and VPS41 for HOPS, and TGFBRAP1 (VPS3) and VPS8 for CORVET. The two complexes are therefore drawn in a single panel, with each shared-core subunit shown once.

Sixteen of the twenty proteins agree with their own cross-links at 80% or above. Four fall below: VPS8 at 60.0% (3 of 5), WASHC1 at 75.0% (6 of 8), WASHC4 at 76.2% (16 of 21) and EXOC2 at 77.8% (14 of 18) (Fig. SN4). As the majority of cross-links being satisfied in every case, we therefore excluded no subunit from the inter-protein analysis on the basis of intra-protein agreement.

We next examined per-residue confidence as an independent measure of model quality. WASHC2C is a clear outlier: 83% of its residues and 82% of its cross-linked residues fall below pLDDT 50, against 0 to 37% for every other subunit in the same complex (Fig. SN5). WASHC2C was therefore excluded from the next step.

***Subunit interface models***

We next mapped inter-protein cross-links onto the models, excluding cross-links involving WASHC2C (Fig. SN6). Violations were more frequent between subunits than within them, and in three of the four complexes they concentrate on a single subunit.

- CORVET/HOPS:
  All six violations lie on VPS33A (VPS11-VPS33A, 2 of 2 cross-links violated; VPS18-VPS33A, 2 of 2; VPS33A-VPS8, 2 of 2), giving 29 of 35 satisfied (82.9%), and the shared-core violations recur identically in HOPS (15 of 19, 78.9%). Because VPS33A's own model is supported by 7 of 7 intra-protein cross-links, its violations reflect subunit placement rather than fold accuracy. The complex-specific interfaces of both CORVET (TGFBRAP1-VPS11, TGFBRAP1-VPS18) and HOPS (VPS11-VPS39) were fully satisfied.
- WASH:
  All seven violations lie on interfaces involving WASHC1, which satisfies only 1 of its 8 inter-protein cross-links (WASHC1–WASHC3, 1 of 3; WASHC1–WASHC4, 0 of 3; WASHC1–WASHC5, 0 of 2), while every interface not involving WASHC1 is fully satisfied (WASHC3–WASHC4, 4 of 4; WASHC4–WASHC5, 4 of 4), giving 9 of 16 satisfied (56.2%). WASHC1 agrees with its own intra-protein cross-links at 75.0%, but it is the least confidently modelled of the retained subunits, with a median pLDDT of 55.5 against 69.7 for WASHC3, 91.3 for WASHC5 and 92.1 for WASHC4. WASHC1 is therefore questionable on both counts: as an individual model and in its placement relative to the rest of the complex. We therefore also excluded WASHC1 in the final model.
- Exocyst:
  Violations are spread over 17 of 22 observed interfaces and involve all eight subunits, with per-subunit violation rates (counted over each subunit's intra- and inter-protein cross-link ends) forming a continuous gradient from EXOC8 (45.3%) to EXOC6 (8.1%). This pattern is consistent with the extended, minimally interacting and conformationally flexible architecture of the human exocyst reported in a recent structural study, in which a single C-terminal loop of EXOC4 (410–492) docks into a pocket on EXOC5 and is required for holocomplex formation, its deletion abolishing assembly of the two subcomplexes, while the remainder of the complex is flexible (*2*). Consistent with this, our cross-links at the EXOC4–EXOC5 interface are only 50% satisfied (4 of 8), as expected if a single loop-mediated connection permits substantial relative motion of the two subcomplexes.

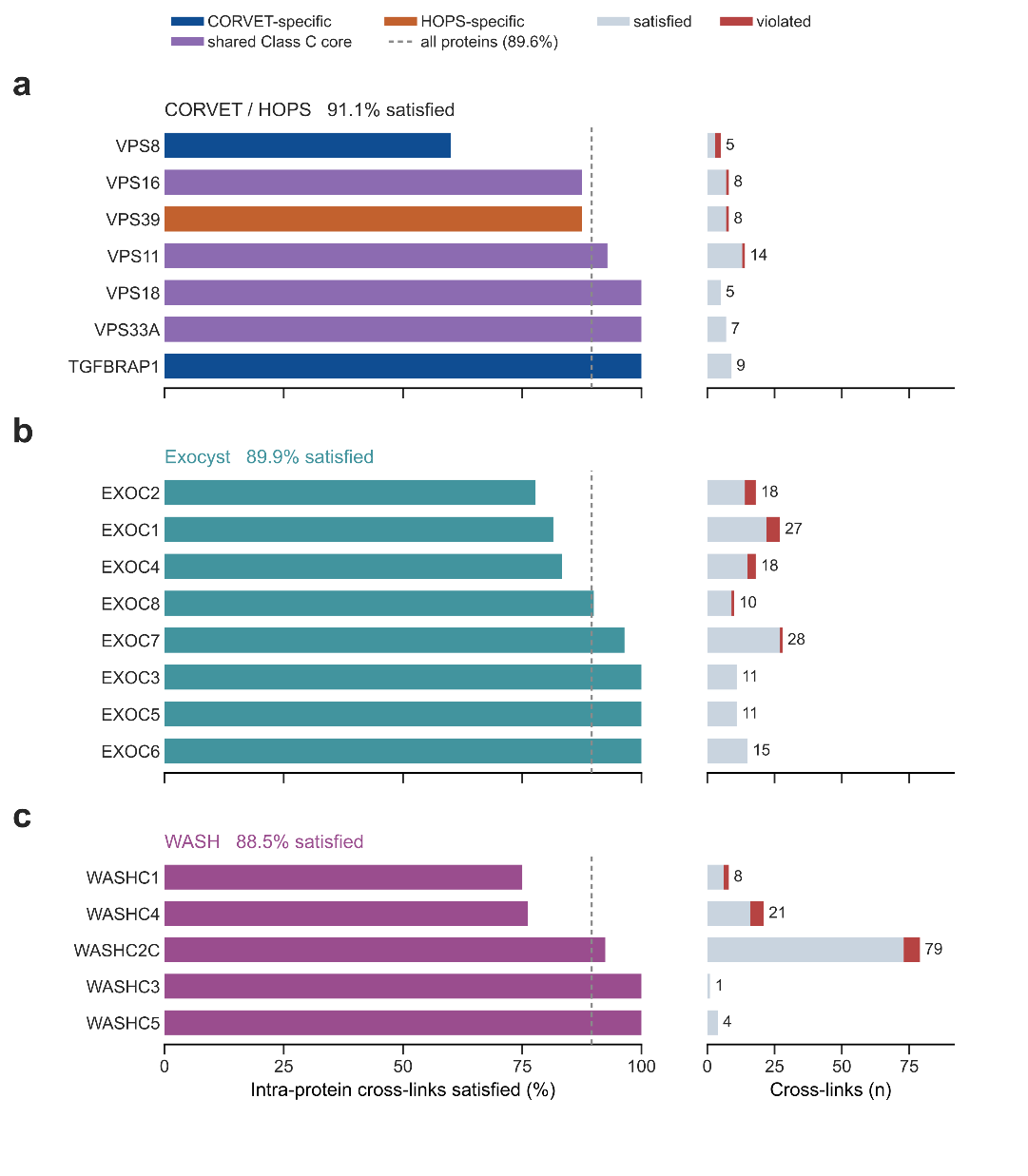

**Fig. SN4 | Intra-protein cross-link agreement per protein, by complex.** a, CORVET and HOPS, with bars colored CORVET-specific, shared Class C core or HOPS-specific. b, Exocyst. c, WASH. In each panel the left column gives the percentage of each protein’s intra-protein cross-links satisfied by its structural model, ordered from lowest to highest; the right column gives the number of cross-links, split into satisfied and violating, with the total at the right of each bar. Dashed line, pooled rate of the twenty distinct proteins (275/307, 89.6% satisfied). The shared Class C core subunits (VPS11, VPS16, VPS18, VPS33A) are drawn once, in the combined CORVET / HOPS panel.

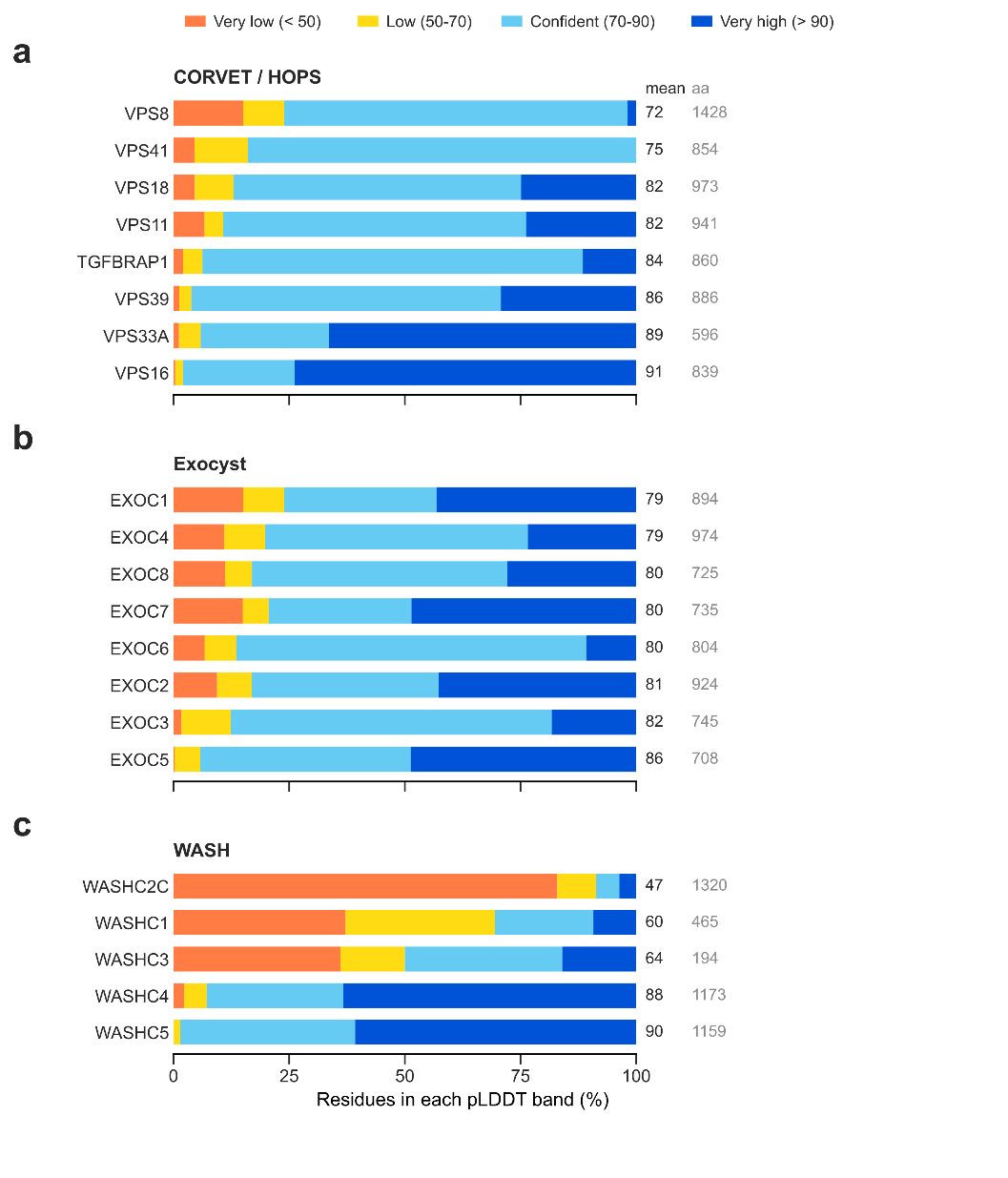

**Fig. SN5 | Model confidence per subunit, by complex.** Fraction of each subunit’s residues in the standard AlphaFold pLDDT bands (very low (< 50), low (50–70), confident (70–90) and very high (> 90)). a, CORVET / HOPS, drawn as one panel with each shared Class C core subunit (VPS11, VPS16, VPS18, VPS33A) shown once alongside the complex-specific subunits (TGFBRAP1 and VPS8 for CORVET; VPS39 and VPS41 for HOPS), b, the exocyst and c, WASH. Within each panel, subunits are ordered by mean pLDDT, lowest at the top; the columns at the right give the mean pLDDT and the protein length in residues. Values are per-chain pLDDT read from the same complex models the cross-links were mapped onto, so confidence and agreement describe the same structure. The shared Class C core subunits are modelled in both CORVET and HOPS; the CORVET copy is shown, the two differing by at most 0.2 in mean pLDDT.

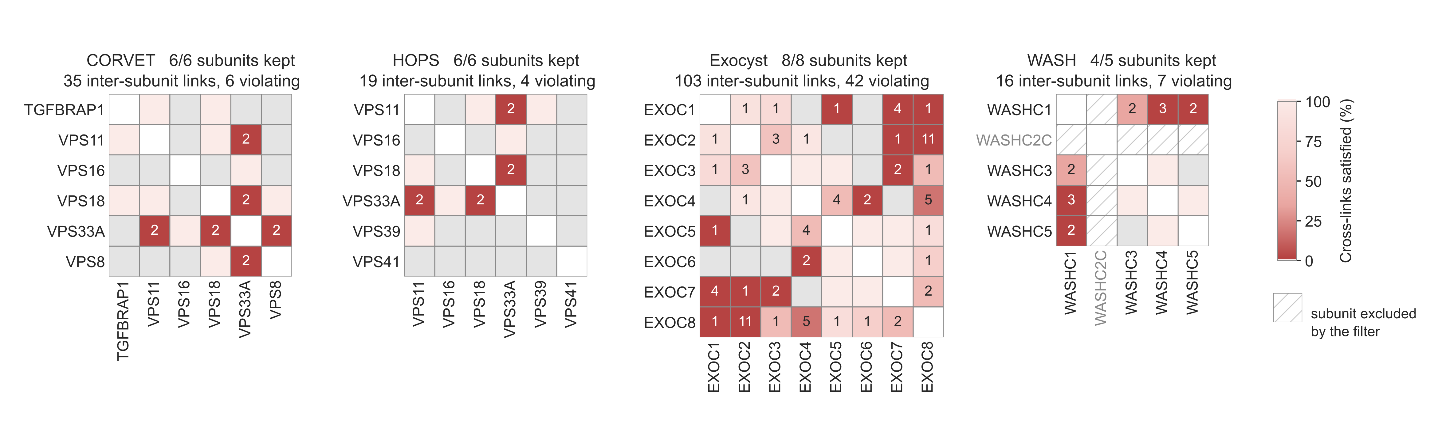

**Fig. SN6 | Inter-protein cross-link agreement per subunit pair, by complex**. Inter-protein cross-link agreement after removing WASHC2C due to its low pLDDT. CORVET and HOPS are shown as separate matrices; their shared Class C core interfaces are therefore drawn in both and are the same measurements. Fill color gives the percentage of cross-links satisfied and the number in a cell the count of violating cross-links, omitted where none were violated. Grey, no cross-link of that protein pair detected. Excluded proteins are retained in position and hatched, with labels in grey. Diagonal cells are blank because intra-protein cross-links define the exclusion and are not part of the map being tested.

**Supplementary Note 4: Consistency of WASH complex cross-links between the HEK293 and Jurkat datasets**

***Rationale***

The WASH complex model was predicted using cross-links from the HEK293 dataset. To test whether this model aligns with XL-MS data from Jurkat cells, we assessed whether the WASH complex cross-links identified in Jurkat cells are also consistent with it.

***Results***

Cross-links from both datasets were mapped onto the predicted WASH complex structure and classified as satisfied or violated at a Cα–Cα distance threshold of 40 Å (Fig. SN7). The two datasets showed the same pattern: intra-molecular agreement was high in both (88.5% in HEK293, 91.0% in Jurkat), and violated inter-molecular cross-links were confined to the same interfaces, those involving WASHC1 and the WASHC2C–WASHC4 pair (Fig. SN7). After removing WASHC1 (owing to its low pLDDT and the high proportion of violated cross-links to other subunits) and WASHC2C (owing to its low pLDDT), all remaining observed interfaces were fully satisfied in both datasets (8 of 8 cross-links each; Fig. SN8). Together, these results show that the predicted WASH complex structure is supported equally well by the HEK293 and Jurkat data.

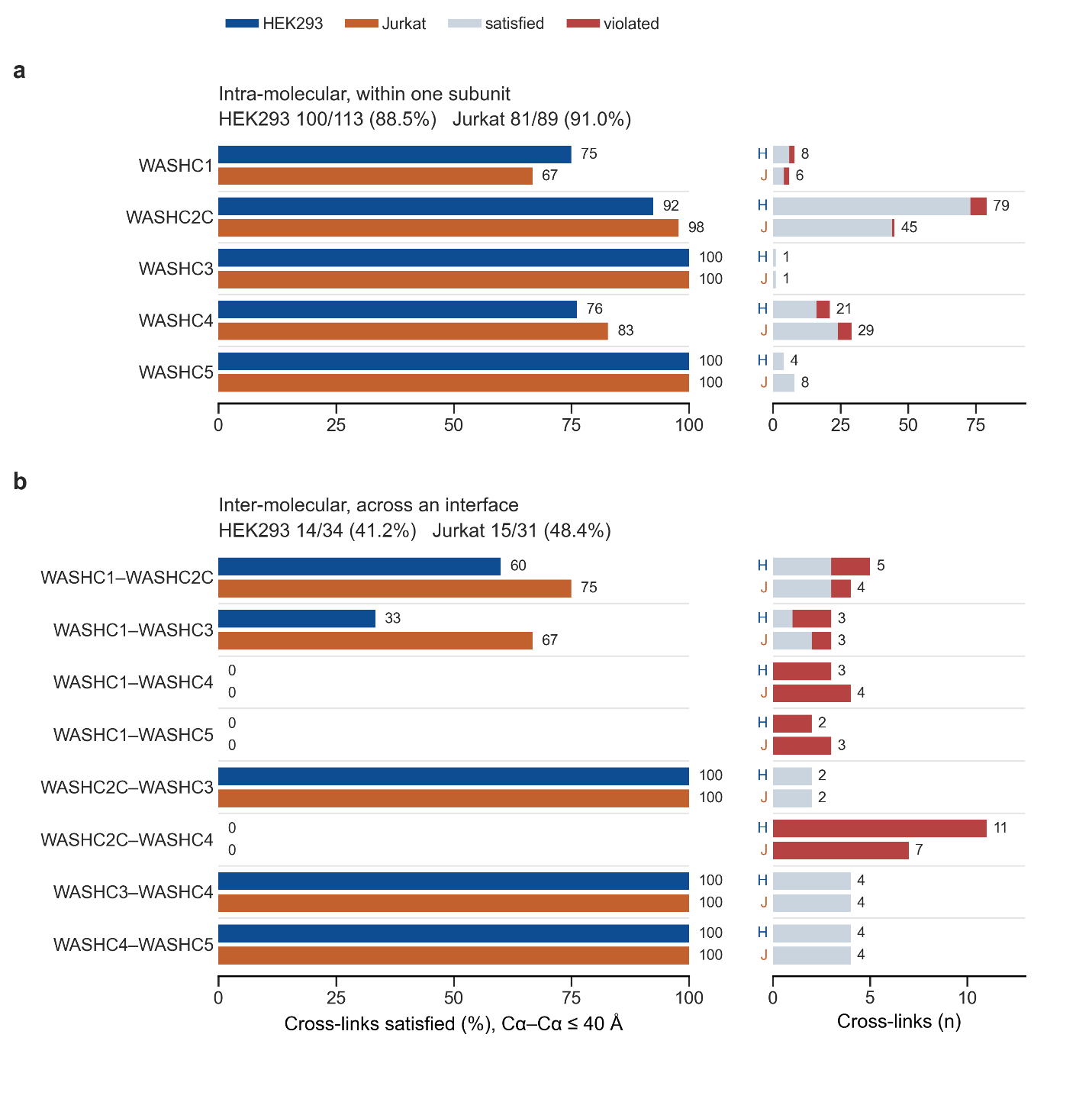

**Fig. SN7 | Cross-link agreement of the WASH complex model in HEK293 and Jurkat cells**. Cross-links from the HEK293 (blue) and Jurkat (orange) DUC-XL-MS datasets were mapped onto the same predicted WASH complex structure and classified as satisfied or violated at a Cα–Cα threshold of 40 Å. a, Intra-molecular cross-links; b, Inter-molecular cross-links. Left, percentage of cross-links satisfied per subunit (a) or subunit pair (b), with values at the bar ends. Right, the underlying cross-link counts, split into satisfied (light gray) and violated (red); labels give the total per bar. Subunit pairs without observed cross-links (WASHC2C–WASHC5, WASHC3–WASHC5) are not shown.

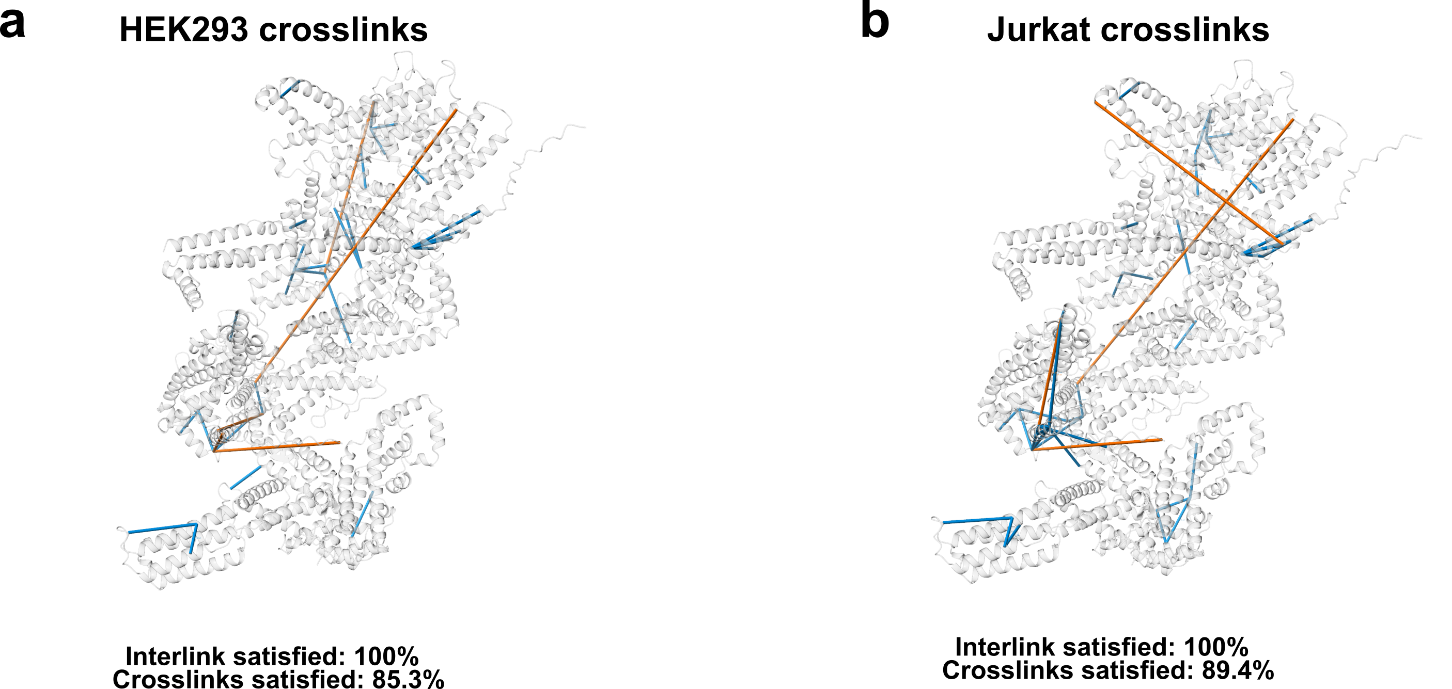

**Fig. SN8 |** **Mapping of XL–MS cross-links onto the predicted WASHC3–WASHC4–WASHC5 structure.** Cross-links identified in the HEK293 (**a**) and Jurkat (**b**) datasets were mapped onto the predicted structure of the WASHC3–WASHC4–WASHC5 subcomplex (WASHC1 and WASHC2C excluded owing to low model confidence). Cross-links satisfying the 40 Å Cα–Cα distance threshold are shown in blue; those exceeding it are shown in orange.

**Supplementary Note 5: GO enrichment of dataset-specific endosomal interactions in DUC-XL-MS and EndoMAP**

***Rationale***

DUC-XL-MS profiles whole cells, whereas EndoMAP builds its network from affinity-enriched endosomes with EEA1 as the bait. We set out to compare how much of the endosomal interactome each dataset covers. For consistency of method, only the EndoMAP cross-linking data (EndoMAP XL) were used, excluding the blue-native co-elution data, so that both datasets were based on the same type of measurement.

***Results***

Endosomal interactions were defined as those in which at least one of the cross-linked proteins carries an endosomal localization. After filtering, the two cross-linking maps are of similar size at the endosome, 371 interactions in DUC-XL-MS and 356 in EndoMAP XL, yet they share only 24 interactions (Fig. SN9A). The overlap between the detected proteins is correspondingly low, with 73 shared proteins (Fig. SN9B). The limited agreement therefore originates from a difference in which proteins each method detects rather than from a disagreement about the interactions themselves.

To ask what distinguishes the two datasets, Gene Ontology enrichment was performed against the union of their endosomal sub-networks. The 31 proteins carrying the 24 shared interactions are strongly enriched for small, stable machineries (Fig. SN10A): terms for vesicle docking and for transport and fusion at the late endosome and lysosome are recovered completely or nearly so, whereas larger terms such as multivesicular body assembly are recovered only partially.

DUC-XL-MS is enriched for earlier, cytosol-facing steps, and its advantage is substantial in absolute numbers (Fig. SN10B): it recovers ten to fifteen more proteins per term for endocytic recycling, macroautophagy, retrograde transport from endosome to Golgi, multivesicular body assembly and membrane fission, in most cases approaching complete coverage of the term. EndoMAP XL leads in the opposite direction, on the late fusion steps at the lysosome and vacuole, but only by one or two proteins in each case.

The two unique sets also differ in where their proteins reside (Fig. SN11). Proteins unique to DUC-XL-MS are more often endosomal, and more often assigned to the endoplasmic reticulum and mitochondrion, whereas proteins unique to EndoMAP XL are more often lysosomal or at the plasma membrane.

Whole-cell and endosome-enriched cross-linking therefore yield complementary views of the endosomal interactome: enriching the organelle before cross-linking sacrifices its transiently associated interactors but concentrates cross-link density on its core machinery. These results indicate that combining methods, or even different sample preparations within one method, can substantially improve the coverage of an organelle's interactome.

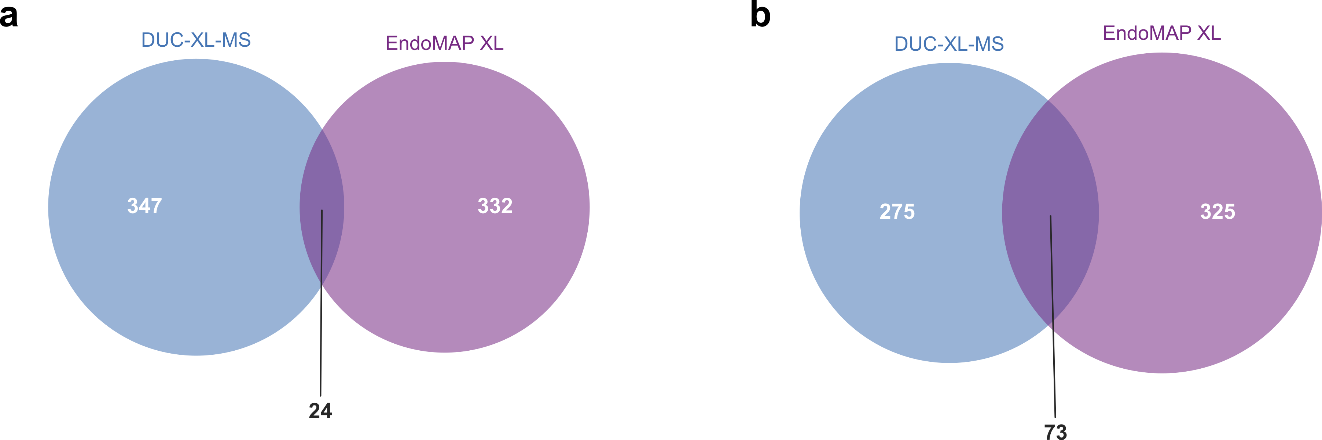

**Fig. SN9 | Overlap of endosomal interactions and proteins between DUC-XL-MS and EndoMAP XL.** A, Endosomal PPIs, defined as those with at least one partner annotated to the endosome, after applying the removal rules described in the Methods. B, The proteins involved in endosomal PPIs. DUC-XL-MS dataset is restricted to HEK293 and EndoMAP dataset is from XL-MS only.

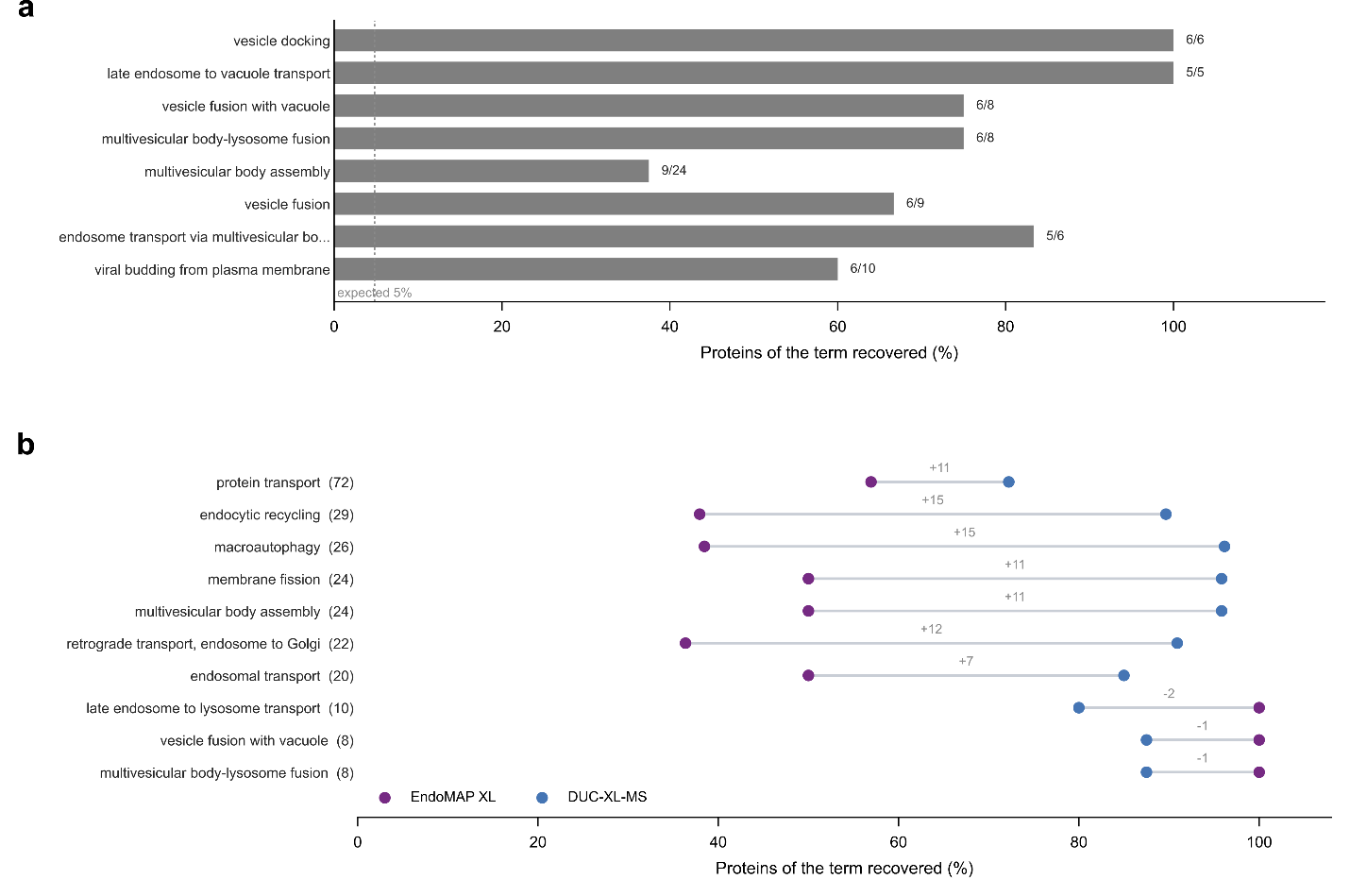

**Fig. SN10 | GO comparison of the DUC-XL-MS and EndoMAP XL datasets.** Both panels are analyzed on the same background, the 673 proteins of the two endosomal sub-networks combined, of which 642 carry a Gene Ontology biological process annotation. A, The eight most significantly over-represented biological process terms among the 31 proteins carrying the interactions found by both datasets, ranked by false discovery rate and tested by the hypergeometric distribution with Benjamini-Hochberg correction. Bar labels give proteins recovered over proteins in the term. B, Recovery of ten terms by each dataset separately, with no statistical test applied; the terms were chosen to span the range of term sizes and to include both directions of difference, and three of them also appear in A. Terms are ordered by size, given in brackets after each name, and each connector is labelled with the difference in proteins between the two datasets, positive where DUC-XL-MS recovers more. Blue, DUC-XL-MS; purple, EndoMAP XL.

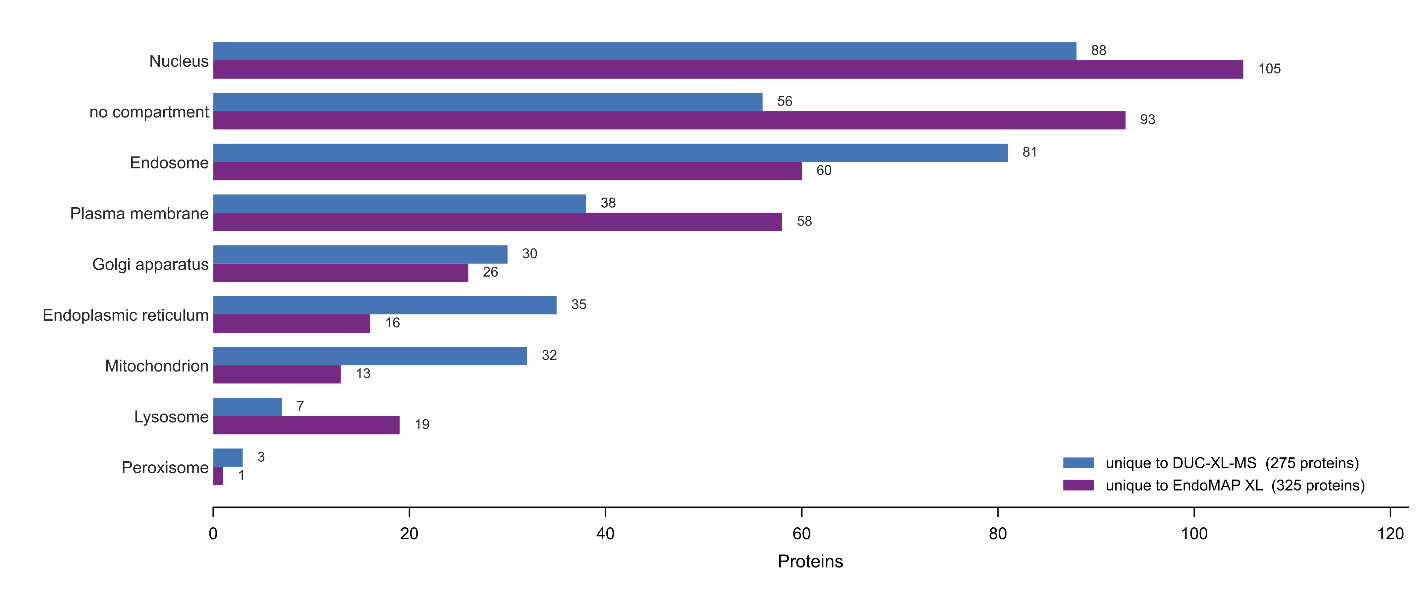

**Fig. SN11 | The proteins unique to each dataset located in different compartments.** Canonical compartments of the proteins carrying the endosomal interactions unique to each sub-network. A protein annotated to several compartments is counted in each, so the bars sum to more than the totals given in the legend, and a protein with no canonical compartment is counted once in the last row.

**Supplementary Note 6: Analysis of structural modeling of COPI**

***Rationale***

Beyond the complex modeling shown in the main text (CORVET, HOPS, exocyst, and WASH), we also modeled the COPI coatomer complex. This model exhibited the lowest agreement with the experimental XL-MS restraints, with only 38.3% of all cross-links and 26.6% of inter-protein cross-links satisfied. In addition, the predicted model contained extensive steric clashes, indicating that the discrepancy could not be explained simply by conformational flexibility but instead reflected a fundamental inconsistency between the cross-linking data and the structural model being reconstructed. We therefore analyzed this case in detail to understand the limitations of our modeling approach.

***Results***

We compared our model with the experimentally determined COPI coatomer structure (PDB 5A1U, Mus musculus coatomer with Saccharomyces cerevisiae Arf1). The individual subunit folds are largely preserved between the experimental and predicted structures (TM-scores 0.48–0.81 (*1, 3*); Table SN4), but the overall assembly architecture differs substantially (Fig. SN12). The discrepancy therefore lies in the arrangement of the subunits rather than in their folds.

Unlike the four complexes reported in the main text, COPI forms a repeating membrane-associated coat in which neighboring coatomer copies are closely packed (Fig. SN13). Cross-links therefore arise both within one coatomer and between adjacent copies, and mass spectrometry cannot distinguish the two. Because our modeling interprets every cross-link as a restraint on a single isolated coatomer, cross-links that span adjacent copies become restraints that no single copy can satisfy (Fig. SN13).

The COPI case thus defines an applicability boundary of our XL-assisted Monte Carlo assembly approach: it is designed for discrete complexes in which cross-links can be assigned to a defined assembly, not for repeating higher-order architectures in which intra- and inter-complex cross-links are experimentally indistinguishable. Modeling such systems will require explicit representation of the repeating architecture and assignment of cross-links to individual copies or asymmetric units, potentially aided by complementary information such as cryo-EM or cryo-ET.

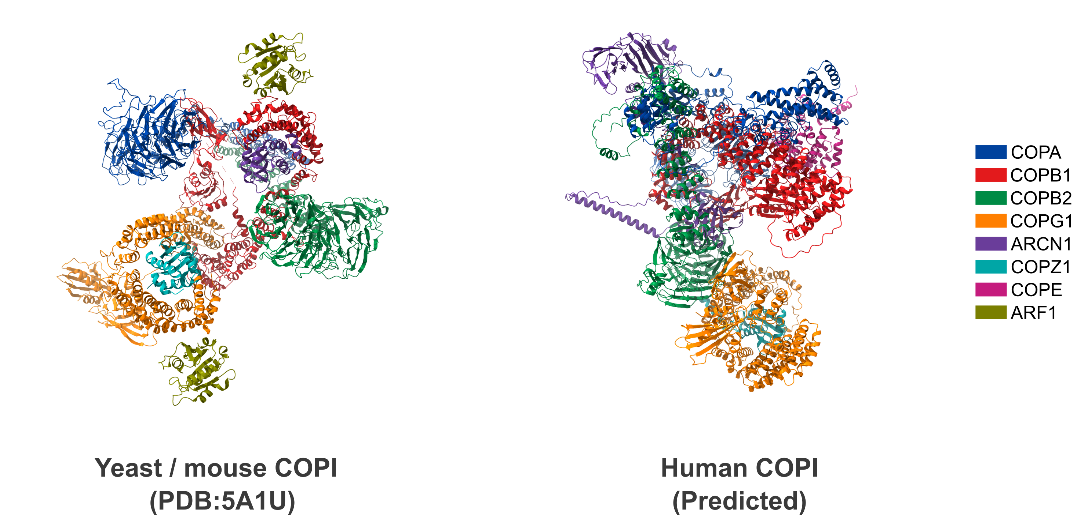

**Fig. SN12 | Comparison of the cryo-EM COPI coat structure with the predicted human COPI model.** Left: COPI coatomer structure (PDB 5A1U), comprising *Mus musculus* coatomer subunits and *Saccharomyces cerevisiae* Arf1. Right: our predicted human COPI model. The predicted model adopts a substantially different global organization and contains severe steric clashes between assembled subunits, with a MolProbity all-atom clash score (*4*) of 74.39.

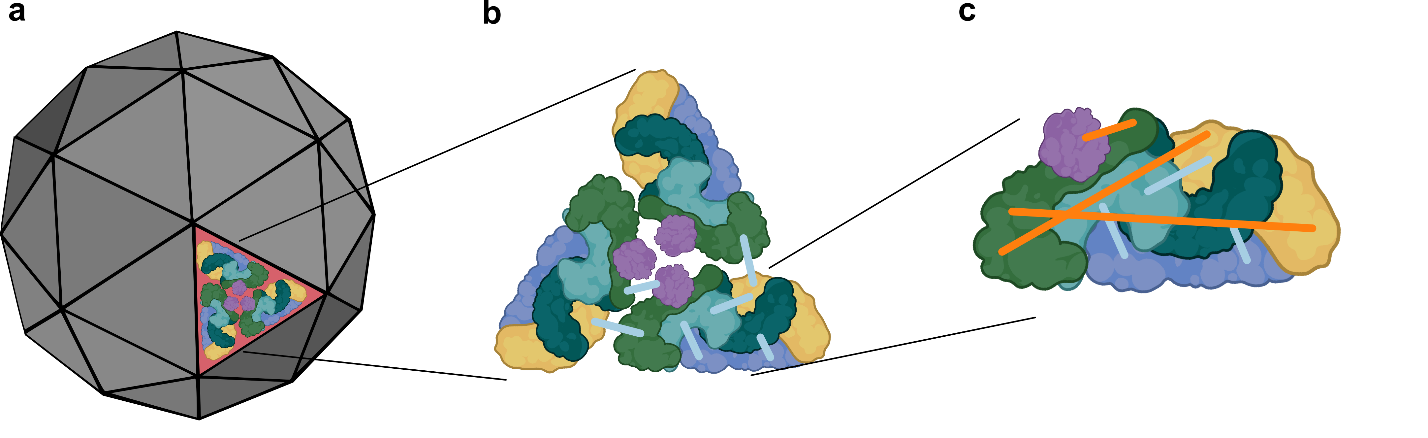

**Fig. SN13 | Cross-link assignment ambiguity in multi-copy complexes.** a, Native COPI assembles into a repeating membrane-associated coat in which neighboring coatomer complexes pack closely together. b, A triad of neighboring coatomer complexes. Intra- and inter-complex cross-links, correctly assigned in their native structural context, are shown in light blue. c, The assignment ambiguity arising during single-complex reconstruction. Inter-complex cross-links originating from neighboring coatomer copies (orange) are experimentally indistinguishable from intra-complex cross-links and are therefore incorrectly imposed on a single coatomer copy. Although these cross-links reflect genuine inter-assembly contacts, their misassignment introduces incompatible spatial restraints, distorting the reconstructed COPI architecture.

**Table SN4 | Structural conservation of individual COPI subunits between the cryo-EM coat structure (PDB 5A1U) and the predicted human COPI model.**

| **Gene** | **5A1U length (aa)** | **COPI core length (aa)** | **Sequence identity (%)** | **Aligned residues** | **TM-score** | **RMSD (Å)*** |
| --- | --- | --- | --- | --- | --- | --- |
| ARCN1 | 135 | 511 | 100.0 | 135 | 0.550 | 2.03 |
| COPA | 813 | 1224 | 99.6 | 813 | 0.483 | 6.12 |
| COPB1 | 813 | 953 | 99.3 | 813 | 0.614 | 6.24 |
| COPB2 | 803 | 906 | 99.3 | 803 | 0.777 | 4.60 |
| COPG1 | 824 | 874 | 97.8 | 824 | 0.596 | 6.19 |
| COPZ1 | 139 | 177 | 100.0 | 139 | 0.806 | 1.73 |

*RMSD denotes the TM-align-optimized Cα RMSD *(1)*.
